# Diversity and divergence of two sympatric, sibling octopus species

**DOI:** 10.64898/2026.04.30.721928

**Authors:** Gabrielle C. Coffing, Silas Tittes, Scott T. Small, Andrew D. Kern

## Abstract

Coleoid cephalopods have convergently evolved many traits shared with vertebrates, including camera-type eyes, large brain-to-body size ratios, and complex behaviors. Most evolutionary studies of cephalopods have compared individual genomes of taxa that diverged tens to hundreds of millions of years ago, yet very few have examined more recent evolution from a population genetics perspective. Here we present a comparative population genomic analysis of the sympatric sister species *Octopus bimaculatus* and *Octopus bimaculoides* using whole-genome resequencing. Despite similar morphologies, these species differ substantially in their life histories, ecologies, and geographic distributions. Using demographic inference, we estimated that the two species diverged approximately one million years ago and that *O. bimaculatus* has maintained a consistently larger effective population size since divergence. Consistent with these demographic histories, we found stronger signatures of positive selection in *O. bimaculatus*, including a positive correlation between recombination rate and nucleotide diversity, more selective sweeps, and a higher proportion of mutations fixed by adaptation—all consistent with more efficient natural selection in larger populations. Protein-coding genes overlapping with selective sweeps were enriched for various functions that included many related to brain and eye development, suggesting that traits characteristic of coleoid cephalopods continue to be shaped by positive selection on recent timescales in these species. Comparing coding-sequence divergence on the Z chromosome to the autosomes, we also find evidence for a female-biased mutation rate, consistent with an independent estimate from a deeper-timescale cephalopod comparison.

## Introduction

Coleoid cephalopods are a diverse group of molluscs that include squids, octopuses, and cuttlefish. They have independently evolved numerous traits shared with vertebrates, including large brains, camera-type eyes, and complex behaviors. This diverse array of derived traits has made them a fascinating group to study across a wide range of biological sub-disciplines, including evolutionary biology, which has long been interested in understanding the nature of convergence and historical contingency (Gould 1989). Research on coleoid cephalopods has yielded a better understanding of much of their unique biology, including the proteins involved in neurogenesis (Albertin et al. 2015), mechanisms of sex determination (Coffing et al. 2025), and single-cell atlases that reveal the cell types and gene expression profiles underlying their visual systems (Songco-Casey et al. 2022; Gavriouchkina et al. 2025). However, most research to date has been motivated by evolutionary questions relevant to the entire clade, representing ancient trait evolution that originated hundreds of millions of years ago. Studies of more recent evolution from a population genetics perspective have been far less common (Bein et al. 2023; Lau et al. 2025; McKeown, Arkhipkin, and Shaw 2019; Cheng et al. 2021), and there is little understanding of the ongoing role of natural selection in any cephalopod species. Taking advantage of recently developed genomic resources, we set out to study natural selection from a population genetics perspective in the Two-spot octopus.

The commonly named Two-spot octopus comprises two sympatric sister species: *Octopus bimaculoides* (Pickford and McConnaughey 1949) and *Octopus bimaculatus* (Verrill 1883). Both species are characterized by blue eye spots, or ocelli, located beneath the eyes near the base of arms two and three (Pickford and McConnaughey 1949; Morris, Abbott, and Haderlie 1980). Due to their nearly identical external appearances, it was not until 1949 that the Two-spot octopus was recognized as two distinct species (Pickford and McConnaughey 1949). Despite their morphological similarity, the sister species are differentiated by their geographic ranges and reproductive strategies (Domínguez-Contreras, Munguia-Vega, Ceballos-Vázquez, Arellano-Martínez, et al. 2018). It has also been suggested that premating barriers to reproduction exist between species (Pickford and McConnaughey 1949). Generally, adults of the two species can be differentiated morphologically by the shape of the ocelli and the number of suckers per arm (Hofmeister and Voss 2024). At sexual maturity, *O. bimaculoides* is generally smaller than *O. bimaculatus* (Pickford and McConnaughey 1949).

Both Two-spot octopus species inhabit rocky reefs, muddy areas, and sandy bottom habitats. The range of *O. bimaculoides* is more northerly, extending from the coast of southern California, U.S.A. to Magdalena Bay on the Baja California peninsula, Mexico (Morris, Abbott, and Haderlie 1980; Domínguez-Contreras, Munguia-Vega, Ceballos-Vázquez, Arellano-Martínez, et al. 2018; Alejo-Plata et al. 2014). In contrast, the northern limit of the *O. bimaculatus* range is in southern California, with the range extending further south into the northern half of the Gulf of California (Lang and Hochberg 1997; Domínguez-Contreras, Munguia-Vega, Ceballos-Vázquez, Arellano-Martínez, et al. 2018; R. F. Ambrose 1997) (fig. 1A). The two species may occupy slightly different ecological niches: *O. bimaculoides* is frequently found in the high intertidal *Laminaria* zone, where seaweed beds begin (Pickford and McConnaughey 1949). These zones contain rocky tidal flats, coastal lagoons, shallow bays, and mudflat areas that are often exposed at low tide (Markaida and Castellanos-Martínez 2024). *O. bimaculatus* is typically found in deeper parts of the intertidal and subtidal communities, below the *Laminaria* zone (Pickford and McConnaughey 1949), and is often caught near California’s Channel Islands (Hofmeister and Voss 2024; R. Ambrose 1988; Hofmeister and Voss 2017). *O. bimaculoides* has been found as deep as 20 m and *O. bimaculatus* as deep as 50 m (Hofmeister and Voss 2024; Markaida and Castellanos-Martínez 2024).

**Figure 1:**
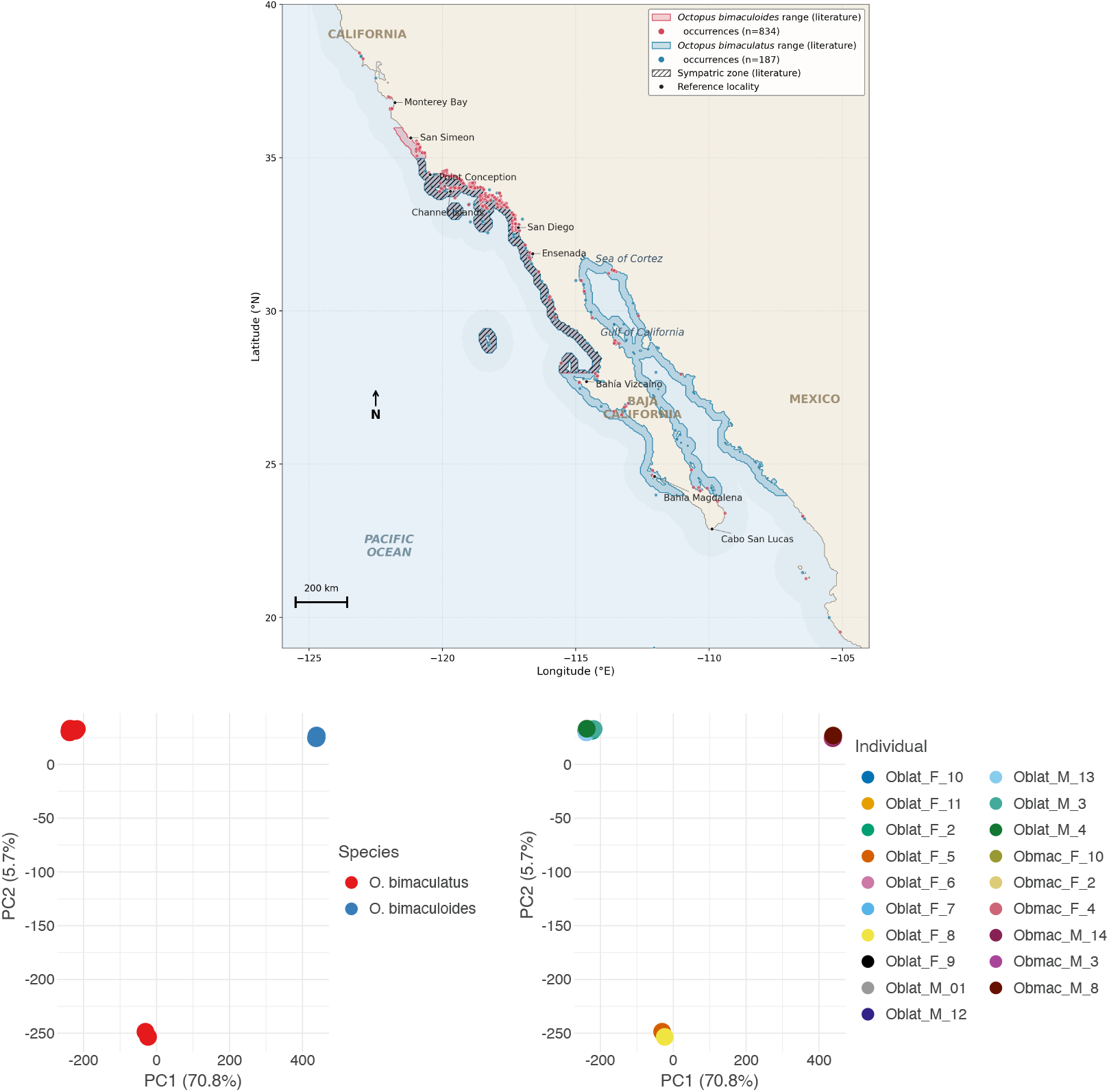
(A) Approximate geographic ranges of *O. bimaculoides* (pink) and *O. bimaculatus* (blue) along the northeastern Pacific coast from published data. (Pickford and McConnaughey 1949; Voight 1988; Hofmeister and Voss 2017; Domínguez-Contreras, Munguia-Vega, Ceballos-Vázquez, Granados-Amores, et al. 2024; Jereb et al. 2014). *O. bimaculoides* extends along the Pacific coast from California south to the western Baja California peninsula and is absent from the Gulf of California; *O. bimaculatus* extends from Point Conception, California, south to Cabo San Lucas, Baja California Sur, and throughout the Gulf of California. Polygons are restricted to nearshore cells within ∼ 30 km of the coast. Filled points are GBIF and OBIS occurrence records (*O. bimaculoides*: *n* = 834; *O. bimaculatus*: *n* = 187). The hatched region marks the zone of sympatry; labeled points mark reference localities. (B) Principal component analysis of chromosome 1 for *O. bimaculoides* and *O. bimaculatus* individuals. Sub-panels are colored by species and by individual ID.

The sister species also differ in their reproductive seasons, fecundity, and egg characteristics. *O. bimaculoides* lays relatively small festoons, or chain-like structures, of large eggs that develop directly into juvenile benthic hatchlings. In contrast, *O. bimaculatus* lays clutches of over 20,000 small eggs that undergo a planktonic paralarval drift stage, allowing individuals to disperse far from their hatching sites over a period of up to three months (Domínguez-Contreras, Munguia-Vega, Ceballos-Vazquez, Arellano-Martínez, et al. 2018; Villanueva et al. 2016). In both species, the timing of the reproductive season depends on geographic location; in shared parts of their ranges, their reproductive periods are generally non-overlapping (Domínguez-Contreras, Munguia-Vega, Ceballos-Vázquez, Arellano-Martínez, et al. 2018) (Table S2).

As marine stocks are continuously fished at unsustainable levels around the world, humans are predicted to become more reliant on resources at lower trophic levels, including cephalopods (Pauly et al. 2002; Sala et al. 2004). The global cephalopod trade has been increasing despite inadequate management practices; in 2022, cephalopods made up 7% of the world’s total exports of aquatic animal products (FAO 2024). There is no commercial fishing of the Two-spot octopus in the U.S.; however, many small-scale fishers in Northwest Mexico sell octopus catch locally and in commercial markets (Domínguez-Contreras, Munguia-Vega, Ceballos-Vázquez, Arellano-Martínez, et al. 2018). The difficulty of classifying octopus catch to the species level prevents official reports from including this information in their fishery statistics (Domínguez-Contreras, Munguia-Vega, Ceballos-Vazquez, Arellano-Martínez, et al. 2018). Without species-level data, it is impossible to know whether a particular population is being overexploited. Given the different life history strategies of the Two-spot octopus sister species, they likely require distinct management strategies.

A previous study examined genetic diversity and population structure using 16S and COI sequences of *O. bimaculoides* and *O. bimaculatus* individuals from Northwest Mexico, finding that *O. bimaculatus* has higher levels of gene flow and a larger effective population size than *O. bimaculoides* (Domínguez-Contreras, Munguia-Vega, Ceballos-Vazquez, Arellano-Martínez, et al. 2018). Here we expand on these findings by investigating population structure, diversity and divergence, demographic history, and evidence of selection in Two-spot octopus individuals from the California coast using whole-genome resequencing. Our analyses reveal two species with sharply contrasting evolutionary histories: *O. bimaculatus* maintains a large, stable effective population size with signatures of linked selection, adaptive protein evolution, and selective sweeps, whereas *O. bimaculoides* has experienced a sustained demographic decline that has left little detectable imprint of positive selection. Z-chromosome patterns further support a sex-biased mutation rate and an achiasmate meiotic regime in the heterogametic sex, consistent with Haldane–Huxley expectations for this clade.

## Results

### Population structure

We generated whole-genome resequencing data from 19 Two-spot octopus individuals (6 *O. bimaculoides* and 13 *O. bimaculatus*) collected along the coast of southern California between 2019 and 2022 (Supplementary Table S1). To our knowledge, this represents the first whole-genome resequencing dataset for any cephalopod analyzed in a population genetics framework. Samples were sequenced on two Illumina NovaSeq 6000 S4 lanes and reads were mapped to the 30-chromosome *O. bimaculoides* reference genome assembly (Coffing et al. 2025). We note that the sample size for *O. bimaculoides* (*n* = 6) is modest and may limit power for some downstream analyses; we address this caveat where relevant below. After filtering for quality, depth, and repetitive content, we retained 1.77 million SNPs in the *O. bimaculoides* VCF and 14.59 million SNPs in the *O. bimaculatus* VCF—an approximately 8-fold difference that reflects the substantially higher nucleotide diversity in *O. bimaculatus* (see below). We additionally generated a merged VCF containing 14.52 million SNPs across both species, which was used for between-species analyses including PCA and demographic inference.

Examination of read depth across chromosomes revealed that two *O. bimaculatus* individuals originally labeled as female (Oblat F 7 and Oblat F 8) did not show the expected halved coverage on the Z chromosome, leading us to reclassify them as male (Supplementary Table S1). This depth-based sex assignment is consistent with the ZZ/ZO sex determination system recently characterized in *O. bimaculoides* (Coffing et al. 2025).

Principal component analysis (PCA) reveals a clear separation of the two species (fig. 1). On chromosome 1, PC1 and PC2 account for 70.8% and 5.7% of total variation, respectively, and PCA plots for all other chromosomes show similar patterns (Supplementary Figs. S1– S4). Along PC2, two *O. bimaculatus* individuals—Oblat F 5 and Oblat F 8—separate from the remaining *O. bimaculatus* samples, possibly reflecting geographic structure or relatedness among the sampled individuals. For downstream analyses, we chose to retain all samples as the overall patterns of summary statistics and inference were not impacted by their exclusion (results not shown).

### Within-species diversity and between-species divergence

We calculated nucleotide diversity (*π*), Tajima’s *D, d*_*XY*_, and Weir and Cockerham *F*_*ST*_ in non-overlapping 1 Mb windows across all 30 chromosomes for both species, applying the accessibility mask generated during variant filtering (see Materials and Methods). In total, 38.92% and 34.08% of the total genome was accessible in *O. bimaculoides* and *O. bimaculatus*, respectively. Per-chromosome summary statistics for each individual are shown in Supplementary Figs. S5–S64.

Genome-wide, nucleotide diversity (*π*) was approximately 3.7-fold higher in *O. bimaculatus* than in *O. bimaculoides* (Mann-Whitney U test, *p <* 0.001; fig. 2; Table 1). This difference is consistent with the 8-fold difference in SNP counts between species and likely reflects the larger effective population size of *O. bimaculatus* (see Demographic history, below). The distribution of Tajima’s *D* also differed significantly between the two species (Mann-Whitney U test, *p <* 0.001): *O. bimaculoides* showed a slightly positive genome-wide average, consistent with a recent population contraction, while *O. bimaculatus* had a slightly negative average, consistent with a stable or mildly expanding population (Table 1). These patterns were broadly consistent across chromosomes (fig. 2; Supplementary Figs. S5–S64).

**Table 1:**
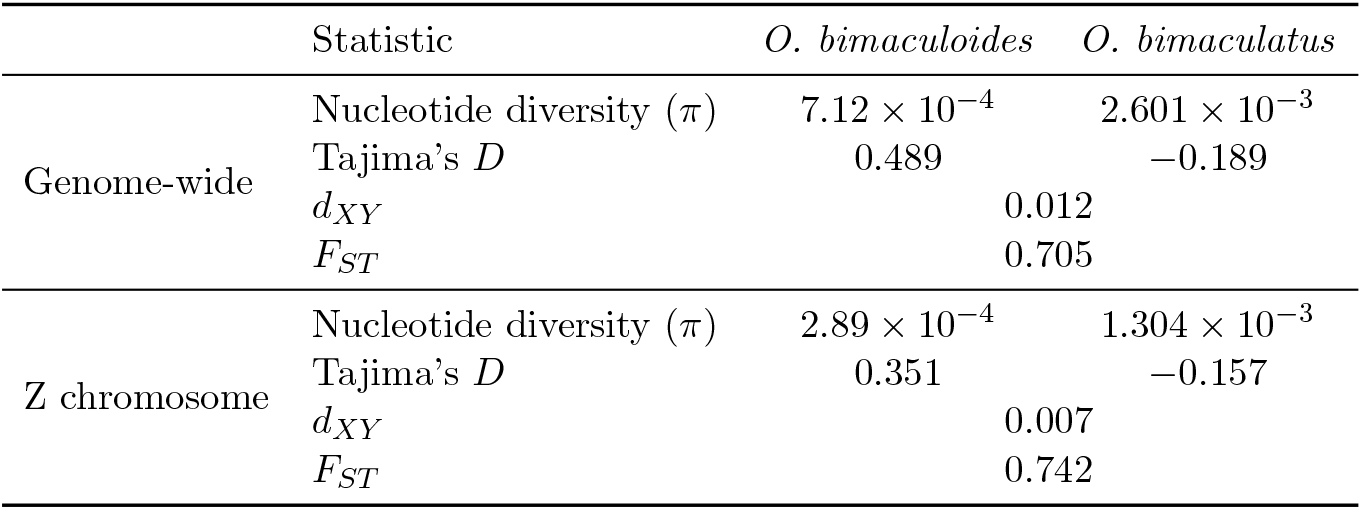
Summary of genome-wide diversity and divergence statistics for the Two-spot octopus species. Values are means across non-overlapping 1 Mb windows.

**Figure 2:**
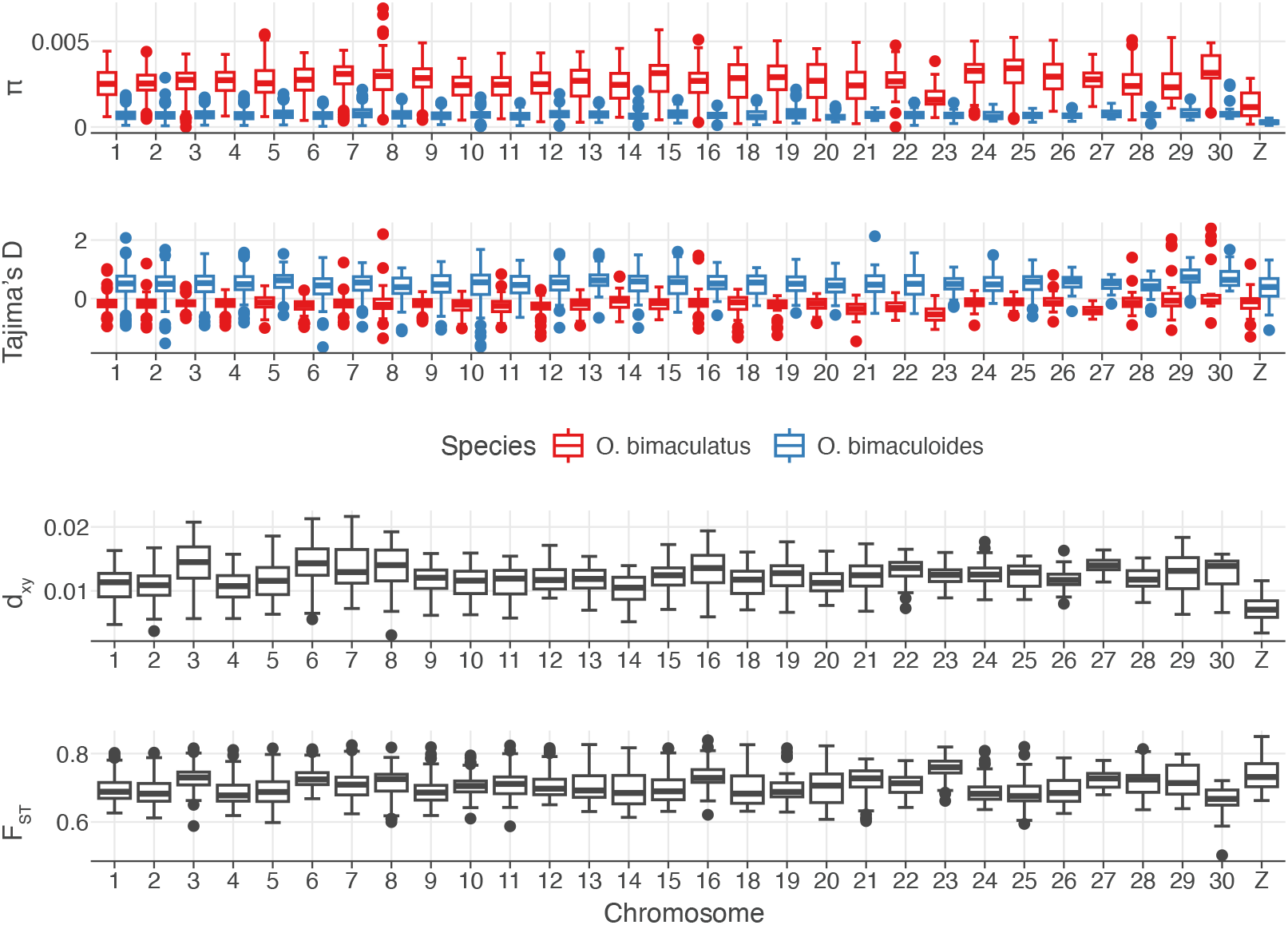
Diversity and divergence statistics calculated for *O. bimaculoides* and *O. bimaculatus*.

We estimated between-species divergence using *d*_*XY*_, finding a genome-wide average of 0.012. This level of absolute divergence is comparable to that observed between humans and chimpanzees (*d*_*XY*_ ≈ 0.01; (Rodrigues, Kern, and Ralph 2024)), consistent with a relatively recent split. In contrast, *F*_*ST*_ between the two species was high (genome-wide average = 0.705), though this relative measure is expected to be inflated when within-population diversity is low, as is the case for *O. bimaculoides* (Cruickshank and Hahn 2014; Charlesworth 1998). The juxtaposition of moderate *d*_*XY*_ and high *F*_*ST*_ therefore reflects the low diversity within *O. bimaculoides* rather than exceptionally high differentiation. Per-chromosome *d*_*XY*_ and *F*_*ST*_ estimates are shown in Supplementary Figs. S65–S74.

On the Z chromosome specifically, nucleotide diversity was reduced roughly 2-fold relative to the autosomal average in both species (*π*_*Z*_ = 2.89 *×* 10^−4^ in *O. bimaculoides* and 1.304 *×* 10^−3^ in *O. bimaculatus*; Table 1), and these Z-vs-autosome differences were significant in both species (Mann-Whitney U test, *p <* 0.001). Between-species divergence on the Z was also lower than the autosomal average (*d*_*XY*_ = 0.007 vs. 0.012; *p <* 0.001), while *F*_*ST*_ was higher (*F*_*ST*_ = 0.742 vs. 0.705; *p <* 0.001; fig. 2). Tajima’s *D* on the Z followed the same species-specific patterns as the autosomes, with a positive value in *O. bimaculoides* (0.351) and a slightly negative value in *O. bimaculatus* (− 0.157). The reduced diversity and absolute divergence on the Z are consistent with its smaller effective population size (three-quarters that of the autosomes under a ZZ/ZO system) and potentially stronger effects of linked selection on sex chromosomes. The elevated *F*_*ST*_ on the Z mirrors the pattern reported in Coffing et al. (2025), where divergence on the Z chromosome was depressed relative to autosomes in a deeper timescale comparison between *O. bimaculoides* and *O. sinensis*, and is consistent with the Fast-Z effect observed in other taxa with ZW/ZO sex determination (Charlesworth, Coyne, and Barton 1987; Mank, Axelsson, and Ellegren 2007).

### Octopus demographic history

We used `SMC++` (Terhorst, Kamm, and Song 2017) to jointly estimate the history of effective population size (*N*_*e*_) and the divergence time of the two species. We ran `SMC++` independently on each of the 30 chromosomes using the merged VCF (containing variants called in both species) and accessibility mask, assuming a per-generation mutation rate of *µ* = 2.4 *×* 10^−9^ as estimated for the Southern blue-ringed octopus (*Hapalochlaena maculosa*; (Whitelaw et al. 2022)), the closest relative for which a mutation rate is available. `SMC++` implements a “clean split” model that assumes no gene flow after divergence; we report the median and interquartile range across the 30 chromosome-level estimates.

The two species show markedly different population size trajectories (fig. 3). *O. bimaculatus* has maintained a large and relatively stable *N*_*e*_ throughout its inferred history. In contrast, *O. bimaculoides* experienced a modest population expansion peaking around 63,000 generations ago, followed by a sustained decline to its present, substantially smaller *N*_*e*_. These trajectories are concordant with the site frequency spectrum summaries reported above (Table 1): the slightly positive genome-wide Tajima’s *D* in *O. bimaculoides* is a hallmark of recent population contraction, while the near-zero *D* in *O. bimaculatus* is consistent with demographic stability.

**Figure 3:**
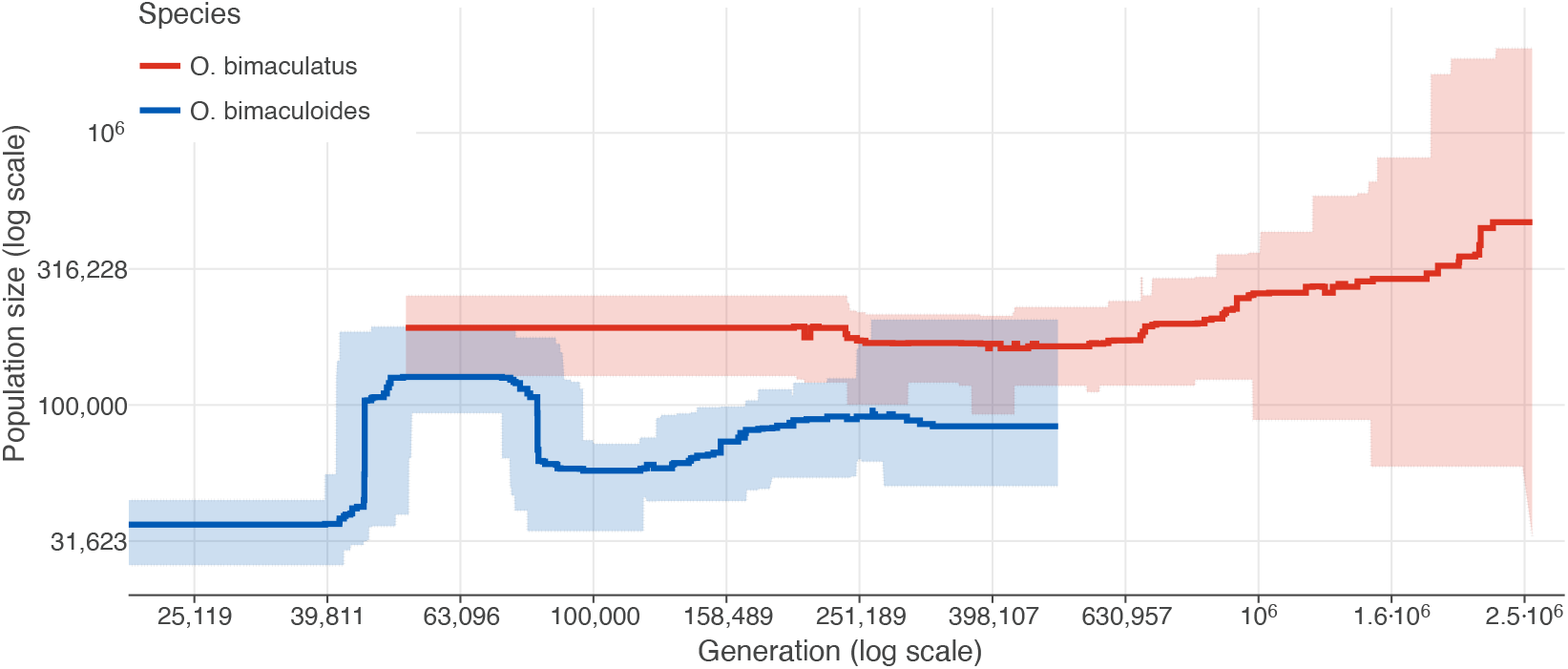
Demographic history estimations generated with `SMC++` for *O. bimaculoides* and *O. bimaculatus*. The dark blue and dark red lines are the median demographic history estimations from all 30 chromosomes. Upper and lower quartiles estimated from the 30 chromosomes are represented by the light blue and light red ribbons.

`SMC++` estimates the species split at approximately 250,000–400,000 generations ago (fig. 3), with uncertainty across chromosomes reflected in the overlapping confidence bands. Both species are semelparous with lifespans of 1–2 years (Table S2); assuming a generation time of approximately 1.5 years, this corresponds to a divergence on the order of 375,000– 600,000 years ago. This `SMC++`-based estimate is considerably more recent than the rough *d*_*XY*_ -based calculation of ∼ 2.5 million generations (*d*_*XY*_ */*2*µ*), which is expected because *d*_*XY*_ reflects coalescence times that predate the population split. The relatively recent divergence is consistent with the moderate levels of *d*_*XY*_ and the near-identical morphology of the two species.

The demographic asymmetry between species—large, stable *N*_*e*_ in *O. bimaculatus* versus smaller, declining *N*_*e*_ in *O. bimaculoides*—has important implications for the efficacy of natural selection, which we explore in the following sections. Theory predicts that selection is more efficient in larger populations, where the fate of beneficial and deleterious mutations is less dominated by genetic drift.

### Recombination rate and patterns of linked selection

We used `ReLERNN` (Adrion, Galloway, and Kern 2020) to estimate local recombination rates along the genome separately for each species, predicting rates in 12 Mb windows using the same accessibility mask applied to our other analyses. Genome-wide mean recombination rates are similar between species: 3.92 *×* 10^−9^ per bp per generation in *O. bimaculoides* and 3.78 *×* 10^−9^ in *O. bimaculatus*, although the distribution of window-level estimates is significantly higher in *O. bimaculoides* (Mann-Whitney U test, *p* = 0.0032; fig. 4A). The per-chromosome recombination landscapes are shown in Supplementary Fig. S75. On most chromosomes, both species show broadly concordant patterns of rate variation along the chromosome, with elevated recombination toward chromosome tips and reduced rate estimates in the central regions on many of the larger chromosomes. This pattern would be consistent with metacentric chromosomes, where the centromere is located near the middle of the chromosome, leading to reduced recombination in the central regions and elevated recombination toward the chromosome ends.

**Figure 4:**
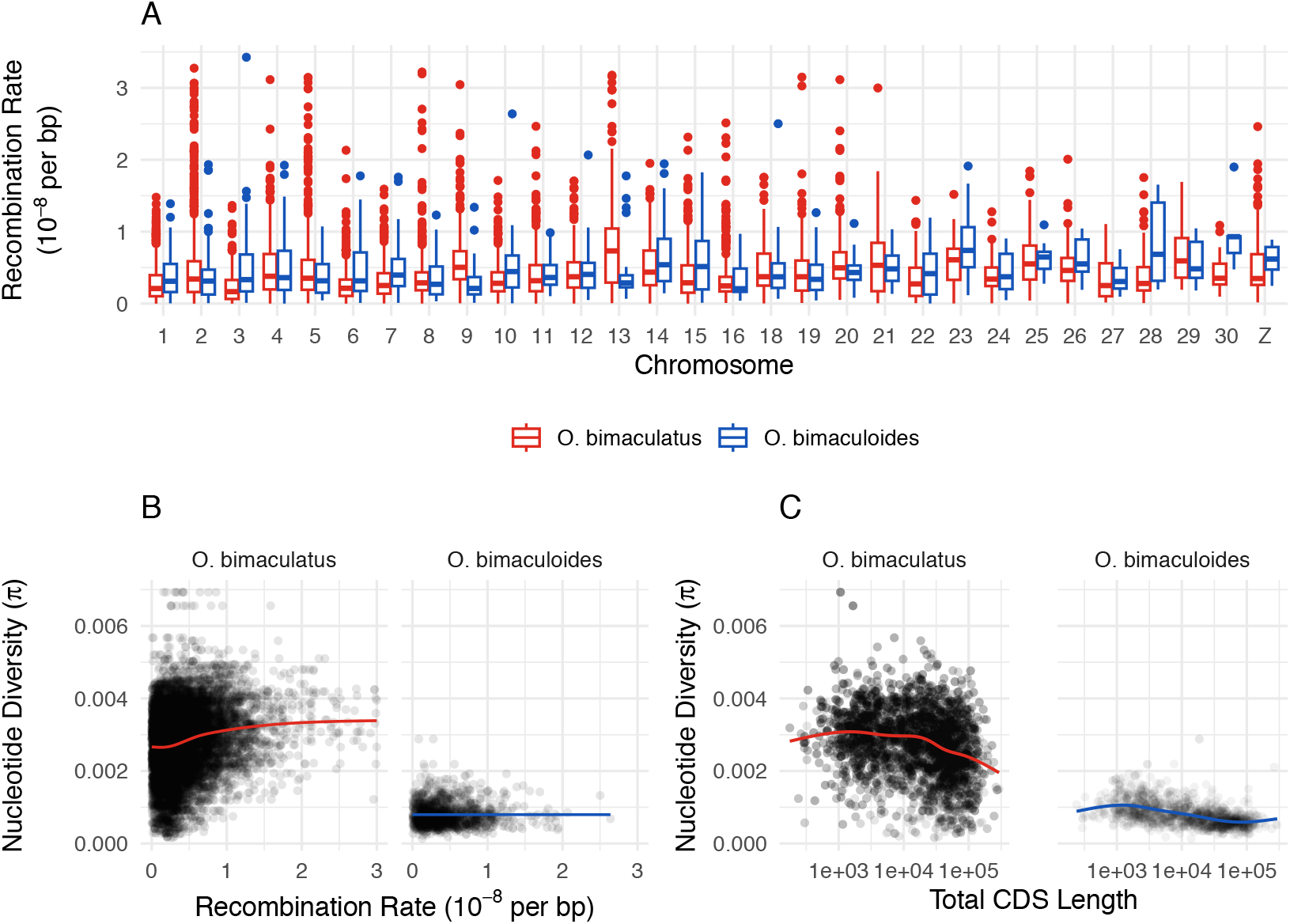
Recombination and linked selection for *O. bimaculoides* and *O. bimaculatus*. (A) Recombination rate variation across chromosomes for *O. bimaculoides* (blue) and *O. bimaculatus* (red). (B) The relationship between nucleotide diversity (*π*) and recombination rate for *O. bimaculoides* (blue) and *O. bimaculatus* (red). (C) The relationship between nucleotide diversity (*π*) and density of coding sequences for *O. bimaculoides* (blue) and *O. bimaculatus* (red). Loess trend lines were added to both panels B and C.

Notably, estimated recombination rates on the Z chromosome are not obviously reduced relative to the autosomes in either species (fig. 4A; Supplementary Fig. S75). Because the Z has a smaller effective population size and recombines only in males, the expected effective recombination rate on the Z depends on assumptions about sex-specific recombination (see Discussion).

Correlations between rates of recombination and levels of nucleotide diversity have long been used as a signature of linked selection—the reduction of neutral diversity by nearby selected variants, whether through selective sweeps or background selection (Begun and Aquadro 1992; Corbett-Detig, Hartl, and Sackton 2015). We observe a striking difference between the Two-spot octopus species in this regard (fig. 4B). In *O. bimaculatus*, there is a clear positive relationship between local recombination rate and *π*: genomic windows with higher recombination rates harbor more nucleotide diversity, as expected when linked selection is a dominant force shaping genome-wide variation. In *O. bimaculoides*, this relationship is essentially absent (fig. 4B). The lack of a diversity–recombination correlation in *O. bimaculoides* is likely a consequence of its smaller *N*_*e*_, in which genetic drift overwhelms the signal of linked selection (Corbett-Detig, Hartl, and Sackton 2015; Buffalo 2021; Lewontin 1974).

We also examined the relationship between nucleotide diversity and the density of coding sequences (total CDS length) in the same 1 Mb windows (fig. 4C). Under linked selection models, regions with more coding sequences—and thus more targets of selection—are expected to show reduced neutral diversity. Both species show a negative relationship between *π* and CDS density, though the pattern is more pronounced in *O. bimaculatus*. We note, however, that the dynamic range of *π* in *O. bimaculoides* is much narrower due to its globally lower diversity, limiting the power of this comparison. Together, the recombination and CDS density results indicate that linked selection has had a stronger genome-wide impact on *O. bimaculatus*, consistent with theoretical expectations for a species with a larger effective population size.

### Divergence in coding regions

To characterize sequence divergence in protein-coding regions between the two species, we estimated per-gene nonsynonymous (*d*_*N*_) and synonymous (*d*_*S*_) divergence across 12,827 single-copy orthologs. Across all orthologs, mean *d*_*N*_ = 0.0056 (median 0.0017) and mean *d*_*S*_ = 0.0100 (median 0.0000) are consistent with the low genome-wide *d*_*XY*_ estimates reported above; the zero-valued median for *d*_*S*_ reflects the modest divergence between these sister species, with many genes having no observed substitutions of either class. Restricting to the 5,825 orthologs with *d*_*S*_ *>* 0 so that per-gene ratios are defined, the distribution of *d*_*N*_ */d*_*S*_ is strongly skewed toward low values (mean 0.447, median 0.283; Supplementary Fig. S76), consistent with pervasive purifying selection removing nonsynonymous mutations from the majority of coding sequence. Nevertheless, 662 of these orthologs (11.4%) had *d*_*N*_ */d*_*S*_ *>* 1, identifying a substantial tail of genes whose divergence patterns are consistent with relaxed constraint or positive selection.

Turning from windowed divergence to per-gene rates at coding sites, we next compared evolutionary rates between the Z chromosome and the autosomes. Synonymous divergence on the Z (chr17; *n* = 272 orthologs) was significantly reduced relative to the autosomes (*n* = 12,555 orthologs): mean *d*_*S*_ = 0.0082 on the Z versus 0.0100 on the autosomes (onesided Mann-Whitney test, *p* = 0.008; two-sided *p* = 0.016), and a greater fraction of Z-linked orthologs had no observed synonymous substitutions (62.1% versus 54.4%; Fisher’s exact test, OR = 0.73, *p* = 0.012). In contrast, mean nonsynonymous divergence did not differ significantly between the Z and autosomes (*d*_*N*_ = 0.0049 vs. 0.0056; two-sided Mann-Whitney, *p* = 0.26). The reduction of synonymous but not nonsynonymous divergence on the Z echoes the reduced Z-linked *d*_*XY*_ and *π* reported above and, because synonymous sites are largely neutral, motivates a direct estimate of sex-biased mutation rates from the ratio of Z-linked to autosomal synonymous divergence.

Taking this ratio one step further, we used the framework of Miyata et al. (1987) to estimate the degree of sex-biased mutation, in which the male-to-female mutation rate ratio *α*_*m:f*_ = (3*R* − 2)*/*(2 − *R*) with *R* being the ratio of Z-linked to autosomal divergence. Restricting the calculation to synonymous sites at protein-coding genes and bootstrapping across five randomly chosen individuals from each species, we estimated *α*_*m:f*_ = 0.35 (median 0.30; 95% C.I. = 0.03–1.02), implying that the female mutation rate at synonymous sites is roughly 2.8 *×* that of males. This is consistent with the female-biased mutation rate reported by Coffing et al. (2025) in a deeper-timescale comparison between *O. bimaculoides* and *O. sinensis*, though the confidence interval here includes unity and so we interpret this point estimate as suggestive rather than conclusive.

Returning to the 662-gene *d*_*N*_ */d*_*S*_ *>* 1 tail, we asked whether it is enriched for particular classes of genes with respect to functional annotation. The dominant feature of the tail is a striking depletion of functionally annotated, broadly conserved genes: only 7.1% of the 662 orthologs with *d*_*N*_ */d*_*S*_ *>* 1 carried any functional annotation, compared with 27.4% of the remaining 5,163 orthologs with *d*_*S*_ *>* 0 and *d*_*N*_ */d*_*S*_ ≤ 1 (Fisher’s exact test, OR = 0.20, *p* = 1.7 *×* 10^−36^). Genes in the high-*d*_*N*_ */d*_*S*_ tail spanned systematically shorter genomic intervals (median gene span 390 bp vs. 605 bp), and only 203 of 662 (30.7%) carried at least ten combined synonymous and nonsynonymous substitutions. Together these patterns indicate that the bulk of the *d*_*N*_ */d*_*S*_ *>* 1 tail comprises short, lineage-specific, and likely rapidly evolving orphan genes for which individual ratio estimates carry substantial stochastic error, and on which selective constraint may be relaxed rather than positive.

Despite the strong over-representation of incompletely annotated genes in this tail, the minority of *d*_*N*_ */d*_*S*_ *>* 1 orthologs that could be confidently annotated were dominated by gene families with well-documented histories of rapid coding sequence evolution across animals. We note that the functional annotations used in this section were derived from best reciprocal BLAST hits to *Drosophila melanogaster* orthologs (see Methods) and may not directly reflect gene function in cephalopods. The most extreme annotated ratio was observed for a carbohydrate sulfotransferase (*chst11* ; `obimac 0027389`, *d*_*N*_ */d*_*S*_ = 5.55, 31.5 nonsynonymous and 1.5 synonymous substitutions), with additional striking hits at a sodium-dependent amino acid transporter (*SLC6A19* ; `obimac 0001150`, *d*_*N*_ */d*_*S*_ = 3.61), a three-prime repair exonuclease (*Trex2* ; `obimac 0011880`, *d*_*N*_ */d*_*S*_ = 2.82), and a cystinosin lysosomal transporter (*CTNS* ; `obimac 0027637`, *d*_*N*_ */d*_*S*_ = 2.76). Classical targets of positive selection involving gamete recognition and fertilization appeared in the tail, most notably the sperm-egg fusion receptor *Izumo1r* /Juno (`obimac 0025275`, *d*_*N*_ */d*_*S*_ = 1.69), alongside immunity- and defense-associated loci including a ctenitoxin-like venom homolog (`obimac 0008163`, *d*_*N*_ */d*_*S*_ = 1.37), melanotransferrin (*MELTF* ; `obimac 0009523`, *d*_*N*_ */d*_*S*_ = 1.40), and glutathione peroxidase 1 (*GPX1* ; `obimac 0003550`, *d*_*N*_ */d*_*S*_ = 1.17), and a second DNA-repair factor, the RAD51 paralog *Rad51d* (`obimac 0003405`, *d*_*N*_ */d*_*S*_ = 1.32). A small cluster of peptidyl-prolyl isomerases and co-chaperones also appeared above the *d*_*N*_ */d*_*S*_ = 1 threshold, including *PPIB* (`obimac 0000088`, *d*_*N*_ */d*_*S*_ = 1.09), *FKBP8* (`obimac 0007845`, *d*_*N*_ */d*_*S*_ = 1.08), and the SNARE homolog *YKT6* (`obimac 0006315`, *d*_*N*_ */d*_*S*_ = 1.11). Formal tests of category enrichment based on keyword matching of annotations were underpowered by the low overall annotation rate in this tail; the only functional class that approached significance was immune/defense/toxin-related genes (2/662 vs. 2/5,163; one-sided Fisher’s exact *p* = 0.066). The high-*d*_*N*_ */d*_*S*_ tail was not enriched on the Z chromosome (10/662 Z-linked in the tail vs. 93/5,163 Z-linked in the rest; Fisher’s exact *p* = 0.75), consistent with the absence of a faster-Z signal in the genome-wide distribution of *d*_*N*_ */d*_*S*_. We therefore interpret the *d*_*N*_ */d*_*S*_ *>* 1 tail as comprising two overlaid populations: a dominant mass of short, lineage-restricted genes whose elevated ratios are most parsimoniously explained by relaxed purifying selection and stochastic noise, and a smaller but functionally coherent subset of gamete-recognition, defense, DNA-repair, and cell-surface-modifying genes whose individual ratios are consistent with episodic positive selection of the type repeatedly documented across animal phyla. Distinguishing drift from selection in these two populations motivates the polymorphism-aware analysis in the next section.

### Positive selection on protein coding genes

In addition to examining the genome-wide signatures of linked selection described above, we directly assessed evidence of positive selection on protein coding genes using McDonald-Kreitman (MK) tests. Using the same set of single-copy orthologs described above, we conducted MK tests with `MKado` using each species alternately as the ingroup and outgroup. We estimated genome-wide *α*—the proportion of nonsynonymous substitutions fixed by positive selection (distinct from the male-to-female mutation rate ratio *α*_*m:f*_ introduced in the previous section; unsubscripted *α* refers throughout this section to the MK adaptation proportion)—using the asymptotic MK method, and tested individual genes for evidence of positive selection by computing the neutrality index (NI).

Consistent with the linked selection results, we found stronger evidence for positive selection in *O. bimaculatus* (fig. 5A). The estimated genome-wide *α* for *O. bimaculatus* was 0.242 (95% CI: 0.120–0.367), compared to 0.064 (95% CI: − 0.033–0.163) for *O. bimaculoides*. While the confidence intervals overlap slightly, the point estimate for *O. bimaculatus* suggests that roughly a quarter of nonsynonymous substitutions were driven by positive selection, whereas the estimate for *O. bimaculoides* is not significantly different from zero. This pattern is expected under the nearly neutral theory: in the smaller *O. bimaculoides* population, weakly beneficial mutations are more likely to be lost to drift, and mildly deleterious mutations are more likely to fix, both of which reduce *α*. We also compared *α* between Z-linked genes and autosomal genes. The estimated *α* for *O. bimaculoides* Z genes was 0.537 (95% CI: -0.1141 - 1.2030) and for autosomal genes was 0.0628 (95% CI: -0.0358 - 0.1638). The estimated *α* for *O. bimaculatus* Z genes was 0.1891 (95% CI: -0.2334 - 0.6220) and for autosomal genes was 0.2437 (95% CI: 0.1266 - 0.3636). Of these four results, only the *O. bimaculatus* autosomes show an estimated *α* significantly greater than zero, consistent with reduced selection efficiency on the *O. bimaculoides* and the Z chromosome as a result of a lower effective population size.

**Figure 5:**
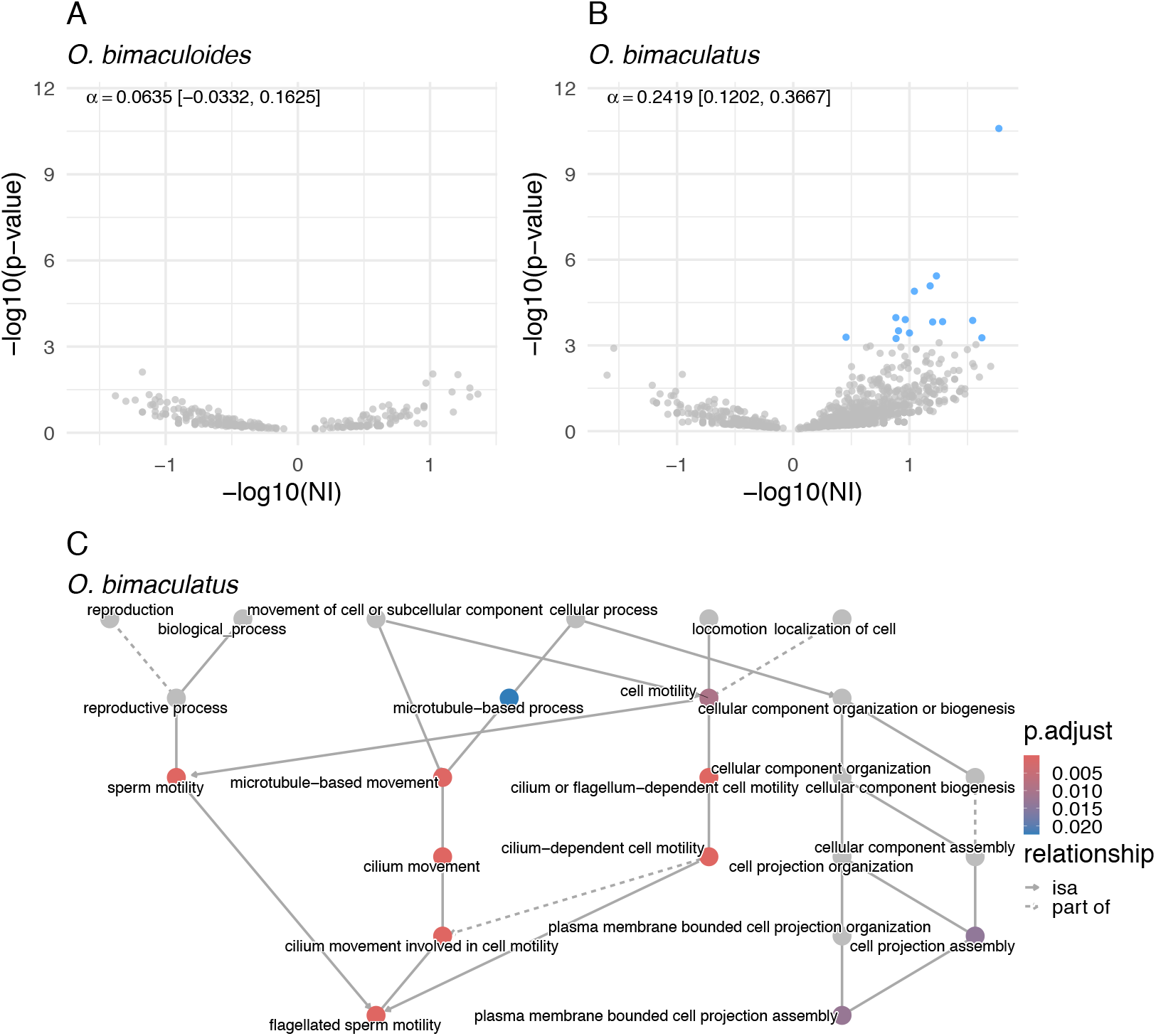
Evidence of positive selection on protein coding genes. Volcano plots showing neutrality index (NI) against *p*-value for McDonald-Kreitman tests for individual genes in *O. bimaculoides* (A) and *O. bimaculatus* (B), with significant genes after Benjamini-Hochberg FDR correction (*q <* 0.05) shown in blue (no genes were significant after FDR correction in *O. bimaculoides*). Genome-wide estimate and 95% confidence intervals of *α* for each species are printed in the top left of each panel. (C) Directed acyclic graph of significantly enriched GO terms among genes with significant MK test results.

At the individual gene level, we conducted MK tests across all one-to-one orthologs and corrected for multiple testing using the Benjamini-Hochberg procedure. No individual genes were significant in *O. bimaculoides* after correction (Supplementary Fig. 5A), while *O. bimaculatus* yielded many genes with significant neutrality indices less than one (fig. 5B; Table 2), indicating an excess of nonsynonymous divergence consistent with positive selection. Gene ontology enrichment analysis of the significant MK test genes in *O. bimaculatus* revealed enrichment for terms related to cell motility, sperm motility, and cilium-dependent movement (fig. 5C). We note that these GO annotations are derived from *Drosophila melanogaster* orthologs and may not directly reflect gene function in cephalopods; nonetheless, the enrichment for motility-related genes is intriguing given the planktonic paralarval dispersal stage unique to *O. bimaculatus*. One notable gene that appeared across many of the motility related GO terms (see supplementary materials for all significant GO terms and affiliated genes) was Dynein, axonemal, heavy chain 3 (DNAH3, FBgn0035581). DNAH3 is is expressed in the sperm flagellum and is required for sperm motility in *Drosophila*, and is associated with male sterility when mutated in humans (Meng et al. 2024). The conservation of this gene’s function across *Drosophila* and mammals suggests the function may be conserved in octopuses as well, and its signature of positive selection in *O. bimaculatus* may reflect adaptation related to sperm motility and perhaps dispersal during the planktonic larval stage.

**Table 2:** McDonald-Kreitman Tests with significant results after FDR correction for *O. bimaculatus*. Columns are the best reciprocal blast hit gene IDs between *O. bimaculoides* and *Drosophila, D*_*n*_: number of fixed nonsynonymous substitutions; *D*_*s*_: number of fixed synonymous substitutions; *P*_*n*_: number of polymorphic nonsynonymous sites; *P*_*s*_: number of polymorphic synonymous sites; *α*: proportion of substitutions fixed by positive selection; NI: neutrality index; *p*-adjusted: FDR-corrected p-value.

| Octopus Gene | Drosophila Gene | $D_n$ | $D_s$ | $P_n$ | $P_s$ | $\alpha$ | NI | $p$-adjusted |
| --- | --- | --- | --- | --- | --- | --- | --- | --- |
| obimac_0000970 | FBpp0290517 | 42 | 10 | 2 | 28 | 0.983 | 0.017 | $2.63 \times 10^{-8}$ |
| obimac_0011888 | FBpp0301170 | 23 | 10 | 6 | 21 | 0.876 | 0.124 | $3.16 \times 10^{-2}$ |
| obimac_0022401 | FBpp0297181 | 15 | 1 | 6 | 14 | 0.971 | 0.029 | $1.72 \times 10^{-2}$ |
| obimac_0024693 | FBpp0110135 | 20 | 12 | 2 | 19 | 0.937 | 0.063 | $1.72 \times 10^{-2}$ |
| obimac_0027271 | FBpp0306552 | 18 | 22 | 4 | 45 | 0.891 | 0.109 | $1.72 \times 10^{-2}$ |
| obimac_0028054 | FBpp0288420 | 40 | 48 | 2 | 24 | 0.900 | 0.100 | $3.43 \times 10^{-2}$ |
| obimac_0002704 | FBpp0088796 | 18 | 3 | 1 | 7 | 0.976 | 0.024 | $4.20 \times 10^{-2}$ |
| obimac_0002368 | FBpp0293284 | 21 | 20 | 4 | 42 | 0.909 | 0.091 | $3.26 \times 10^{-3}$ |
| obimac_0002253 | FBpp0301058 | 30 | 29 | 2 | 33 | 0.941 | 0.059 | $1.90 \times 10^{-3}$ |
| obimac_0006457 | FBpp0428257 | 18 | 7 | 2 | 15 | 0.948 | 0.052 | $1.72 \times 10^{-2}$ |
| obimac_0008052 | FBpp0302015 | 41 | 17 | 6 | 19 | 0.869 | 0.131 | $1.72 \times 10^{-2}$ |
| obimac_0015906 | FBpp0312333 | 17 | 57 | 2 | 101 | 0.934 | 0.066 | $2.82 \times 10^{-3}$ |
| obimac_0017267 | FBpp0084818 | 23 | 10 | 6 | 20 | 0.870 | 0.130 | $4.20 \times 10^{-2}$ |
| obimac_0019307 | FBpp0292877 | 67 | 37 | 35 | 55 | 0.649 | 0.351 | $4.20 \times 10^{-2}$ |

**Table 3:**
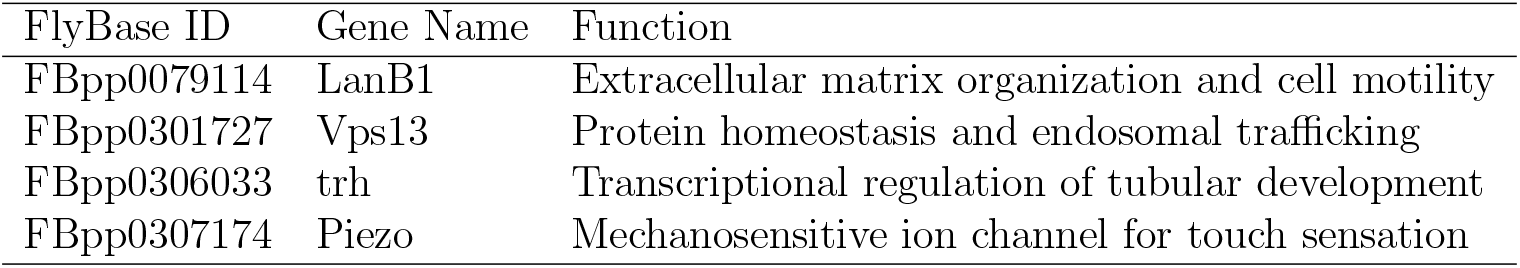
Genes with evidence of positive selection from McDonald-Kreitman tests and overlapping with selective sweeps in *O. bimaculatus*. McDonald-Kreitman tests were not significant after correction. Selective sweeps were identified using `diploS/HIC` with a probability threshold ≥ 0.95. Functional information was collected from FlyBase (Jenkins et al. 2022).

| FlyBase ID | Gene Name | Function |
| --- | --- | --- |
| FBpp0079114 | LanB1 | Extracellular matrix organization and cell motility |
| FBpp0301727 | Vps13 | Protein homeostasis and endosomal trafficking |
| FBpp0306033 | trh | Transcriptional regulation of tubular development |
| FBpp0307174 | Piezo | Mechanosensitive ion channel for touch sensation |

### Selective sweeps

We complemented the MK-based analysis with a scan for selective sweeps using `diploS/HIC`, a machine learning classifier trained on coalescent simulations (Kern and Schrider 2018). For each species, we trained the classifier using simulations with `discoal` (Kern and Schrider 2016) under neutral, hard sweep, and soft sweep conditions, incorporating the species-specific demographic histories inferred by `SMC++` (see above). We simulated in 110 Kb windows (divided by `diploS/HIC` into 11 sub-windows of 10 Kb) and classified 10 Kb subwindows, varying selection coefficients from 0.001 to 0.01 (see Materials and Methods).

The classifier trained under *O. bimaculatus* demographic parameters performed well at distinguishing sweep, linked-to-sweep, and neutral regions, though it had limited power to differentiate hard from soft sweeps (Supplementary Fig. S77). Performance was substantially poorer for *O. bimaculoides* (Supplementary Fig. S78), likely reflecting the combined effects of its smaller *N*_*e*_ and our smaller sample size (*n* = 6), both of which reduce the footprint of sweeps in patterns of variation. Because of this limited power, we restricted downstream analysis to sweeps called in *O. bimaculatus* and to sweep regions shared between both species, and we did not attempt to distinguish hard from soft sweeps. Naturally, the number of inferred and shared sweeps depends on the chosen probability threshold for classification. At a relaxed threshold (classification probability of being either a hard or soft sweep ≥ 0.5), we identified 5,872 sweep regions in *O. bimaculatus* and 2,811 for *O. bimaculoides*; 250 of which were shared between both species. In contrast, at a stringent threshold of 0.95, we identified 1,281 sweep regions in *O. bimaculatus* and none in *O. bimaculoides* (See Fig. S80 and Fig. S81 for all probability thresholds). The density of inferred sweeps (using a cutoff of 0.5) was negatively correlated with recombination rate in *O. bimaculatus* and uncorrlated in *O. bimaculoides* (Spearman’s *ρ* = -0.113, *p* ≪ 0.001; and *ρ* = 0.015, *p* = 0.71, respectively), consistent with the expectation that sweeps leave stronger signatures in regions of low recombination; while sweep density was slightly negatively correlated with coding sequence density in *O. bimaculatus* and slightly positively correlated in *O. bimaculoides* (Spearman’s *ρ* = -0.135, *p* ≪ 0.001; and *ρ* = 0.0814, *p* = 0.0382, respectively). In both species, there is a “u” shaped trend, suggesting a complex interplay of selection and functional content. (Supplementary Fig. S79).

We identified protein coding genes overlapping with high-confidence sweep regions (at a probability threshold ≥ 0.95) and tested for enrichment of gene ontology categories using *D. melanogaster* orthologs as the annotation source (see Materials and Methods). Notably, for sweeps in *O. bimaculatus*, we found significant enrichment for GO terms related to brain and neuron development (Supplementary Fig. S82). Likewise, for sweep regions shared between both species (at a probability threshold ≥ 0.5), enrichment was found for compound eye development and instar larval or pupal morphogenesis (Supplementary Fig. S83). In particular there were 14 genes that accounted for enrichment for R7 cell differentiation shared on both species. R7 cells are a type of photoreceptor cell in the compound eye of *Drosophila* that are responsible for detecting ultraviolet light and are involved in color vision (**chou2010drosophila**), including well-studied genes such as *daughter of sevenless* (FBgn0016794), *spineless* (FBgn0003513), and *shaven* (FBgn0005561). Further investigation of these genes, their tissue-specific expression, and their functions in octopuses may provide insights into the evolution of visual systems in cephalopods. These results suggest that genes underlying traits characteristic of coleoid cephalopods— particularly those involved in the nervous system and visual system— continue to be targets of positive selection on recent evolutionary timescales.

## Discussion

### Estimates of diversity, divergence, and demographic history consistent with life history differences between Two-spot octopus species

We found *O. bimaculoides* to have lower nucleotide diversity (*π*) and a consistently lower effective population size than *O. bimaculatus* (figs. 2 and 3). These estimates are consistent with the differences in life history and reproductive strategies between these species, as well as with previous findings based on mitochondrial genes and microsatellite loci (Domínguez-Contreras, Munguia-Vega, Ceballos-Vázquez, Arellano-Martínez, et al. 2018). *O. bimaculatus* has a planktonic paralarval drift stage, unlike *O. bimaculoides*, which provides more opportunity for dispersal from the hatching site and can potentially increase gene flow between populations. Additionally, *O. bimaculatus* lays far more eggs per clutch than *O. bimaculoides*, which could contribute to a larger population size and higher nucleotide diversity levels. Tajima’s *D* is slightly positive in *O. bimaculoides*, consistent with a history of declining effective population size, whereas the near-zero average Tajima’s *D* in *O. bimaculatus* is consistent with our estimated constant-sized population history.

Genetic differentiation between the species is relatively low and indicative of a recent split time, as expected given their near-identical external morphology, overlapping California ranges, and the prior mitochondrial evidence for limited divergence between these sister taxa (Domínguez-Contreras, Munguia-Vega, Ceballos-Vázquez, Arellano-Martínez, et al. 2018). For instance, our estimates of *d*_*XY*_ between the Two-spot octopus species (average windowed *d*_*XY*_ = 0.012) are roughly on the order of human-chimpanzee divergence (approximately 0.01) (Rodrigues, Kern, and Ralph 2024). Assuming the mutation rate of *µ* = 2.4 *×* 10^−9^ per bp per generation, as estimated from *H. maculosa*, our average *d*_*XY*_ estimate implies divergence roughly 2.5 million generations ago (∼ 5M years). This is likely an overestimate of the split time, as population histories derived using an SMC estimator (`SMC++`) show considerably younger estimates (from ∼ 375k–600k years).

Comparing absolute and relative measures of divergence is also informative. Our estimates of *F*_*ST*_ across the genome are quite high relative to *d*_*XY*_ (average windowed *F*_*ST*_ = 0.705, fig. 2). *F*_*ST*_ is a measure of the proportion of genetic diversity that is attributable to differences among populations. However, *F*_*ST*_ is well known to be sensitive to levels of within-population variation (e.g., nucleotide diversity *π*) (Charlesworth 1998). Because *O. bimaculoides* has globally reduced levels of nucleotide diversity compared to *O. bimaculatus, F*_*ST*_ values are inflated relative to *d*_*XY*_ . Overall, what emerges is a history of relatively recent divergence between sister clades, one of which (*O. bimaculoides*) experienced a significant population decline, along with the expected reduction in diversity, after speciation.

There is currently no evidence of hybridization between the Two-spot octopus species. A preliminary study placed a female *O. bimaculoides* in a tank with a male *O. bimaculatus* individual (and vice versa) and found that they neither interacted nor made any mating attempts, suggesting tentative evidence of pre-mating isolation (Pickford and McConnaughey 1949). Although this study was rudimentary and had a small sample size, its results, combined with a lack of genomic evidence for hybridization and high levels of *F*_*ST*_, suggest that behavioral barriers to mating may exist. Such barriers may be reinforced by the largely nonoverlapping reproductive seasons of the two species in the portions of their ranges where they co-occur (Domínguez-Contreras, Munguia-Vega, Ceballos-Vazquez, Arellano-Martínez, et al. 2018), which would further restrict opportunities for hybridization even in the absence of behavioral isolation. These results raise questions about the nature of pre- and post-zygotic barriers and the factors driving speciation in these two species. Disentangling the nature and timing of speciation between *O. bimaculoides* and *O. bimaculatus*—especially the mechanisms of reproductive isolation and the roles of natural selection and geographic separation—is a valuable area for further research.

### Z chromosome polymorphism

Sex chromosomes are expected to show distinct evolutionary dynamics compared to autosomes, largely due to differences in selection pressures, genetic drift, and effective population sizes (Vicoso and Charlesworth 2006; Charlesworth, Coyne, and Barton 1987). The Fast-Z effect (or Fast-X effect in XY systems) predicts that coding regions of the Z chromosome will evolve faster than those on autosomes, leading to higher *F*_*ST*_ —a relative measure of genetic differentiation between populations—on the Z compared to autosomes, a pattern found in several species (Charlesworth, Coyne, and Barton 1987; Ellegren et al. 2012; Oyler-McCance et al. 2015; Mank, Axelsson, and Ellegren 2007; Wright et al. 2015). One theory proposed to drive this pattern is that the heterogametic sex exposes recessive mutations to selection—intensifying both purifying selection against deleterious alleles and positive selection on beneficial alleles (Charlesworth, Coyne, and Barton 1987).

Between the Two-spot octopus species, we found *F*_*ST*_ to be higher on the Z chromosome compared to autosomes, which is consistent with the Fast-Z effect hypothesis. However, in contrast to this pattern, we observed a significantly lower *d*_*XY*_ and reduced nucleotide diversity on the Z chromosomes compared to the autosomes in both species. The combination of these findings is consistent with the hypothesis that the Z chromosome is prone to purifying selection of recessive, mildly deleterious alleles, which quickly removes diversity (Charlesworth, Coyne, and Barton 1987; Ellegren et al. 2012; Oyler-McCance et al. 2015). These results are also consistent with our previous findings that divergence is reduced on the Z chromosomes when looking at deeper time scales between *O. bimaculoides* and *O. sinensis* (Coffing et al. 2025). Additionally, the lower effective population size of the Z chromosome may contribute to these patterns. Overall, our results are similar to those found in closely related greenish warbler species, in which the Z chromosomes have lower nucleotide diversity and *d*_*XY*_ compared to autosomes combined with higher *F*_*ST*_ (Irwin et al. 2016).

Our coding-sequence comparisons help break down this Z-linked signal. Per-gene divergence on the Z showed significantly reduced synonymous divergence (*d*_*S*_) relative to autosomes, but nonsynonymous divergence (*d*_*N*_) statistically indistinguishable from autosomal. The depression of Z-linked *d*_*XY*_ is therefore driven largely by reduced neutral divergence rather than by altered protein-coding evolution. Under a classical Fast-Z interpretation, we would expect the Z to show elevated *d*_*N*_ */d*_*S*_ relative to autosomes, reflecting more efficient fixation of beneficial alleles and purging of deleterious ones in the hemizygous sex (Charlesworth, Coyne, and Barton 1987; Mank, Axelsson, and Ellegren 2007), yet we find no such enrichment in the genome-wide per-gene *d*_*N*_ */d*_*S*_ distribution. The elevated *F*_*ST*_ we observe on the Z is thus most parsimoniously interpreted not as a Fast-Z protein-evolution signature but as a consequence of reduced within-species diversity combined with the Z’s smaller effective population size (Charlesworth 1998; Cruickshank and Hahn 2014).

A second, complementary explanation for the reduced *d*_*S*_ on the Z is a sex-biased mutation rate. Applying the Miyata et al. (1987) framework to the ratio of Z-linked to autosomal synonymous divergence, we estimated a male-to-female mutation rate ratio of *α*_*m:f*_ ≈ 0.35, implying a female mutation rate roughly 2.8 *×* that of males at synonymous sites. Although the 95% confidence interval around this point estimate includes unity and so we interpret it cautiously, its direction is in close agreement with the female-biased mutation rate independently inferred by Coffing et al. (2025) from the deeper-timescale comparison between *O. bimaculoides* and *O. sinensis*, in which synonymous divergence and windowed divergence both yielded Z/autosome ratios consistent with a female rate substantially exceeding the male rate. Female-biased mutation contrasts with the male-biased pattern typical of amniotes (Bergeron et al. 2023) and could reflect differences in germline cell-division dynamics or repair efficiency between male and female cephalopods; resolving its mechanistic basis remains an interesting target for future work.

Our recombination rate estimates offer an additional window into Z chromosome biology. `ReLERNN` estimates the effective (population-scaled) recombination rate *ρ* = 4*N*_*e*_*r* from patterns of linkage disequilibrium, so both the effective population size and the sexaveraged per-generation rate contribute. In a ZZ/ZO system, the Z chromosome recombines only in ZZ males—hemizygous ZO females have no homolog with which to recombine— and the Z spends two-thirds of its time in males (who carry two copies) and one-third in females (who carry one), giving a sex-averaged per-generation rate of (2*/*3)*r*_*m*_. Combined with the three-quarters effective population size of the Z (Vicoso and Charlesworth 2006; Charlesworth 2009), the expected ratio of effective recombination rates depends critically on whether females recombine on the autosomes. If females recombine at the same rate as males, then *ρ*_*Z*_*/ρ*_*A*_ = (3*/*4) *×* (2*/*3) = 1*/*2, and the Z should appear as a clear outlier with reduced effective recombination. However, if females lack recombination entirely (achiasmate meiosis), the sex-averaged autosomal rate becomes *r*_*m*_*/*2, and the ratio becomes *ρ*_*Z*_*/ρ*_*A*_ = (3*/*4) *×* (2*/*3)*r*_*m*_ */* (1*/*2)*r*_*m*_ = (3*/*4) *×* (4*/*3) = 1: the reduced *N*_*e*_ of the Z and its elevated fraction of time in the recombining sex cancel exactly. Our observation that Z recombination rate estimates are indistinguishable from the autosomes (fig. 4A) matches the achiasmate prediction precisely.

Achiasmate meiosis in the heterogametic sex has evolved independently at least 29 times across animals, a pattern formalized as the Haldane–Huxley rule (Haldane 1922; Huxley 1928; Burt, Bell, and Harvey 1991). The original explanation proposed that reduced autosomal recombination in the heterogametic sex arises as a pleiotropic consequence of selection against recombination between the sex chromosomes. Lenormand and Dutheil (2005) offered an alternative hypothesis, suggesting that loss-of-function mutations eliminating recombination can spread only when they affect the heterogametic sex, because maintaining recombination in the homogametic sex is necessary to preserve proper segregation of sex chromosomes. They further proposed that haploid selection—selection acting on the haploid products of meiosis— may drive heterochiasmy more broadly, predicting reduced recombination in the sex with more intense haploid selection. Well-characterized examples of achiasmy include *Drosophila* males (XY heterogamety), in which recombination is absent in males but occurs normally in females (Comeron, Ratnappan, and Bailin 2012; John, Vinayan, and Varghese 2016), and Lepidoptera females (ZW heterogamety), which are completely achiasmate (Suomalainen 1966). Similar to Lepidoptera, female coleoid cephalopods are heterogametic, making female achiasmate meiosis a plausible hypothesis. To our knowledge, sex-specific recombination rates have not been directly measured in any cephalopod. Resolving this question through genetic crosses or gamete-based recombination mapping would have broad implications for interpreting patterns of linked selection and sex chromosome evolution across this clade.

### Stronger drift likely limits selection in *O. bimaculoides*

Estimating effective recombination rates with `ReLERNN` allowed us to test for signatures of linked selection—the reduction of neutral diversity by nearby selected variants through background selection and selective sweeps (Lewontin 1974; Begun and Aquadro 1992; Corbett-Detig, Hartl, and Sackton 2015). A positive correlation between recombination rate and nucleotide diversity is a hallmark of linked selection: in regions of low recombination, the effects of selection at linked sites extend over larger physical distances, reducing neutral diversity. In regions of high recombination, linkage breaks down more rapidly and neutral diversity is less affected (Begun and Aquadro 1992; Corbett-Detig, Hartl, and Sackton 2015). We observe this pattern clearly in *O. bimaculatus* but not in *O. bimaculoides* (fig. 4B). Similarly, the negative relationship between coding sequence density and *π*—expected when regions with more targets of selection experience stronger diversity reductions—is more pronounced in *O. bimaculatus* (fig. 4C).

This contrast between species is consistent with theoretical predictions and comparative evidence that linked selection is more effective in species with larger effective population sizes (Corbett-Detig, Hartl, and Sackton 2015; Buffalo 2021). Corbett-Detig, Hartl, and Sackton (2015) showed across a broad taxonomic survey that species with larger census population sizes exhibit stronger correlations between recombination and polymorphism, reflecting a greater genome-wide impact of natural selection on linked neutral variation. In *O. bimaculoides*, the smaller *N*_*e*_ means that genetic drift dominates the fate of linked neutral variants, overwhelming the signal of selection and erasing the recombination–diversity correlation. This interpretation is further supported by the patterns of *F*_*ST*_ across the genome. Cruickshank and Hahn (2014) demonstrated that heterogeneous landscapes of *F*_*ST*_ between closely related species often reflect variation in within-population diversity driven by linked selection, rather than differential gene flow. Applied to the Two-spot octopus species, the regions of elevated *F*_*ST*_ we observe are likely shaped by the reduced diversity within *O. bimaculoides* due to its population decline, rather than by barriers to gene flow, consistent with the moderate and relatively uniform *d*_*XY*_ across the genome.

### Stronger evidence for positive selection in *O. bimaculatus*

Our McDonald-Kreitman tests reinforce the conclusion that positive selection acts more effectively in *O. bimaculatus* than in *O. bimaculoides*. The estimated genome-wide *α*— the proportion of nonsynonymous substitutions driven to fixation by positive selection—is substantially higher in *O. bimaculatus* than in *O. bimaculoides*, although the confidence intervals overlap slightly (fig. 5A). After correction for multiple testing, only *O. bimaculatus* yielded individual genes with significant neutrality indices consistent with positive selection (fig. 5B); no genes were significant in *O. bimaculoides*. These results dovetail with the linked selection analyses: in both cases, the smaller effective population size of *O. bimaculoides* appears to limit the efficiency of natural selection, allowing genetic drift to dominate the fate of weakly selected mutations.

A complementary, divergence-only view of positive selection comes from the per-gene *d*_*N*_ */d*_*S*_ distribution. Of the 5,825 one-to-one orthologs with *d*_*S*_ *>* 0, 662 have *d*_*N*_ */d*_*S*_ *>* 1, but this set is dominated by short, poorly annotated, lineage-restricted genes and almost certainly includes a large component of stochastic noise and relaxed constraint rather than true episodic adaptation. The MK framework is considerably better powered for inferring positive selection at individual loci because it incorporates within-species polymorphism in addition to between-species divergence, effectively calibrating each gene’s substitution counts against its neutral expectation. That said, the minority of high-*d*_*N*_ */d*_*S*_ orthologs that are confidently annotated is biologically coherent, populated by classical targets of rapid coding evolution across animals including gamete-recognition factors (e.g., *Izumo1r* /Juno), immunity- and defense-associated loci, and DNA-repair genes. These candidates provide a useful supplement to the MK-significant gene set and would be informative targets for focused follow-up work in these species.

Gene ontology enrichment analysis of the significant MK test genes in *O. bimaculatus* revealed enrichment for terms related to cell motility, sperm motility, and cilium-dependent movement (fig. 5C). Because these annotations are derived from *Drosophila melanogaster* orthologs, we cannot say with certainty what roles the corresponding proteins play in cephalopods. Nonetheless, the enrichment for motility-related genes is intriguing in light of the planktonic paralarval dispersal stage unique to *O. bimaculatus*, and warrants further investigation into whether selection on larval motility has contributed to the divergence between the Two-spot octopus species.

Our selective sweep analysis provides a complementary view of positive selection on more recent evolutionary timescales. Sweeps detected by `diploS/HIC` reflect selection that has occurred within the last ∼4 ln(2*N*_*e*_)*/s* generations (∼ 10^4^–10^6^ years ago), offering a window into ongoing adaptive evolution. In *O. bimaculatus*, genes overlapping high-confidence sweep regions were enriched for GO terms related to brain and neuron development, while sweep regions shared between both species were enriched for compound eye development (Supplementary Figs. S82 and S83). These findings suggest that genes underlying traits characteristic of coleoid cephalopods—particularly those involved in the nervous and visual systems—continue to be targets of positive selection. Given the ecological importance of complex cognition and vision in octopuses, evidence of ongoing selection on these gene categories is noteworthy and merits further study.

We acknowledge that our power to detect sweeps in *O. bimaculoides* was limited by both its smaller effective population size and our smaller sample size (*n* = 6), both of which reduce the genomic footprint of selective sweeps. It is therefore likely that some sweeps in *O. bimaculoides* went undetected. However, this limitation is itself consistent with the broader pattern: natural selection is less efficient in populations with small effective sizes, and the reduced signals of linked selection in *O. bimaculoides* suggest that drift, rather than selection, is the dominant evolutionary force shaping diversity in this species.

## Conclusions

To our knowledge, we have presented the first whole-genome re-sequencing dataset for a cephalopod analyzed in a population genetics framework. A central finding of this study is that the demographic asymmetry between these sibling species—a large, stable effective population size in *O. bimaculatus* versus a smaller, declining one in *O. bimaculoides*—has pervasive consequences for the efficacy of natural selection. Signatures of linked selection, a higher proportion of adaptive substitutions, and more detectable selective sweeps in *O. bimaculatus* all point to a species in which selection operates more efficiently, consistent with theoretical expectations for larger populations. In *O. bimaculoides*, drift appears to be the dominant force shaping genomic variation.

The ancient ZZ/ZO sex determination system in these species offers a novel framework for studying sex chromosome evolution. While we observe elevated *F*_*ST*_ on the Z chromosome, the accompanying reductions in *d*_*XY*_ and nucleotide diversity suggest that this pattern is driven primarily by purifying selection against recessive deleterious alleles rather than accelerated adaptive evolution. Our recombination rate estimates further provide indirect evidence for achiasmate meiosis in the heterogametic sex, a prediction that can be tested directly through genetic crosses or gamete-based recombination mapping. More broadly, the Z chromosome provides a natural contrast to the autosomes for understanding how differences in effective population size and recombination regime shape patterns of diversity and divergence.

Finally, the enrichment of selective sweeps near genes involved in brain and eye development suggests that traits characteristic of coleoid cephalopods continue to be shaped by positive selection on recent evolutionary timescales. As genomic resources for cephalopods expand, comparative population genomic studies across additional species will be essential for understanding the evolutionary forces that have shaped this remarkable clade.

## Materials and Methods

### Sample collection and resequencing

*O. bimaculoides* and *O. bimaculatus* samples were collected from the southern coast of California between 2019 and 2022. Various tissue types were either flash frozen or stored in RNA*later* (Invitrogen; Supplementary Table S1). DNA extraction, sample preparation, and sequencing for all samples were conducted by the University of Oregon Genomics & Cell Characterization Core Facility (GC3F). A separate sequencing library was prepared for each species. Genomic DNA was extracted, pooled, and sequenced on two Illumina NovaSeq 6000 S4 lanes. In total, our study includes six *O. bimaculoides* and thirteen *O. bimaculatus* individuals (Supplementary Table S1). Sequencing data from four of the *O. bimaculatus* individuals were published previously (Coffing et al. 2025).

### Variant calling

We used `SNPArcher` v1.0 (Mirchandani et al. 2024) to call variants separately in each species using the newest *O. bimaculoides* genome assembly (Coffing et al. 2025) as the reference. `SNPArcher` maps reads to the reference with `bwa-mem` (Vasimuddin et al. 2019) and generates a VCF using `GATK` (McKenna et al. 2010). We conducted our own filtering scheme on `SNPArcher`’s raw output VCFs using `BCFtools` v.1.20 (Danecek et al. 2021). We filtered out all repetitive sites with the repeat map that was generated with the `RepeatMasker` (Smit, Hubley, and Green 2013) and `RepeatModeler` (Flynn et al. 2020) pipeline in Coffing et al. (2025). We also set genotypes to missing (i.e., `./.`) according to depth or the allele depth to depth ratio, depending on the genotype. For all homozygous sites, we set genotypes to missing that had an allelic depth below 10. For heterozygous sites, we set sites to missing that had an allelic depth below 10 and allele balance below 30%. Next, we removed sites that did not pass a set of thresholds: quality below 20, quality by depth below 10, mapping quality below 40, genotype missingness above 20%, depth higher than 2 *×* the overall average depth. We additionally removed all indels. In total, we masked 61.08% and 65.06% of the total genome in *O. bimaculoides* and *O. bimaculatus*, respectively. All sites that did not pass our filtering scheme were included in a mask bed file that was used in downstream analyses. Overall, filtering resulted in 1.77 million SNPs in the *O. bimaculoides* VCF and 14.59 million SNPs in the *O. bimaculatus* VCF. In addition to the individual VCFs and masks for each species, we generated a merged VCF with 14.52 million SNPs using `BCFtools` merge and a merged mask using `BEDtools` v2.31.1 merge (Quinlan and Hall 2010). In all final VCFs, we kept invariant sites.

### Depth calculations

Using the bam files that were generated with `SNPArcher` (Mirchandani et al. 2024), we calculated read depth in 500,000 bp windows using `samtools depth` v1.19.2 (H. Li et al. 2009). We examined depth across chromosomes for each individual and found that two of the *O. bimaculatus* individuals that were marked as female (Oblat F 7 and Oblat F 8) did not have half coverage on the Z chromosome. This led us to conclude there was an error in sexing these individuals in the field, and we adjusted their sex label to Male (Supplementary Table S1).

### Calculation of population genetics statistics

We calculated nucleotide diversity (*π*), Tajima’s *D, d*_*XY*_, and Weir and Cockerham *F*_*ST*_ in non-overlapping 1 Mb windows using the Python package `scikit-allel` v1.3.13 (Miles et al. 2024). The mask we generated from our filtering scheme (see above) was used as input for the negative accessibility mask. Population statistics were calculated on the independent VCFs for each species. We used `scikit-allel` to generate PCA plots for each chromosome from the merged VCF using 40,000 randomly-sampled biallelic SNPs with a minor allele count of at least two.

### Demographic analysis

We used `SMC++` v1.15.2 (Terhorst, Kamm, and Song 2017) to estimate the history of effective population size and divergence time of the two octopus species. We input the merged VCF, containing 14.52 million SNPs, into the `vcf2smc` command using the merged mask as input for missing data. This output a separate smc file for each chromosome. The `estimate` command of `SMC++` was used to estimate population size history of each species with a mutation rate of 2.4 *×* 10^−9^ as reported for the Southern blue-ringed octopus (*Hapalochlaena maculosa*) (Whitelaw et al. 2022), which is the closest related species with a reported mutation rate estimate. To infer the divergence time of the two species, we independently generated a joint site frequency spectrum for each species using the `vcf2smc` command. Finally, the command `split` was run to refine the marginal estimates to infer the joint demography. Demographic history was estimated independently for each of the 30 octopus chromosomes. The median and upper and lower quantiles were estimated from the 30 independent history traces.

### Estimating recombination landscapes and the effect of linked selection

We used `ReLERNN` (Adrion, Galloway, and Kern 2020) (https://github.com/kr-colab/relernnnextflow) to infer local recombination rates along the genome separately for *O. bimaculoides* and *O. bimaculatus*. For efficiency, training and predictions were split up into 12 Mb windows along the chromosomes and we provided the same accessibility mask as used in estimating the population genetic summary statistics to account for missing data (see above). We tested a difference in genome-wide recombination rate between the two species using a Mann-Whitney U test. Estimating recombination allowed us to assess how local nucleotide diversity varies with recombination rate between the two species, which in turn allowed us to assess the relative impacts of linked selection acting on the genome of the two species. We did this by comparing nucleotide diversity in 1 Mb windows to estimates of local recombination rate and to the density of coding sequences in the same window.

### Inferring selection on coding sequences

To detect evidence of selection at the protein and genome-wide levels, we conducted McDonald-Kreitman (MK) tests (McDonald and Kreitman 1991) using single-copy orthologs between individuals of each species. To generate inputs for `MKado` v0.4, we first used custom scripts and our species-specific VCFs to map SNPs onto the coding sequences, generating a set of per-sample, per-haplotype consensus coding sequences. We ran these sequences through `OrthoFinder` v3.0.1b1 (Emms and Kelly 2019) to generate a multi-species FASTA file for each ortholog. Using these FASTA files, which now contained polymorphic sequences for both species, we used `MACSE` v2.07 to run codon-aware multiple sequence alignments between each ortholog.

The MK test was calculated using the software `MKado` (Rivera-Colón, Rehmann, and Kern 2026). `MKado` takes as input multiple sequence alignments of coding sequences and calculates the ratio of nonsynonymous to synonymous changes for each gene. We ran `MKado` a total of four times. First, we calculated the standard MK test by running `MKado batch` twice, with each species as the ingroup and then the outgroup. Next, we computed the genome-wide *α* using the asymptotic MK method (Messer and Petrov 2013) (`MKado batch -a`) two times switching the ingroup and outgroup species between runs.

### Per-gene divergence and sex-biased mutation rate estimation

From the same single-copy ortholog alignments used for the MK analysis, we extracted pergene counts of nonsynonymous and synonymous fixed differences between *O. bimaculoides* and *O. bimaculatus*, as well as corrected per-site substitution rates *d*_*N*_ and *d*_*S*_ (Jukes– Cantor correction) using the Python package `dnds` v.2.1 (Qalieh 2023). For each ortholog we computed *d*_*N*_ */d*_*S*_, restricting per-gene ratio analyses to the 5,825 orthologs with *d*_*S*_ *>* 0 so that the ratio was defined. To compare evolutionary rates between the Z chromosome (chr17) and autosomes, we tested Z-vs-autosome differences in *d*_*S*_ and *d*_*N*_ with Mann-Whitney *U* tests (one-sided for *d*_*S*_ under the a priori prediction of reduced Z-linked divergence; twosided for *d*_*N*_), and tested the Z-autosome difference in the fraction of orthologs with zero synonymous substitutions using Fisher’s exact test.

We estimated the male-to-female mutation rate ratio using the framework of Miyata et al. (1987): *α*_*m:f*_ = (3*R* − 2)*/*(2 − *R*), where *R* is the ratio of Z-linked to autosomal mean synonymous divergence. To account for sampling uncertainty given the modest sample sizes per species, we computed a bootstrap distribution over *α*_*m:f*_ by repeatedly drawing random subsets of five individuals per species with replacement, recomputing per-gene *d*_*S*_ and the Z-to-autosome ratio at each replicate, and reporting the mean, median, and 2.5/97.5 percentile confidence interval across replicates.

To characterize the tail of orthologs with *d*_*N*_ */d*_*S*_ *>* 1, we compared this set against the remaining *d*_*S*_ *>* 0 orthologs with respect to (i) presence of functional annotation (Fisher’s exact test on annotated vs. unannotated counts), (ii) gene span, estimated as the genomic interval end − start (Mann-Whitney *U* test), and (iii) Z-linkage (Fisher’s exact test on Z-linked vs. autosomal counts). Functional annotations were derived from best reciprocal BLAST hits to *D. melanogaster* (see below); keyword-based enrichment for immunity, defense, and toxin-related annotations was evaluated with a one-sided Fisher’s exact test.

### Inferring selective sweeps and gene set enrichment

In addition to assessing evidence of positive selection on coding sequences, we also inferred selective sweeps for both species using `diploS/HIC` v1.0.6 (Kern and Schrider 2018), where training the classifier was done using simulations with `discoal` v0.1.7 (Kern and Schrider 2016) under neutral, soft and hard sweep conditions with the inclusion of the `SMC++`-inferred demographic history (see above). We simulated in 110 Kb windows, with classification performed in 11 subwindows, resulting in classified sweep regions of 10 Kb. We trained with the same masks as used in other analyses (see methods above). For sweep simulations, we varied selection coefficients from 0.001 to 0.01. We assumed a mutation rate of 3 *×* 10^−9^. After training, we assessed performance by inspecting the confusion matrices for each species and assessed the impact of choosing different classification probability cutoffs.

To assess what genes may have been involved in the inferred selective sweeps, we used `BEDtools` v2.31.1 to find overlaps between sweep regions and coding sequence intervals. We tested for enrichment of gene ontology categories for these genes by using the R package `clusterProfiler` v4.14.0 (Xu et al. 2024). To do so, we found the best reciprocal protein blast hits between *O. bimaculoides* and *Drosophila melanogaster*. We used the full set of best hits as the background as opposed to all annotated *D. melanogaster* genes to look for enrichment of gene ontology categories.

## Data Availability

Sequencing data generated for this study have been deposited to the NCBI Sequence Read Archive (SRA) under accession numbers PRJNA1459481 for *O. bimaculoides* and PRJNA1459341 for *O. bimaculatus*. Processed data and annotations, including VCF files, have been deposited in Zenodo under DOI: 10.5281/zenodo.19697984. Script, code, and other miscellaneous files used in analyses and figure generation are available on GitHub at https://github.com/kr-colab/Two-spot-Octopus-Population-Genomics.

## Acknowledgments

We thank the University of Oregon Genomics & Cell Characterization Core Facility (GC3F) for their help with DNA extraction, library preparation, and sequencing. We also thank the members of the Kern-Ralph lab for their feedback on this project and manuscript. This work was supported in part by NIH awards R35148253 and R01HG010774 to A.D.K.

## Author Contributions

A.D.K. oversaw the project. G.C.C. generated and filtered VCFs, calculated population genetics statistics, and estimated demographic history. S.T. estimated recombination rates and inferred selective sweeps. S.T., S.T.S., and A.D.K. advised on data curation and filtering. G.C.C. prepared the initial draft of this manuscript. All authors contributed to the writing and editing of the final manuscript.

