## Supplementary materials for "Diversity and divergence of two sympatric, sibling octopus species"

Table S1: Raw sequencing statistics for re-sequencing dataset of octopus. Total reads and percent mapped were calculated by the `SNPArcher` pipeline. Depth per individual was calculated by `mosdepth`. Datasets that were published previously were used as coverage data in Coffing et al. (2025).

| Sample | Sex | Tissue | Total Reads | % Mapped | Depth | Previously published? |
|---|---|---|---:|---:|---:|---|
| Oblat_F_2 | F | Optic lobe | 99.14% | 669M | 35x | Yes |
| Oblat_F_5 | F | Optic lobe | 99.39% | 738M | 35x | No |
| Oblat_F_6 | F | Optic lobe | 99.04% | 719M | 40x | No |
| Oblat_F_7 | M | Optic lobe | 99.00% | 703M | 39x | No |
| Oblat_F_8 | M | Optic lobe | 99.42% | 425M | 21x | No |
| Oblat_F_9 | F | Optic lobe | 99.23% | 920M | 49x | No |
| Oblat_F_10 | F | Optic lobe | 99.05% | 764M | 42x | Yes |
| Oblat_F_11 | F | Optic lobe | 99.07% | 706M | 36x | No |
| Oblat_M_01 | M | Optic lobe | 98.78% | 664M | 34x | No |
| Oblat_M_3 | M | Optic lobe | 99.24% | 664M | 34x | Yes |
| Oblat_M_4 | M | Optic lobe | 99.17% | 779M | 42x | Yes |
| Oblat_M_12 | M | Optic lobe | 99.02% | 664M | 33x | No |
| Oblat_M_13 | M | Optic lobe | 99.15% | 664M | 48x | No |
| Obmac_F_2 | F | Optic lobe | 99.19% | 1.24B | 53x | No |
| Obmac_F_4 | F | Arm, gill | 99.25% | 976M | 37x | No |
| Obmac_F_10 | F | Gill | 99.21% | 1.13B | 44x | No |
| Obmac_M_3 | M | Optic lobe | 99.20% | 1.05B | 43x | No |
| Obmac_M_8 | M | Arm, gill, optic lobe | 99.11% | 913M | 39x | No |
| Obmac_M_14 | M | Testes | 99.25% | 923M | 37x | No |

### Supplementary files

`bimaculoides_drosophila_blasttable.txt` – Best reciprocal blast hits between *O. bimaculoides* and *D. melanogaster* and forward and reverse e-values.

`sweep_windows_filtered_0.5prob.bed` – Bed file of selective sweep regions found separately in both species called with `diploS/HIC` at a probability cutoff of 0.5.

`shared_sweep_genes_0.5.txt` - Genes overlapping selective sweep regions (soft or hard) shared between *O. bimaculatus* and *O. bimaculoides* at a classification probability threshold of 0.5.

`bimaculatus_sweep_genes_0.95.txt` - Genes overlapping selective sweep regions (soft or hard) in *O. bimaculatus* at a classification probability threshold of 0.95.

`bimaculatus_mk_testable.tsv` - List of all genes proposed for McDonald-Kreitman tests in *O. bimaculatus* with neutrality indices and p-values.

`bimaculoides_mk_testable.tsv` - List of all genes proposed for McDonald-Kreitman tests in *O. bimaculoides* with neutrality indices and p-values.

`dn_ds_table.txt` - Per-gene nonsynonymous and synonymous substitution counts, polymorphism counts, and Jukes-Cantor-corrected $d_N$, $d_S$, and $d_N/d_S$ values for the 12,827 single-copy orthologs used in the per-gene divergence analysis.

`go_enrichment_results_mk_and_sweeps.csv` - Significantly enriched gene ontology categories output by `clusterProfiler`'s `enrichGO` function for genes significant in the McDonald-Kreitman tests and genes overlapping selective sweep regions. Final columns were added that indicate species and which analysis the enrichment result corresponds to (mk_genes or sweep_genes).

Table S2: Summary of life history strategies of the Two-spot octopus species. Adapted from Domínguez-Contreras, Munguia-Vega, Ceballos-Vázquez, Arellano-Martínez, et al. (2018).

| Trait | *O. bimaculoides* | *O. bimaculatus* |
|---|---|---|
| Geographic distribution | California, USA to Bahía San Quintín, BC, Mexico | California, USA to Bahía Vizcaíno, BCS, including Gulf of California |
| Reproductive period | Santa Barbara, CA: Dec–May; San Quintín, BCP: Oct–Jan | Pacific coast BCP: Jan–Jun; Gulf of California: Jun–Sep |
| Fecundity | 137–780 eggs in festoons | >20,000 clutch eggs |
| Egg size (length) | 10–12 mm (range 9.5–16 mm) | 4–7 mm (range 1.8–4 mm) |
| Planktonic larval duration | Absent; direct development to benthic juveniles | 2–3 months (60–90 days) |
| Size at sexual maturity | Males: 55 mm ML; Females: 110 mm ML | Males: 124.5 mm ML; Females: 147.0 mm ML |
| Lifespan | 1.0–1.5 years | 1.5–2.0 years |

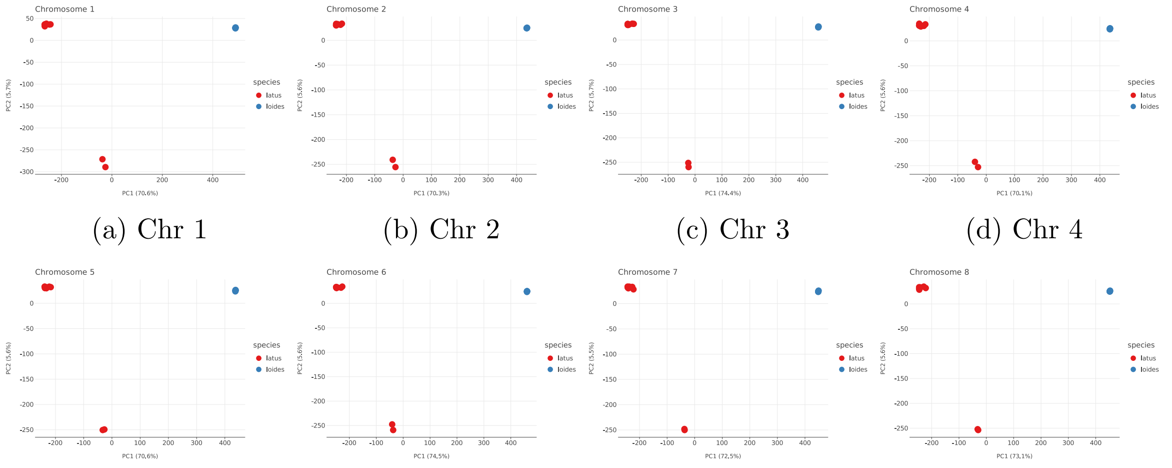


(a) Chr 1 (b) Chr 2 (c) Chr 3 (d) Chr 4

(e) Chr 5 (f) Chr 6 (g) Chr 7 (h) Chr 8

Figure S1: PCA of chromosomes 1–8 of *O. bimaculoides* and *O. bimaculatus* individuals.

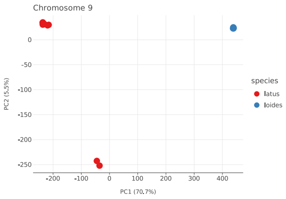


(a) Chr 9

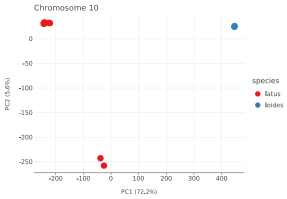


(b) Chr 10

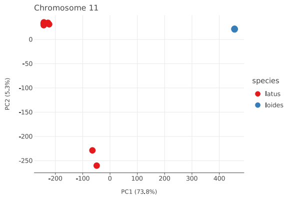


(c) Chr 11

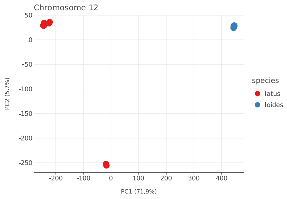


(d) Chr 12

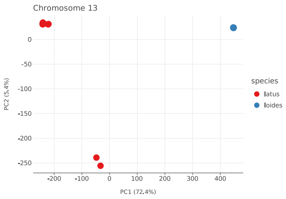


(e) Chr 13

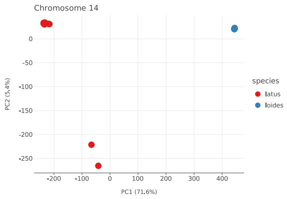


(f) Chr 14

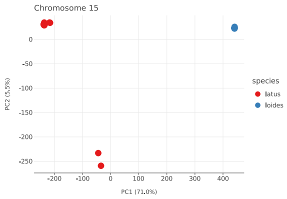


(g) Chr 15

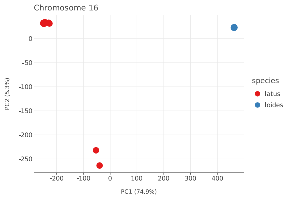


(h) Chr 16

Figure S2: PCA of chromosomes 9–16 of *O. bimaculoides* and *O. bimaculatus* individuals.

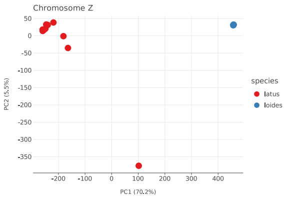


(a) Chr Z

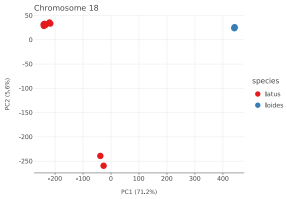


(b) Chr 18

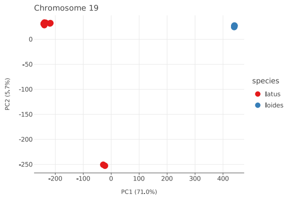


(c) Chr 19

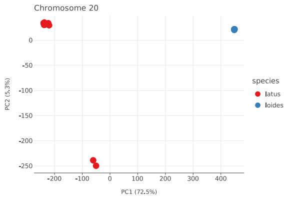


(d) Chr 20

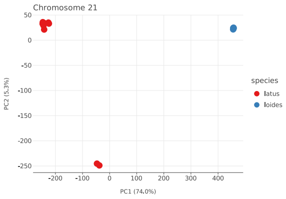


(e) Chr 21

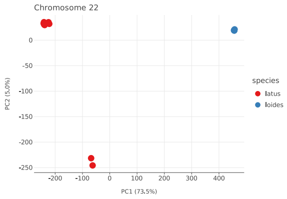


(f) Chr 22

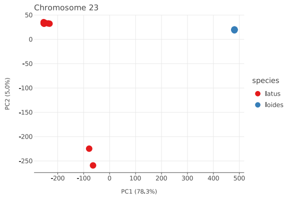


(g) Chr 23

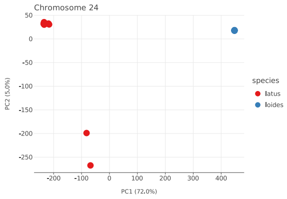


(h) Chr 24

Figure S3: PCA of chromosomes Z and 18–24 of *O. bimaculoides* and *O. bimaculatus* individuals.

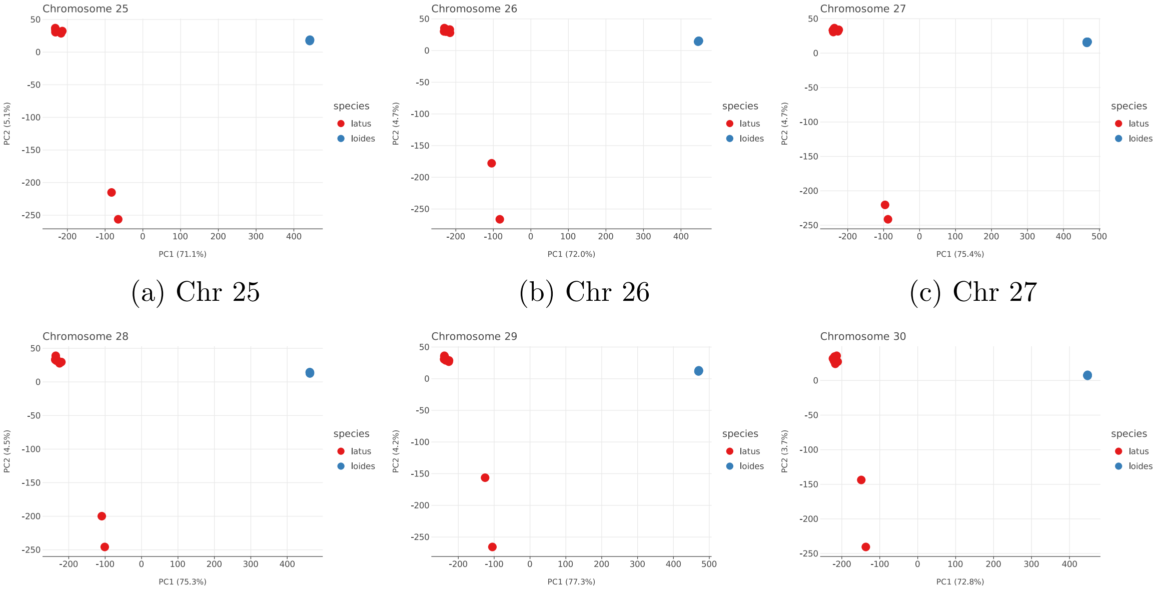


(a) Chr 25 (b) Chr 26 (c) Chr 27

(d) Chr 28 (e) Chr 29 (f) Chr 30

Figure S4: PCA of chromosomes 25–30 of *O. bimaculoides* and *O. bimaculatus* individuals.

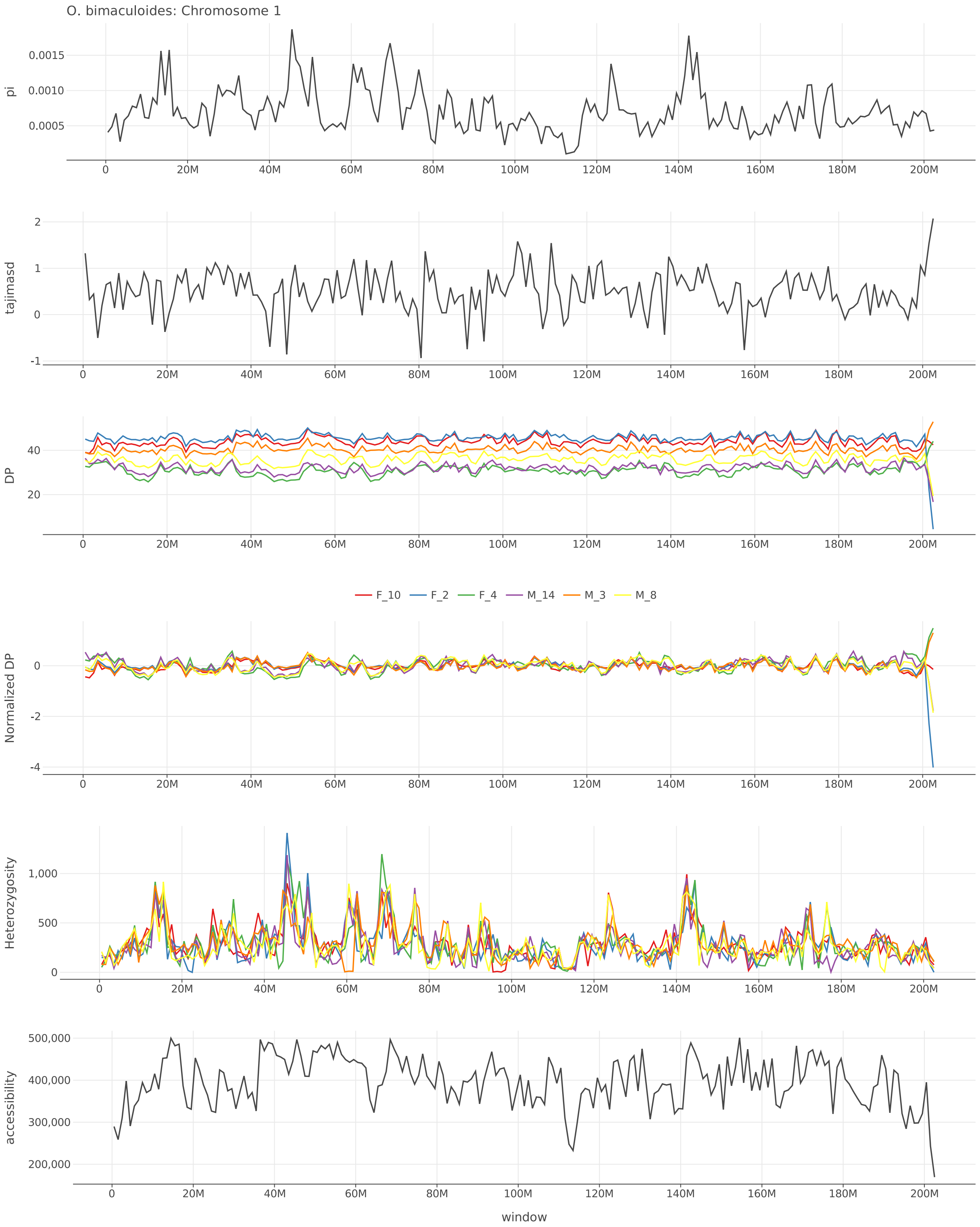


Figure S5: $\pi$, Tajima's $D$, depth, normalized depth, heterozygosity, and accessibility across chromosome 1 of *O. bimaculoides*.

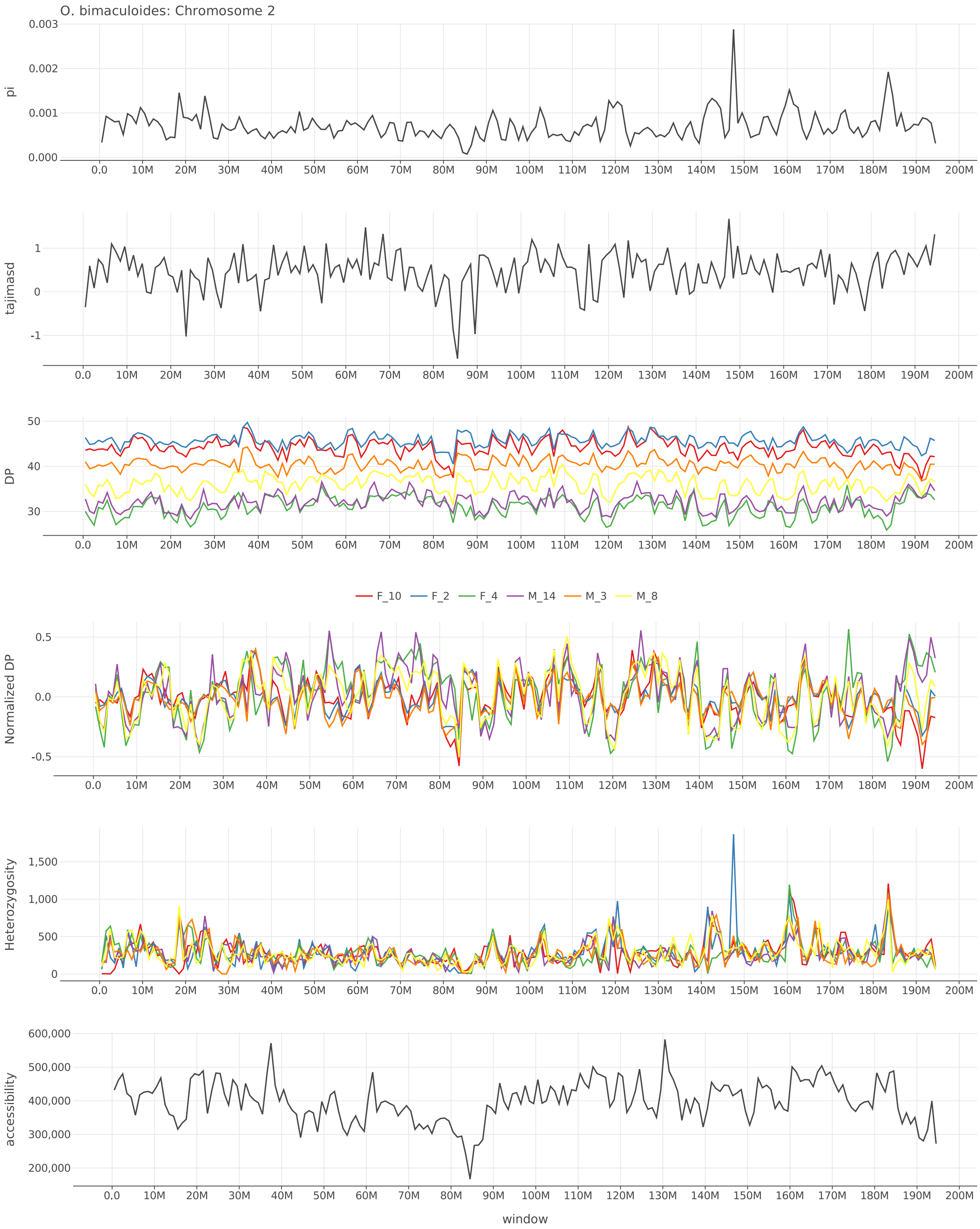


Figure S6: $\pi$, Tajima's $D$, depth, normalized depth, heterozygosity, and accessibility across chromosome 2 of *O. bimaculoides*.

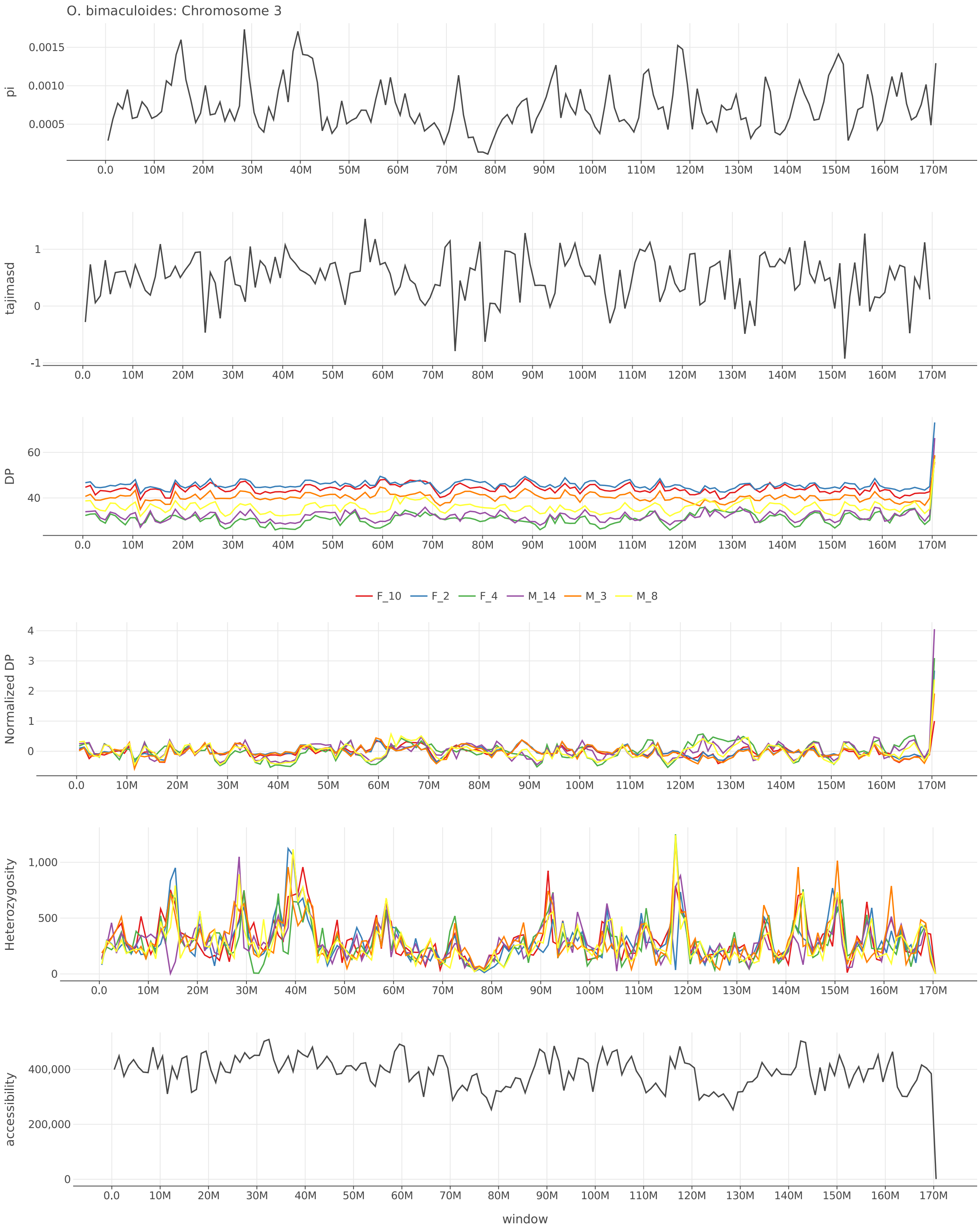


Figure S7: $\pi$, Tajima's $D$, depth, normalized depth, heterozygosity, and accessibility across chromosome 3 of *O. bimaculoides*.

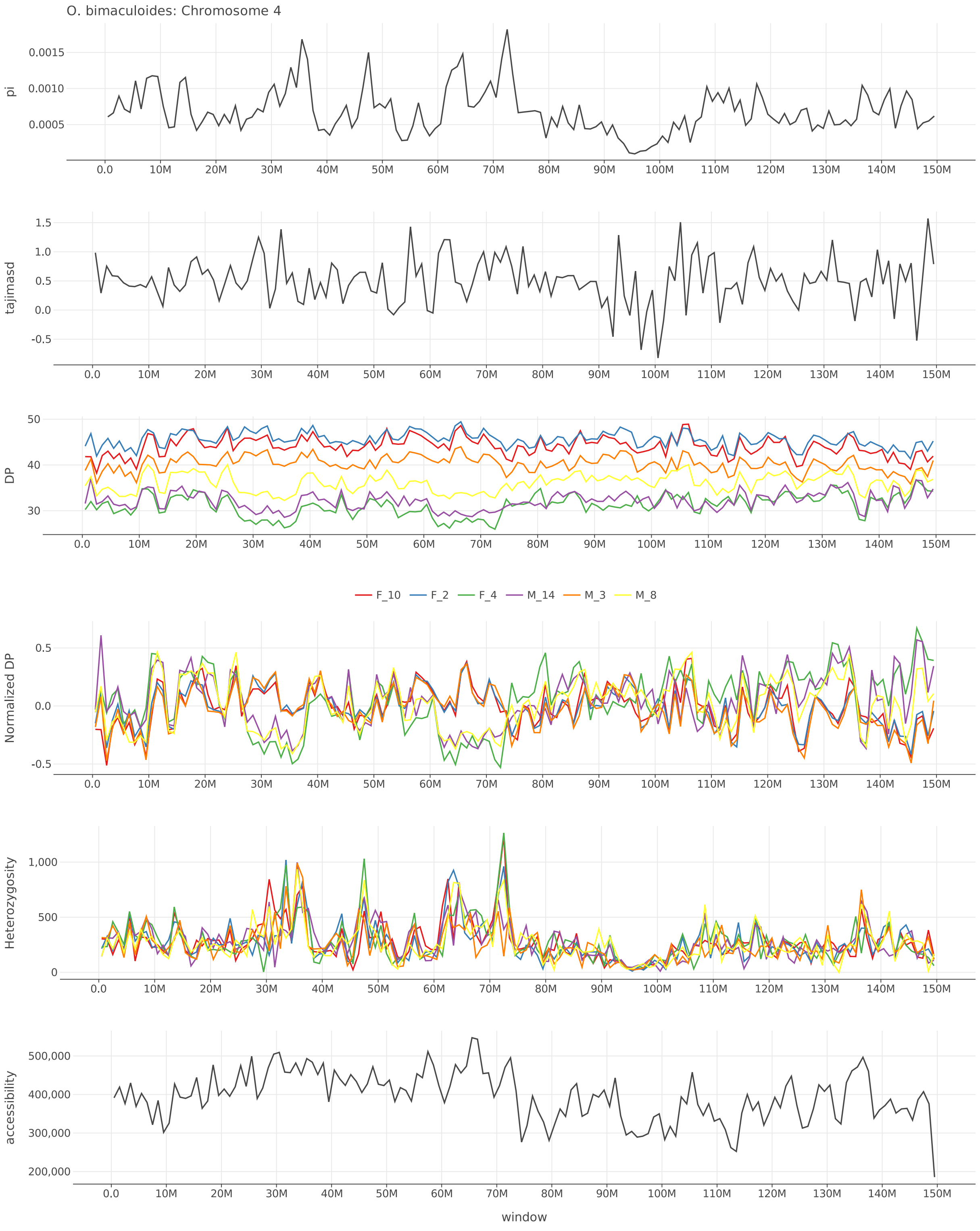


Figure S8: $\pi$, Tajima's $D$, depth, normalized depth, heterozygosity, and accessibility across chromosome 4 of *O. bimaculoides*.

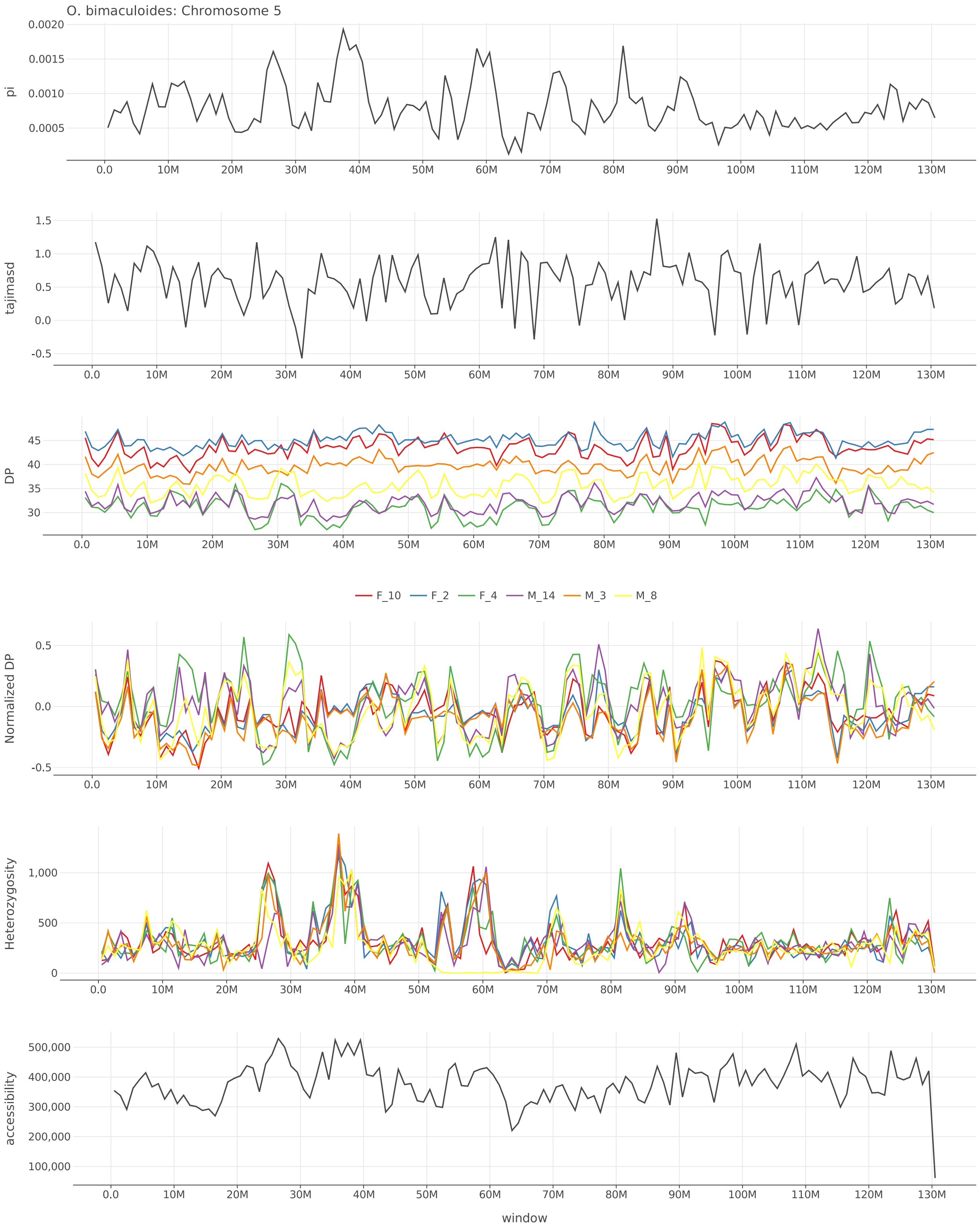


Figure S9: $\pi$, Tajima's $D$, depth, normalized depth, heterozygosity, and accessibility across chromosome 5 of *O. bimaculoides*.

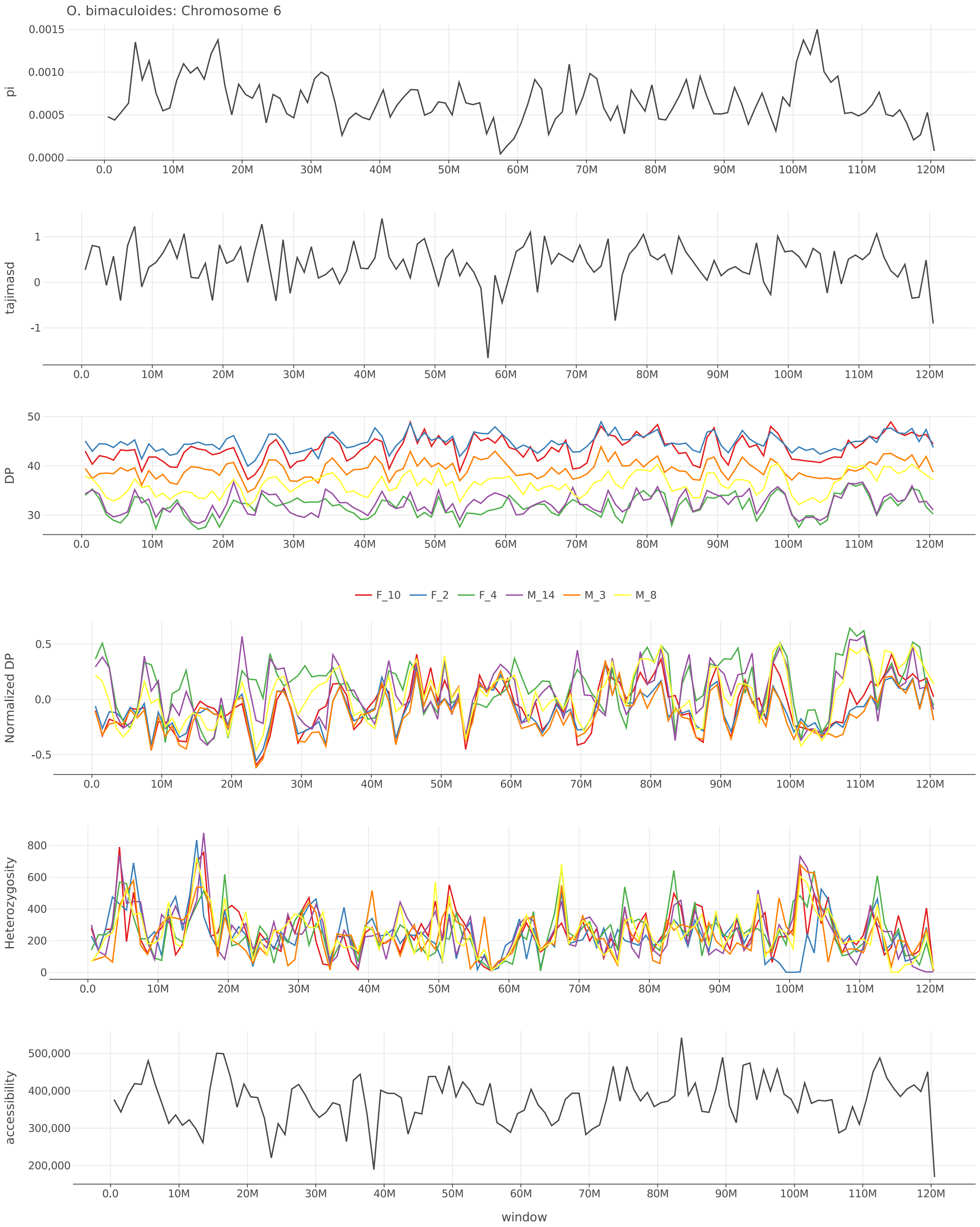


Figure S10: $\pi$, Tajima's $D$, depth, normalized depth, heterozygosity, and accessibility across chromosome 6 of *O. bimaculoides*.

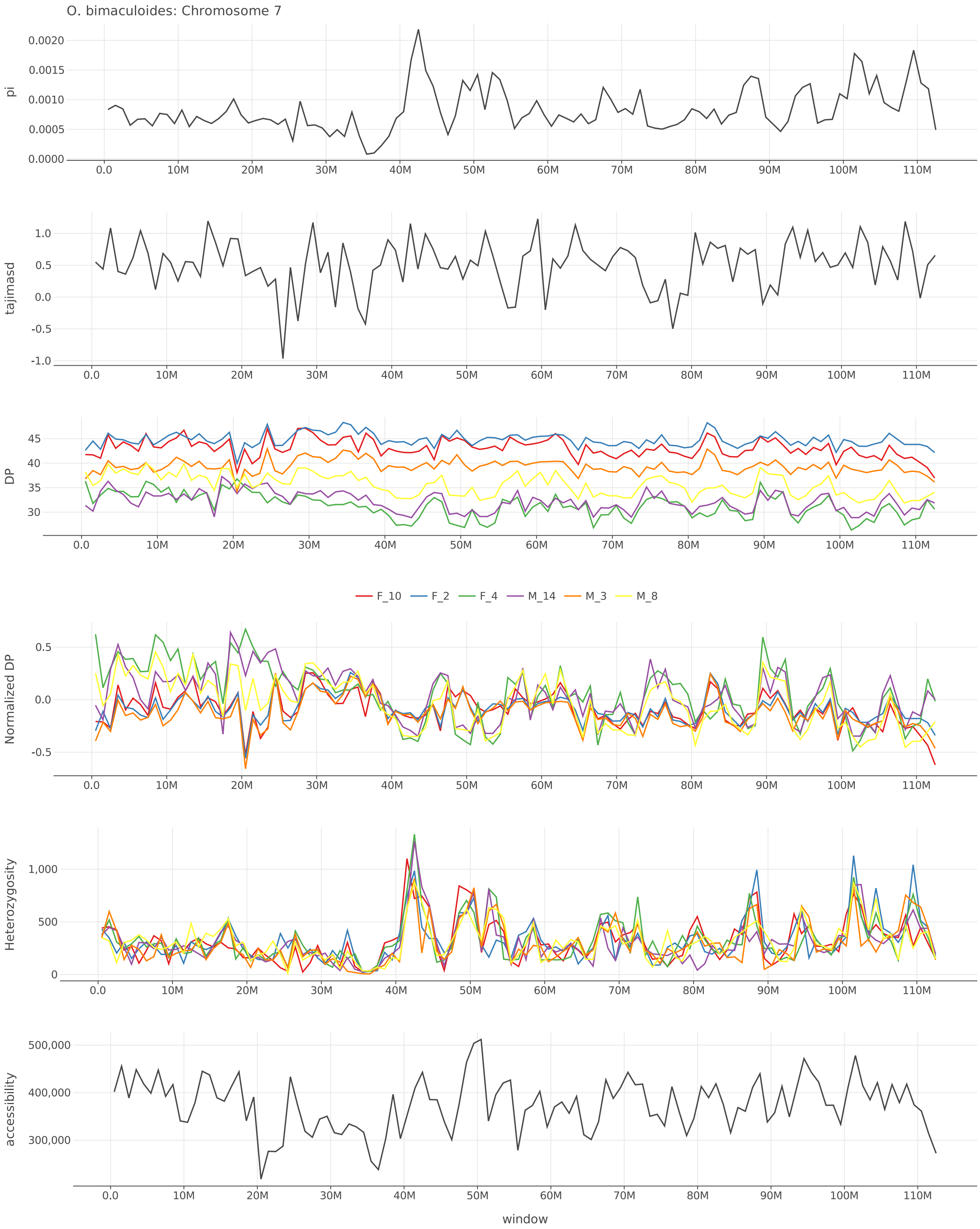


Figure S11: $\pi$, Tajima's $D$, depth, normalized depth, heterozygosity, and accessibility across chromosome 7 of *O. bimaculoides*.

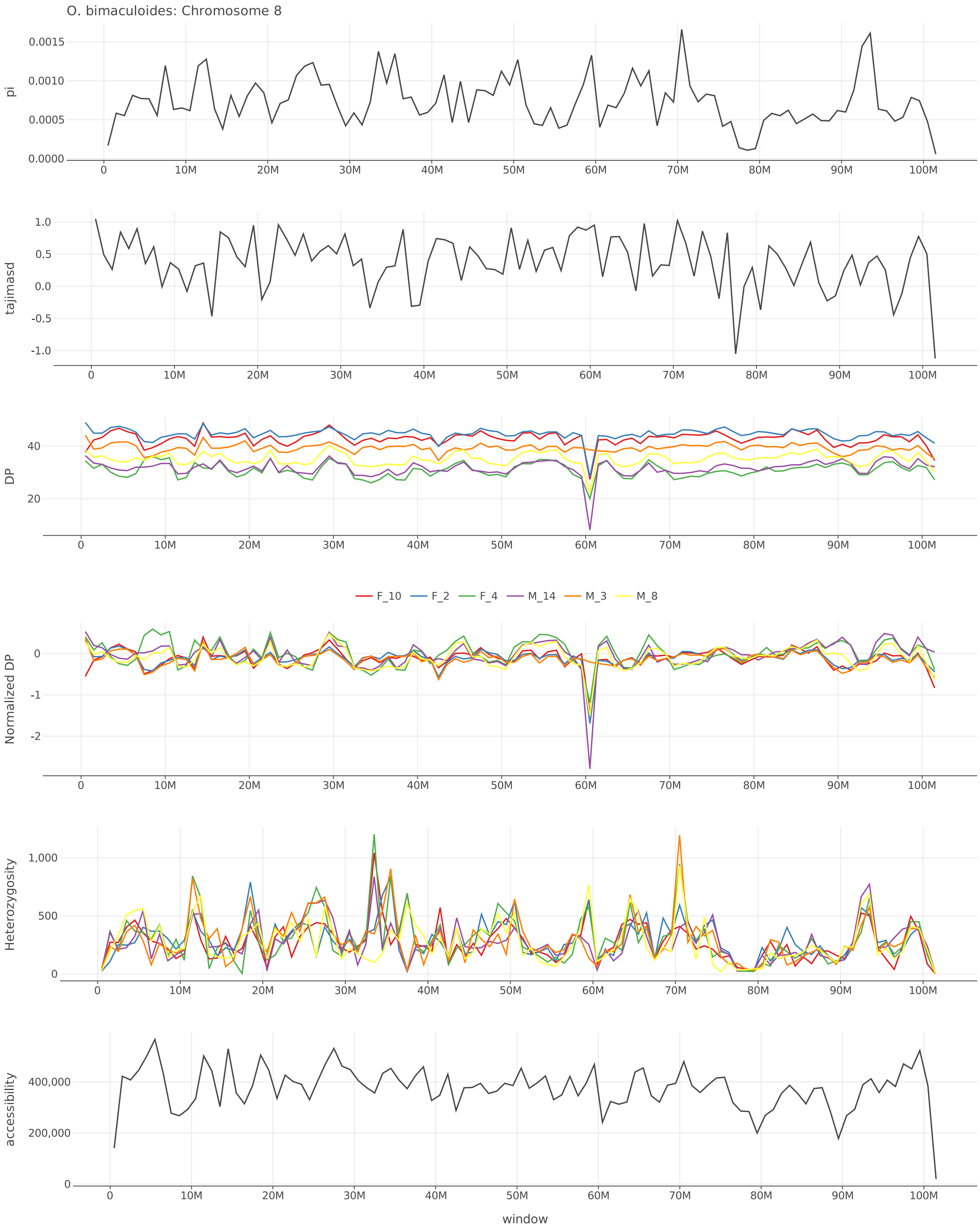


Figure S12: $\pi$, Tajima's $D$, depth, normalized depth, heterozygosity, and accessibility across chromosome 8 of *O. bimaculoides*.

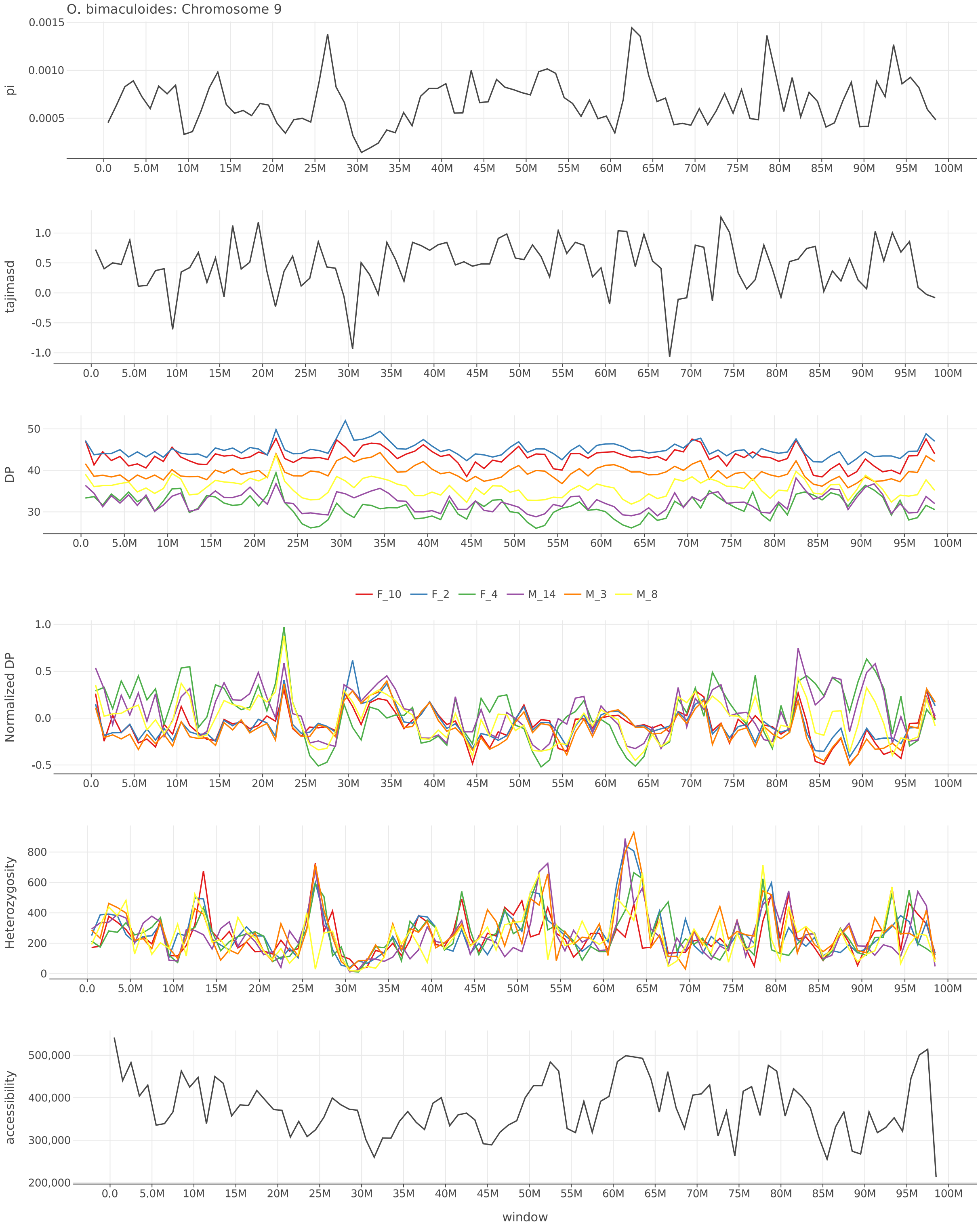


Figure S13: $\pi$, Tajima's $D$, depth, normalized depth, heterozygosity, and accessibility across chromosome 9 of *O. bimaculoides*.

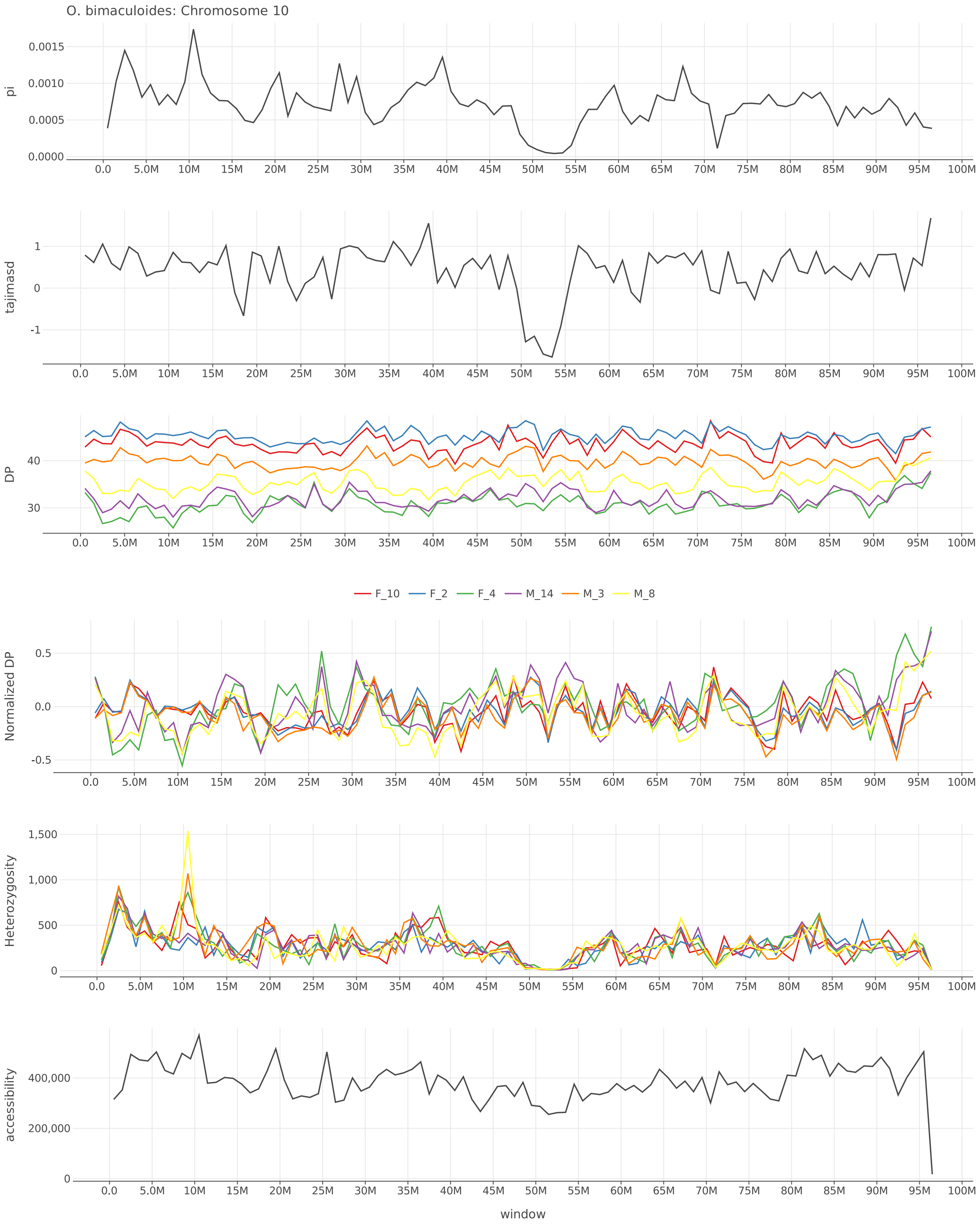


Figure S14: $\pi$, Tajima's $D$, depth, normalized depth, heterozygosity, and accessibility across chromosome 10 of *O. bimaculoides*.

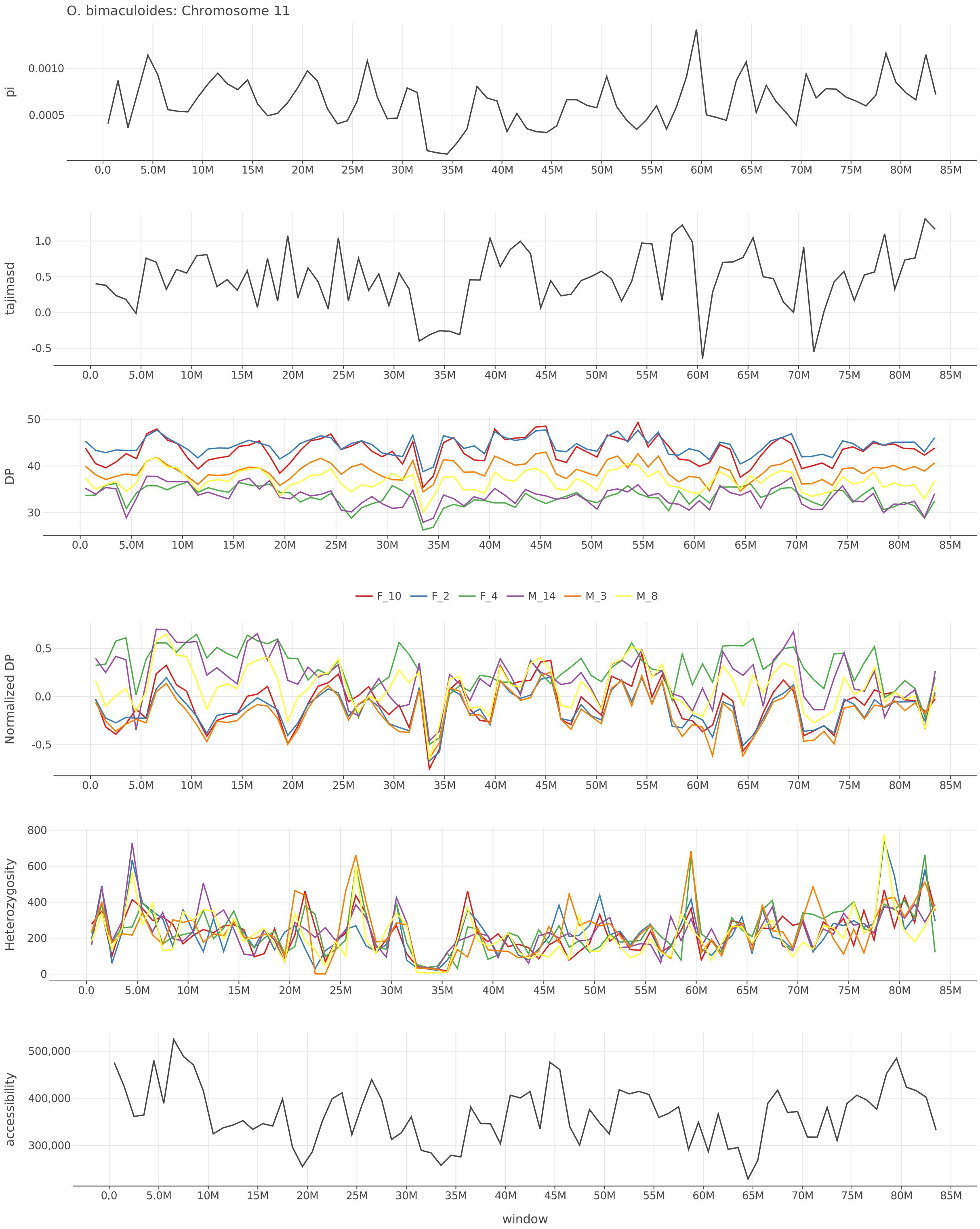


Figure S15: $\pi$, Tajima's $D$, depth, normalized depth, heterozygosity, and accessibility across chromosome 11 of *O. bimaculoides*.

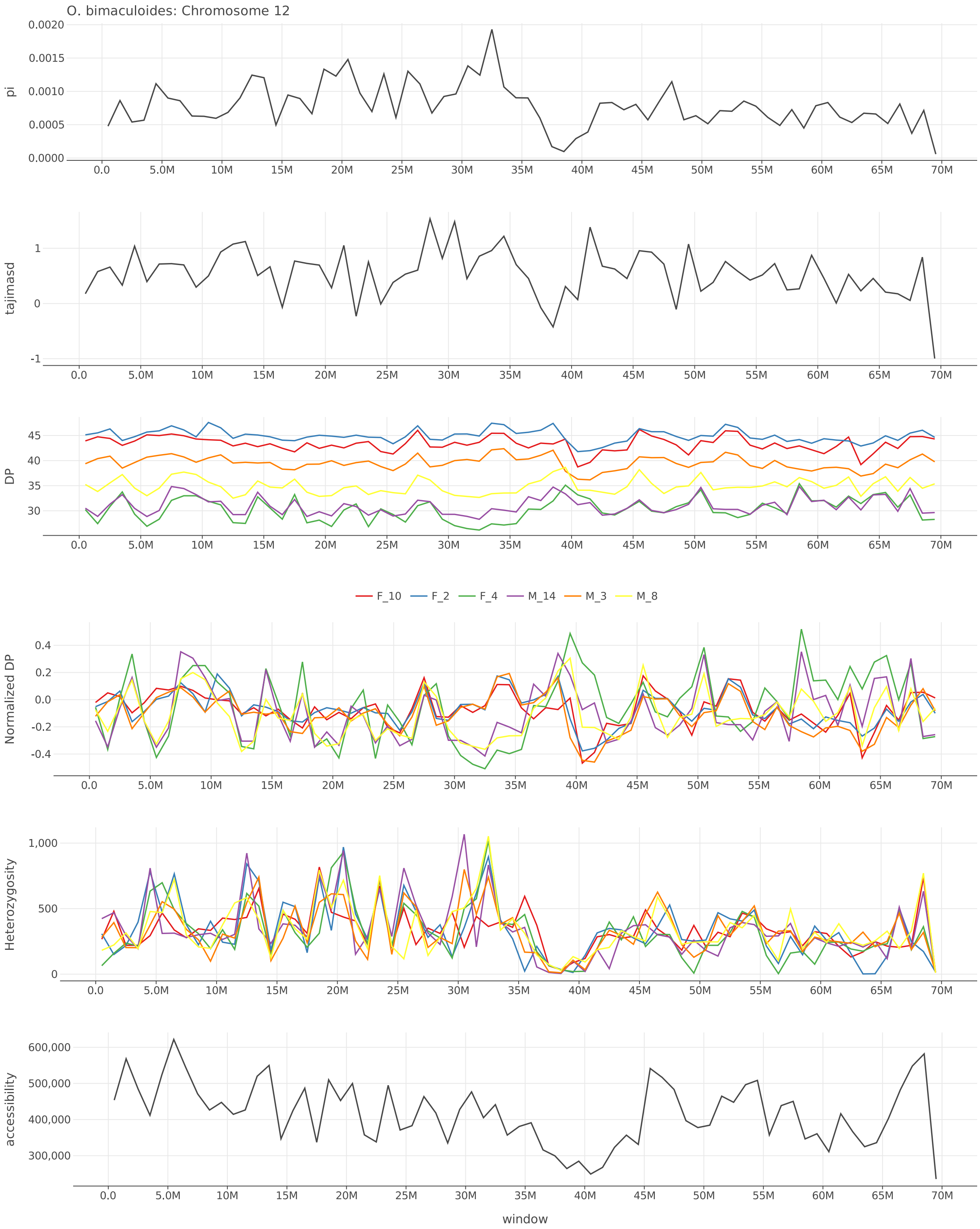


Figure S16: $\pi$, Tajima's $D$, depth, normalized depth, heterozygosity, and accessibility across chromosome 12 of *O. bimaculoides*.

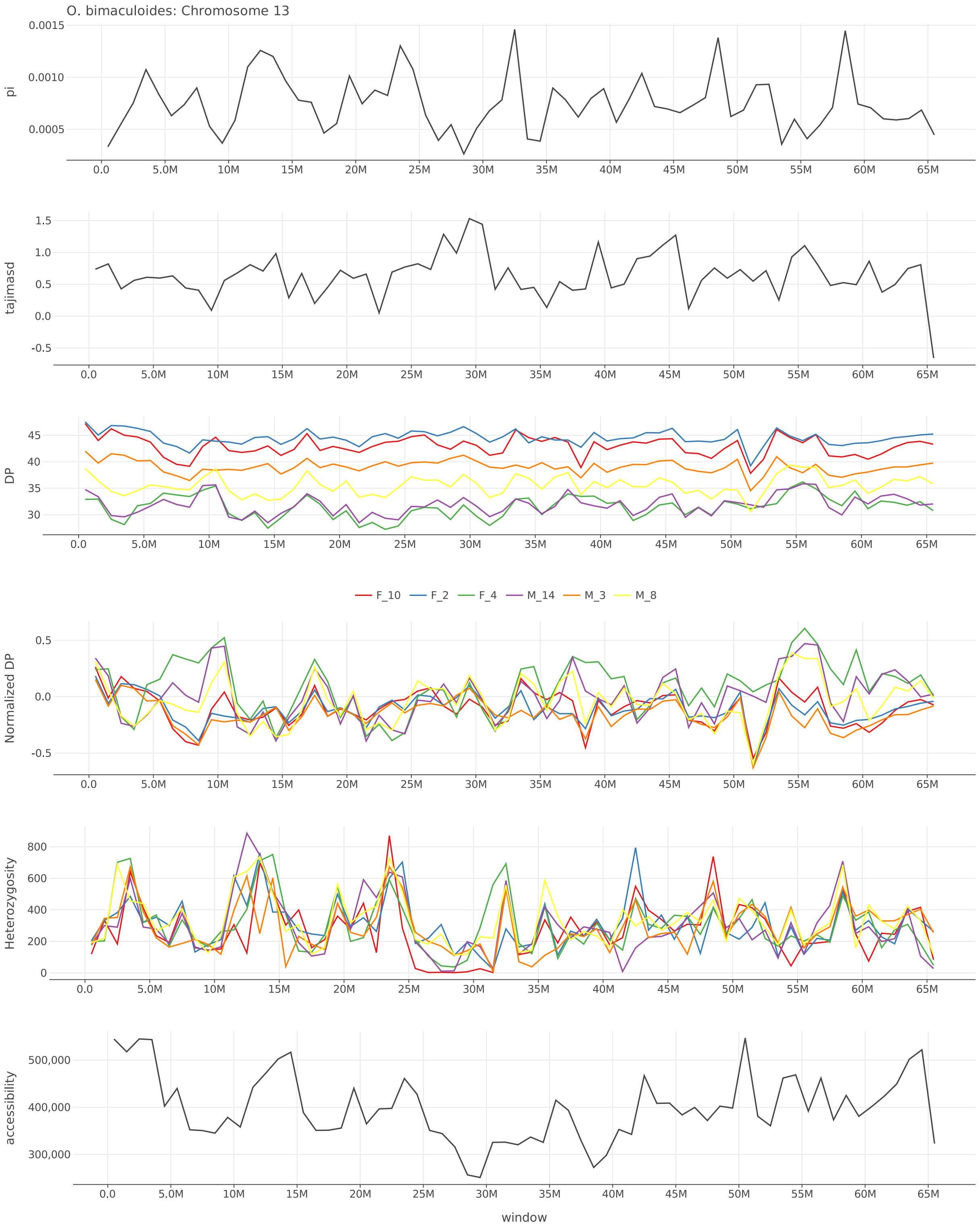


Figure S17: $\pi$, Tajima's $D$, depth, normalized depth, heterozygosity, and accessibility across chromosome 13 of *O. bimaculoides*.

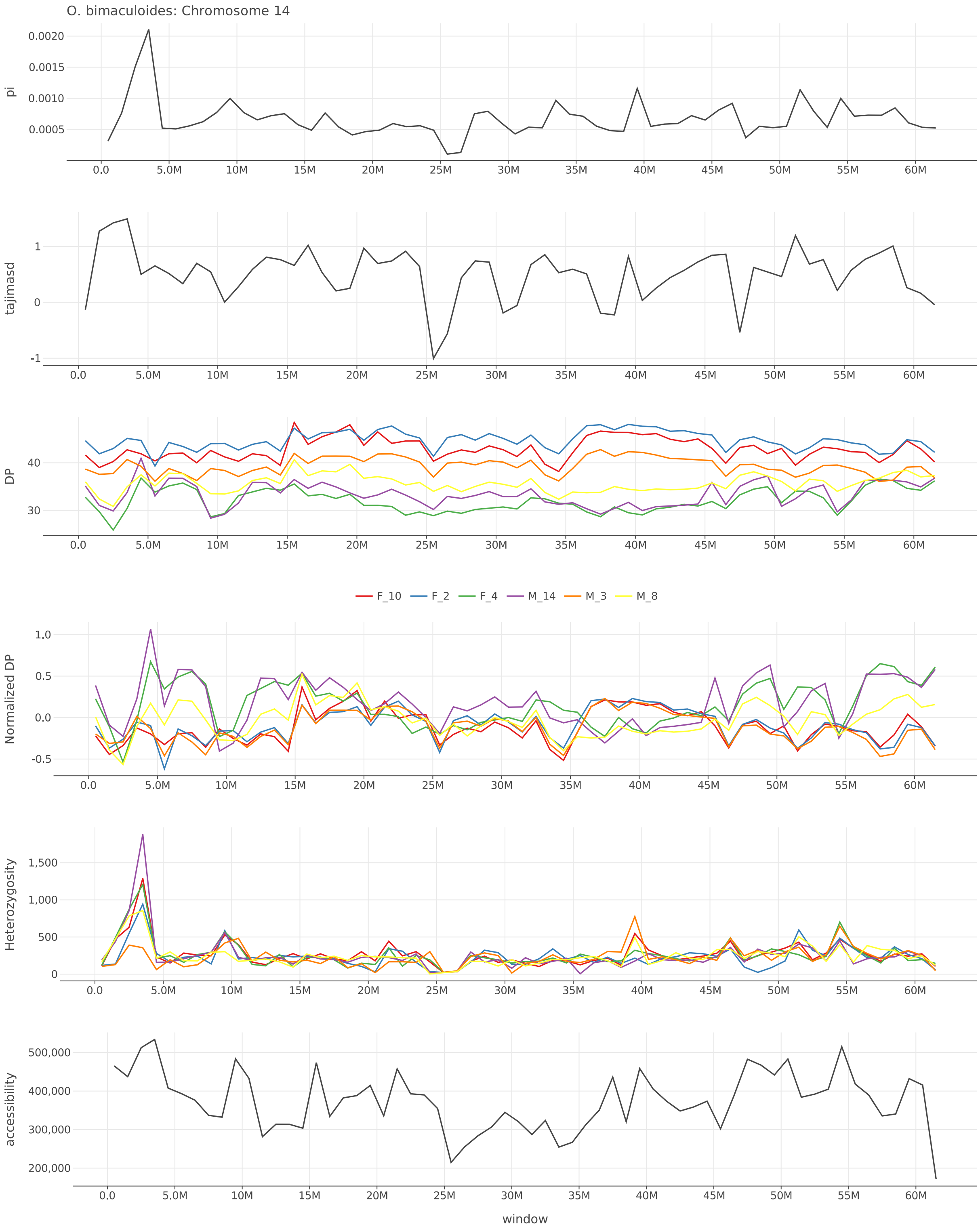


Figure S18: $\pi$, Tajima's $D$, depth, normalized depth, heterozygosity, and accessibility across chromosome 14 of *O. bimaculoides*.

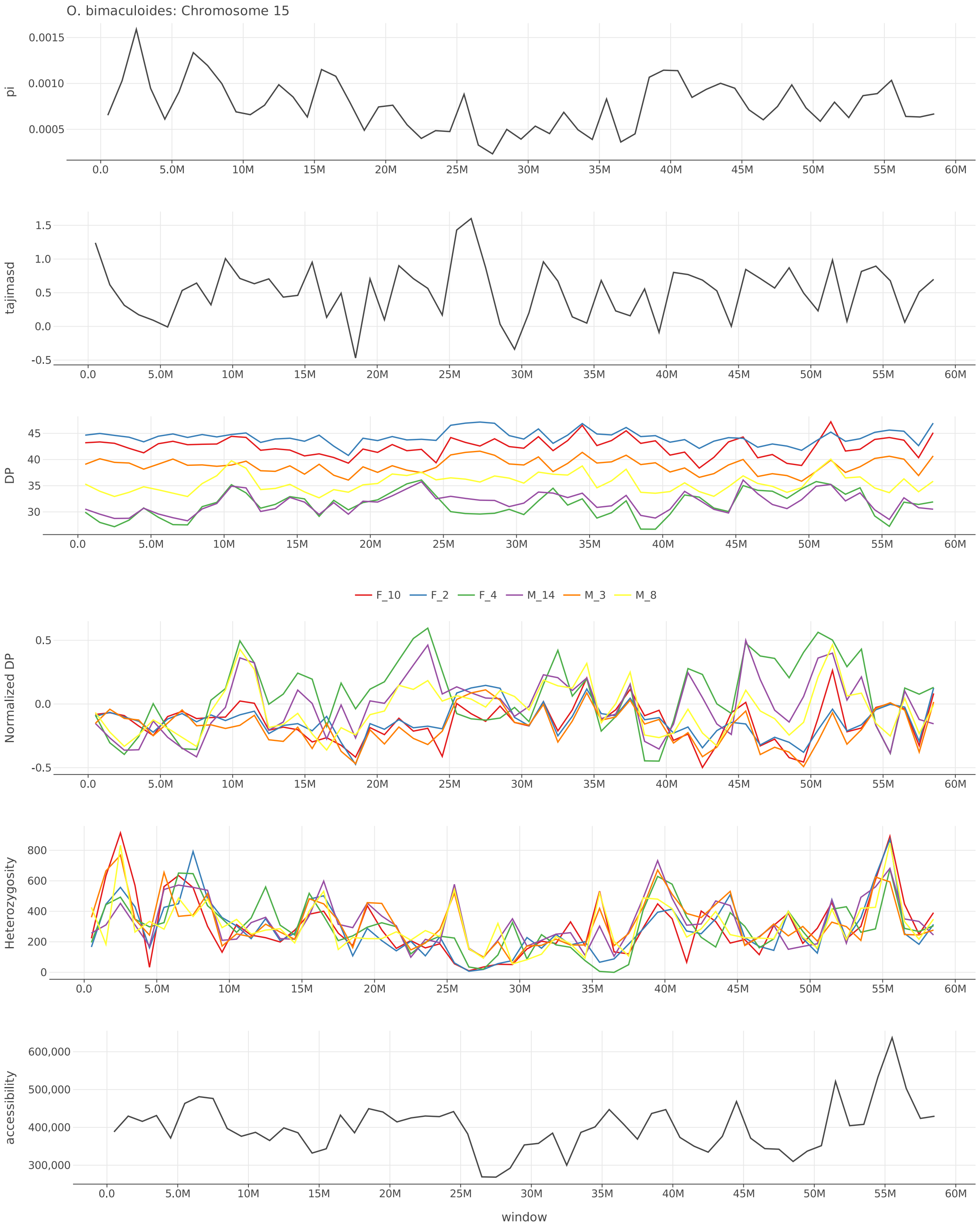


Figure S19: $\pi$, Tajima's $D$, depth, normalized depth, heterozygosity, and accessibility across chromosome 15 of *O. bimaculoides*.

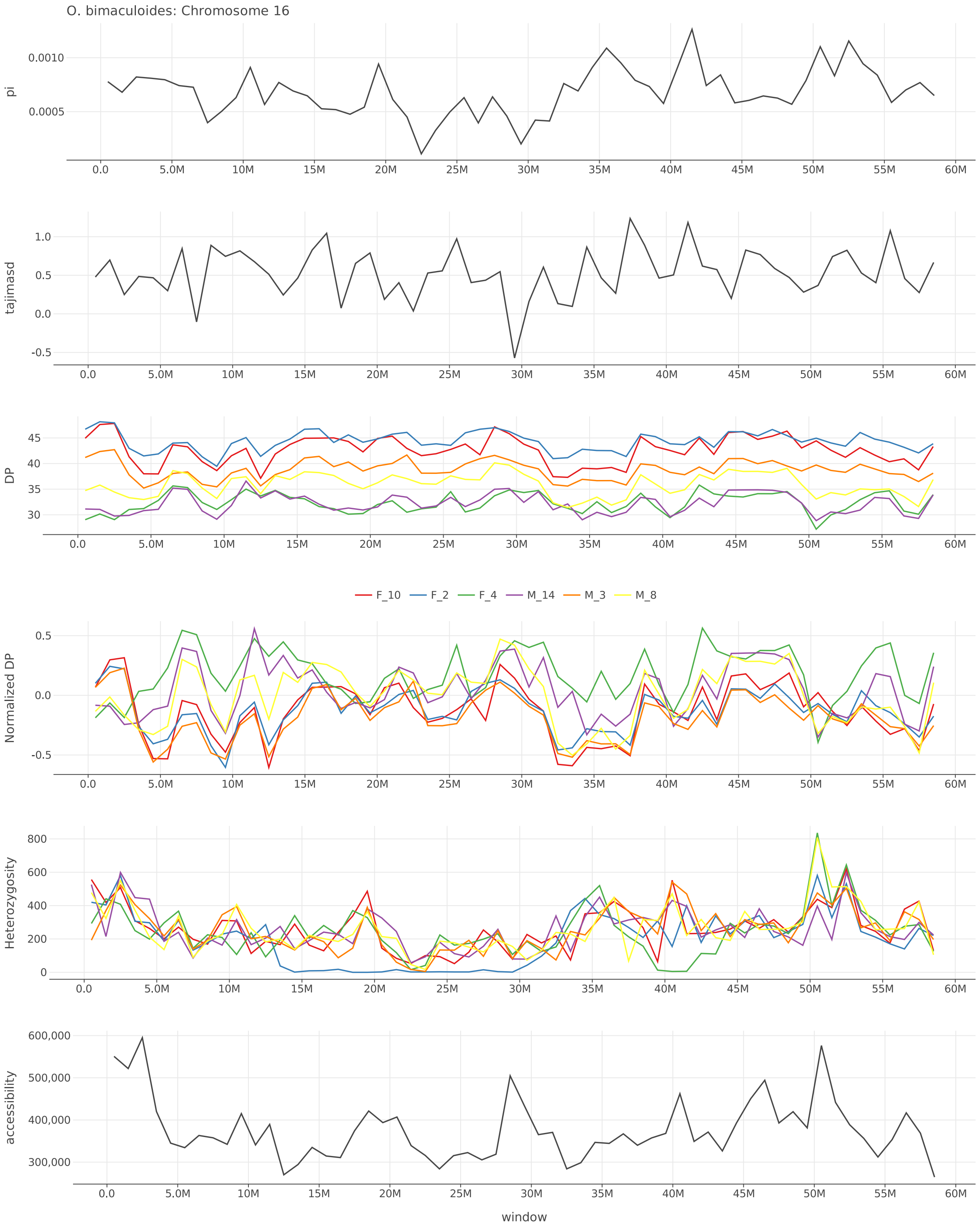


Figure S20: $\pi$, Tajima's $D$, depth, normalized depth, heterozygosity, and accessibility across chromosome 16 of *O. bimaculoides*.

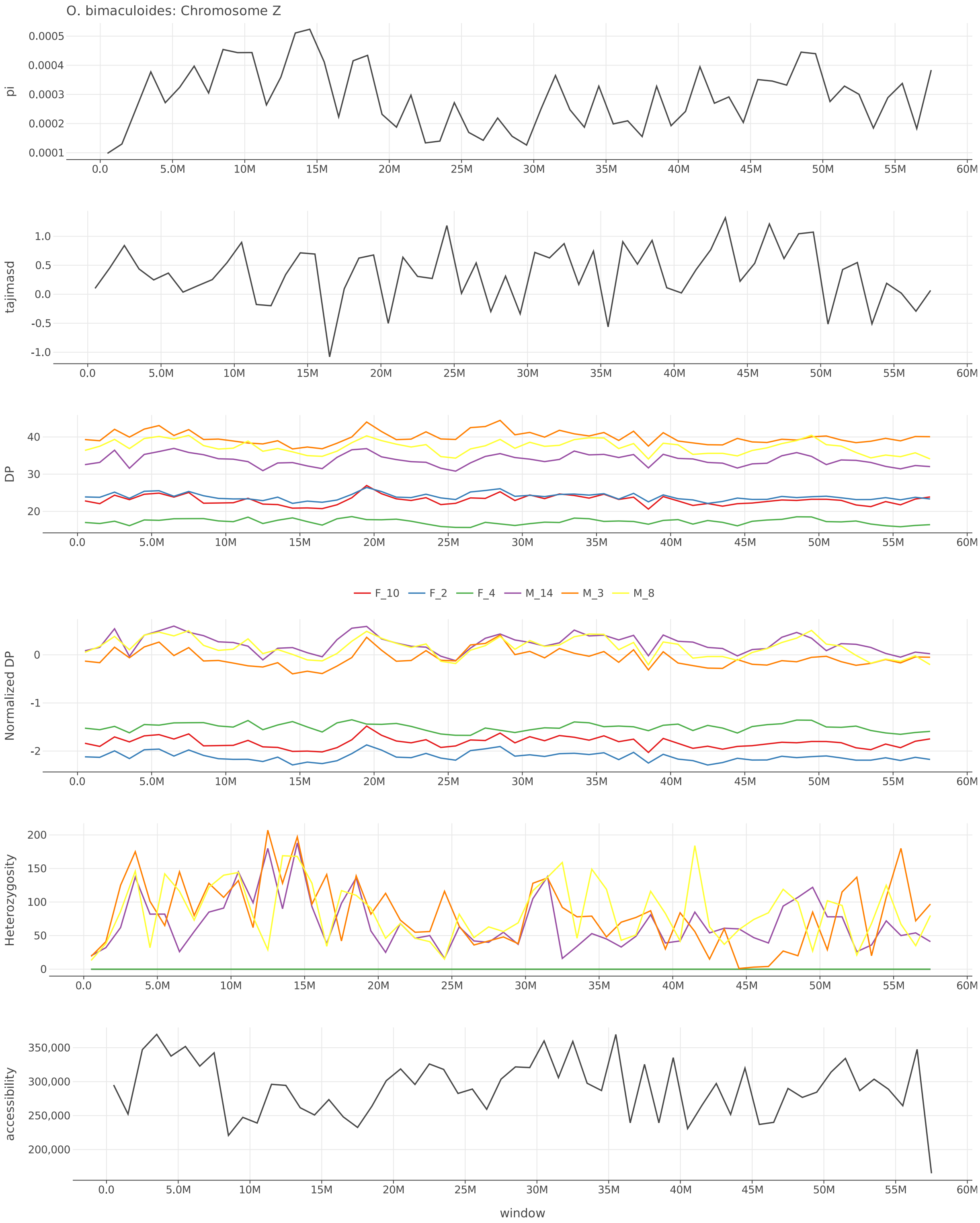


Figure S21: $\pi$, Tajima's $D$, depth, normalized depth, heterozygosity, and accessibility across chromosome Z of *O. bimaculoides*.

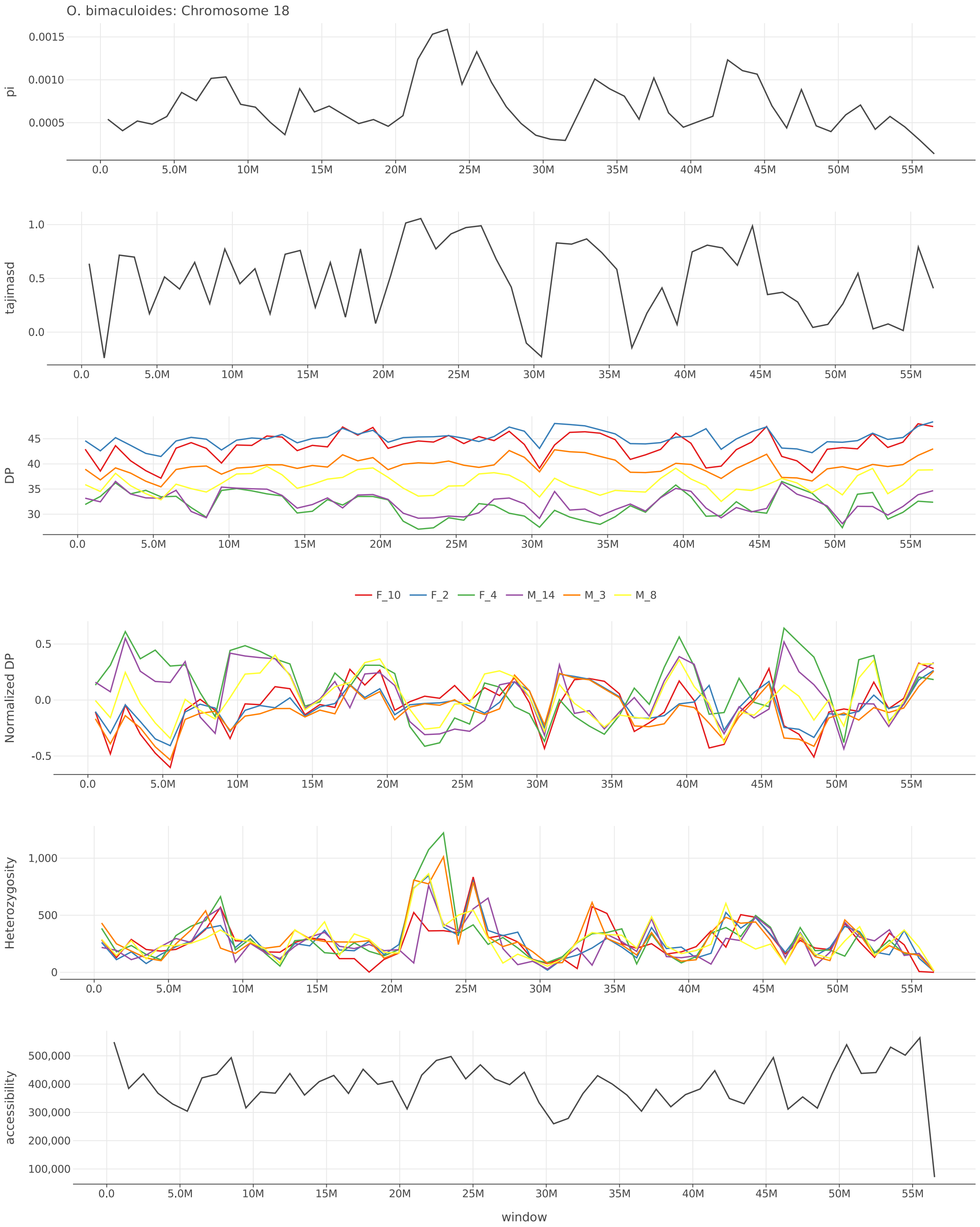


Figure S22: $\pi$, Tajima's $D$, depth, normalized depth, heterozygosity, and accessibility across chromosome 18 of *O. bimaculoides*.

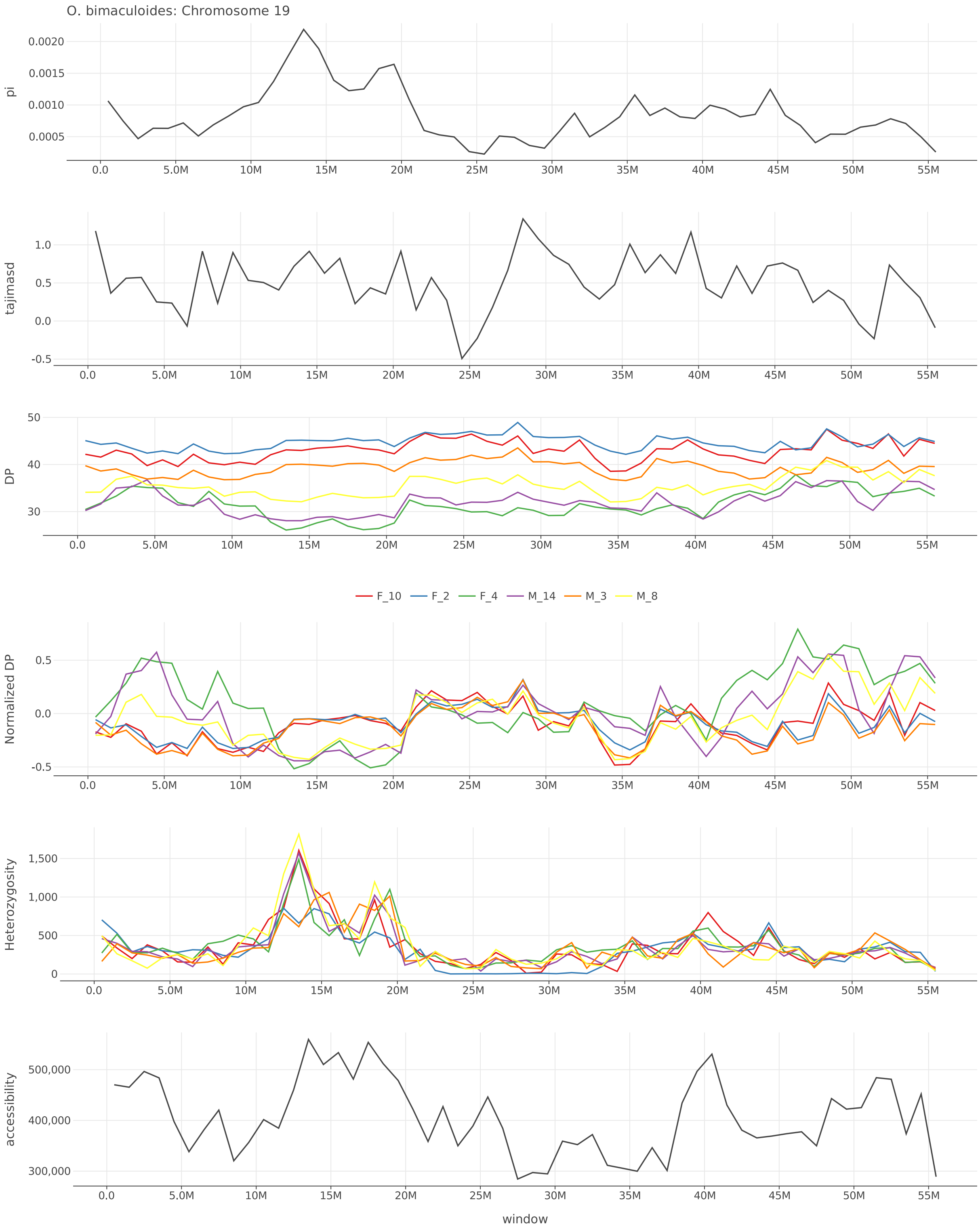


Figure S23: $\pi$, Tajima's $D$, depth, normalized depth, heterozygosity, and accessibility across chromosome 19 of *O. bimaculoides*.

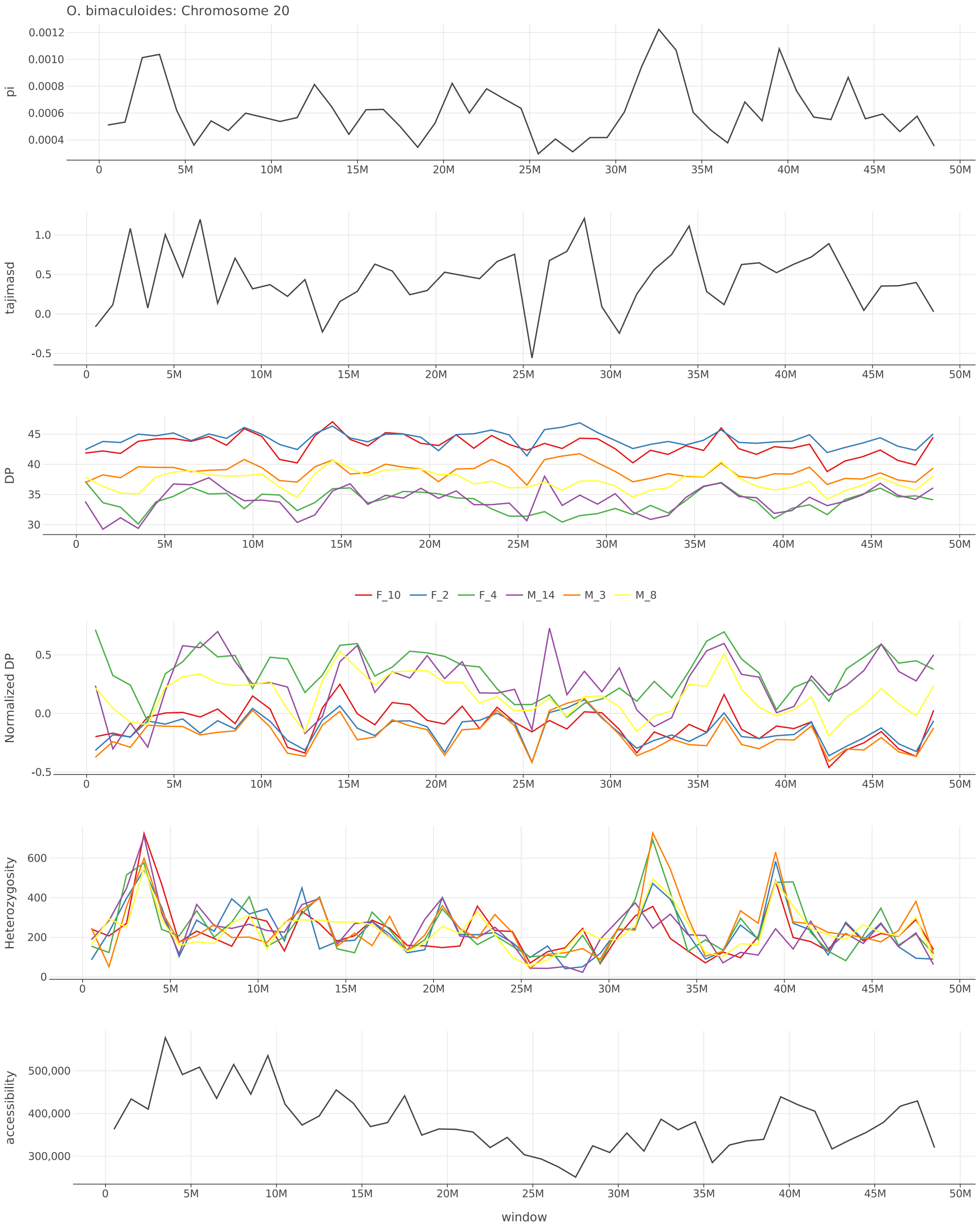


Figure S24: $\pi$, Tajima's $D$, depth, normalized depth, heterozygosity, and accessibility across chromosome 20 of *O. bimaculoides*.

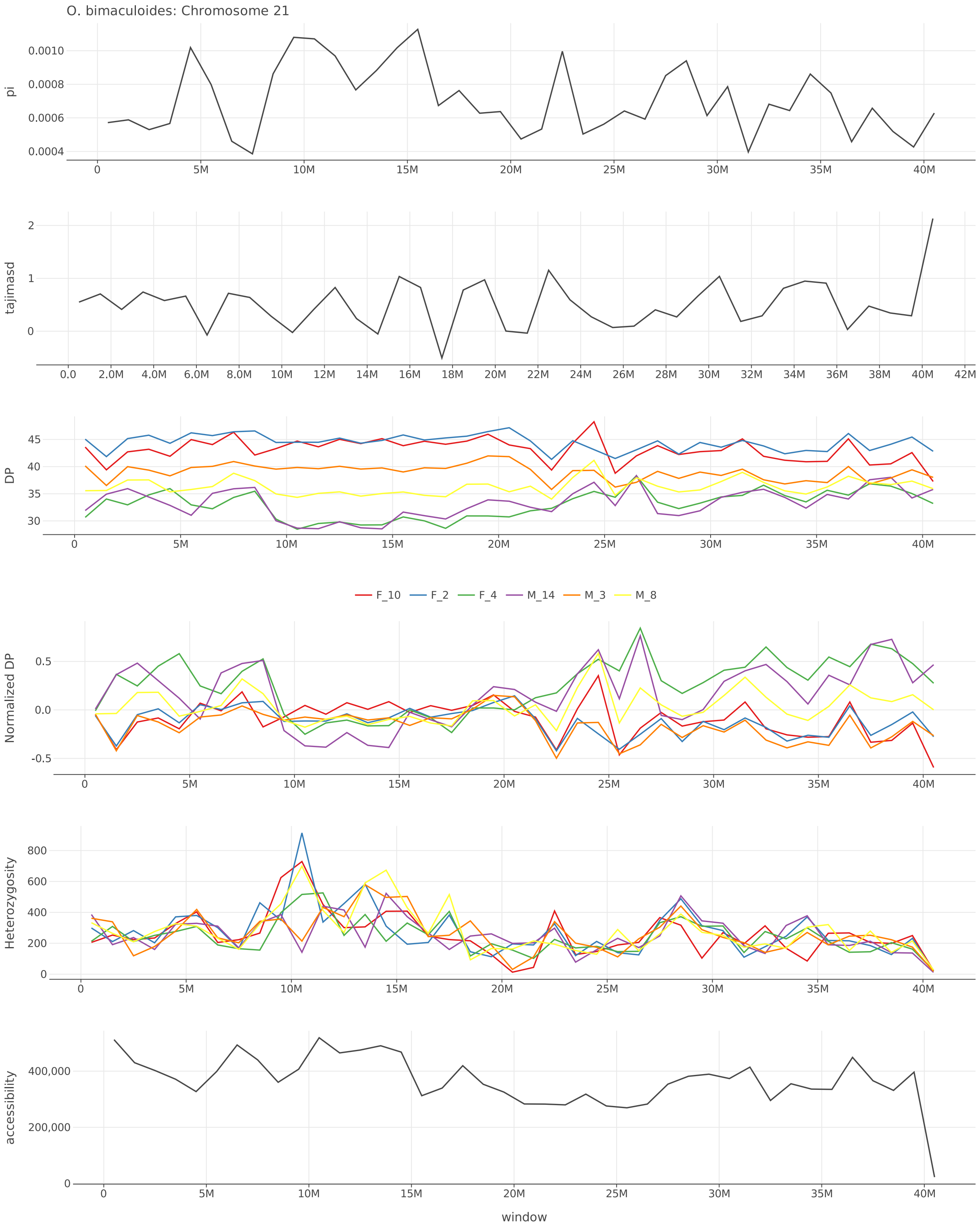


Figure S25: $\pi$, Tajima's $D$, depth, normalized depth, heterozygosity, and accessibility across chromosome 21 of *O. bimaculoides*.

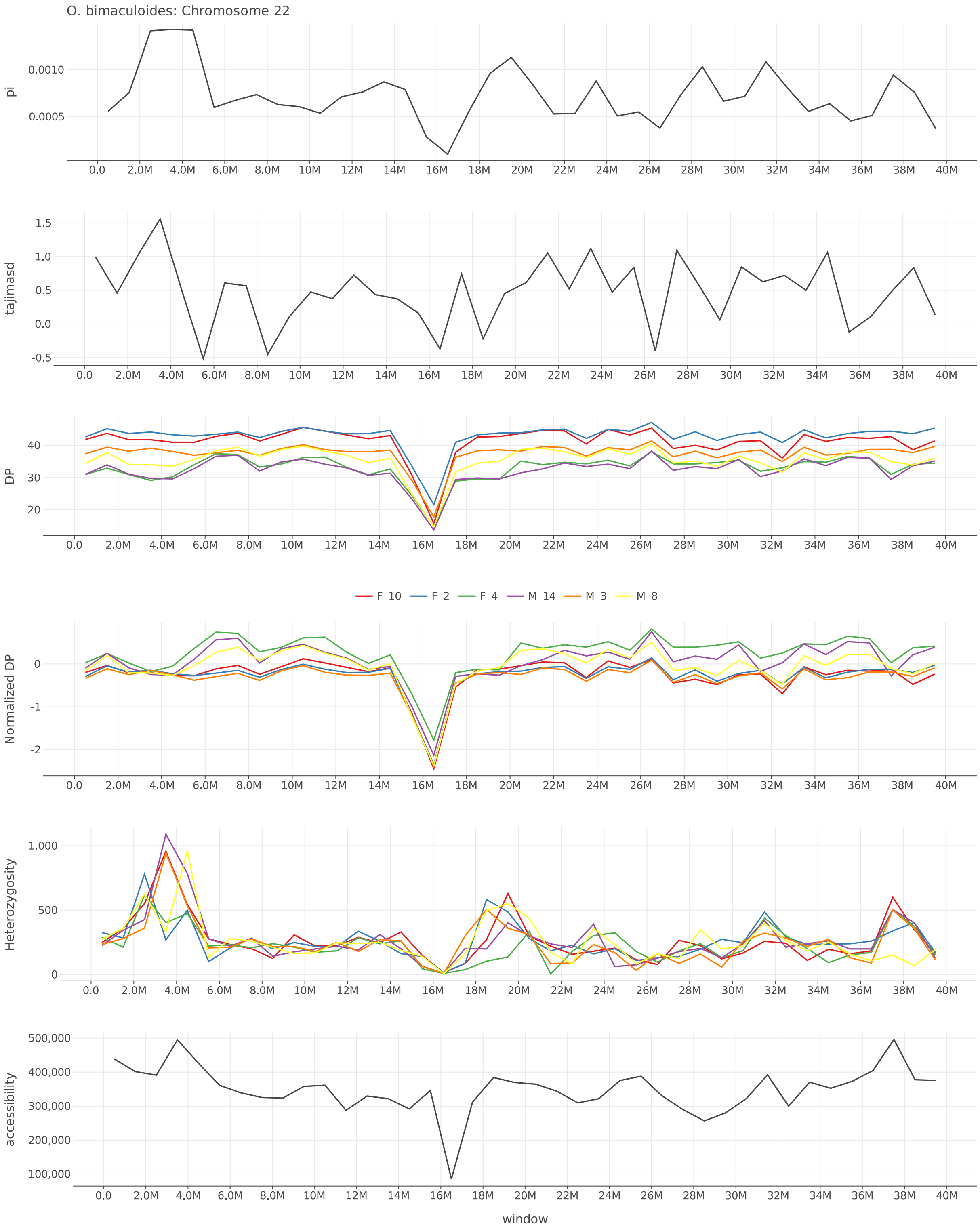


Figure S26: $\pi$, Tajima's $D$, depth, normalized depth, heterozygosity, and accessibility across chromosome 22 of *O. bimaculoides*.

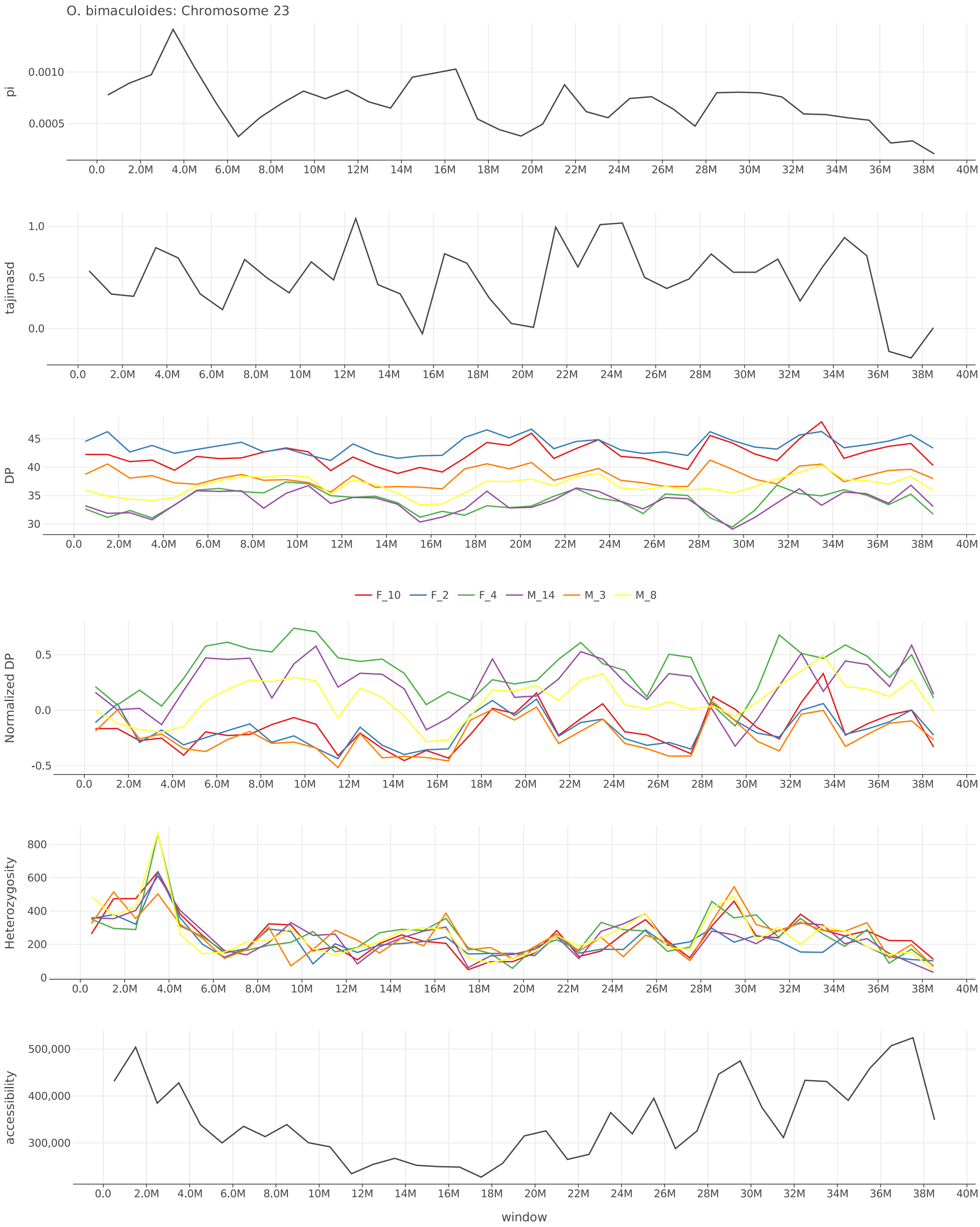


Figure S27: $\pi$, Tajima's $D$, depth, normalized depth, heterozygosity, and accessibility across chromosome 23 of *O. bimaculoides*.

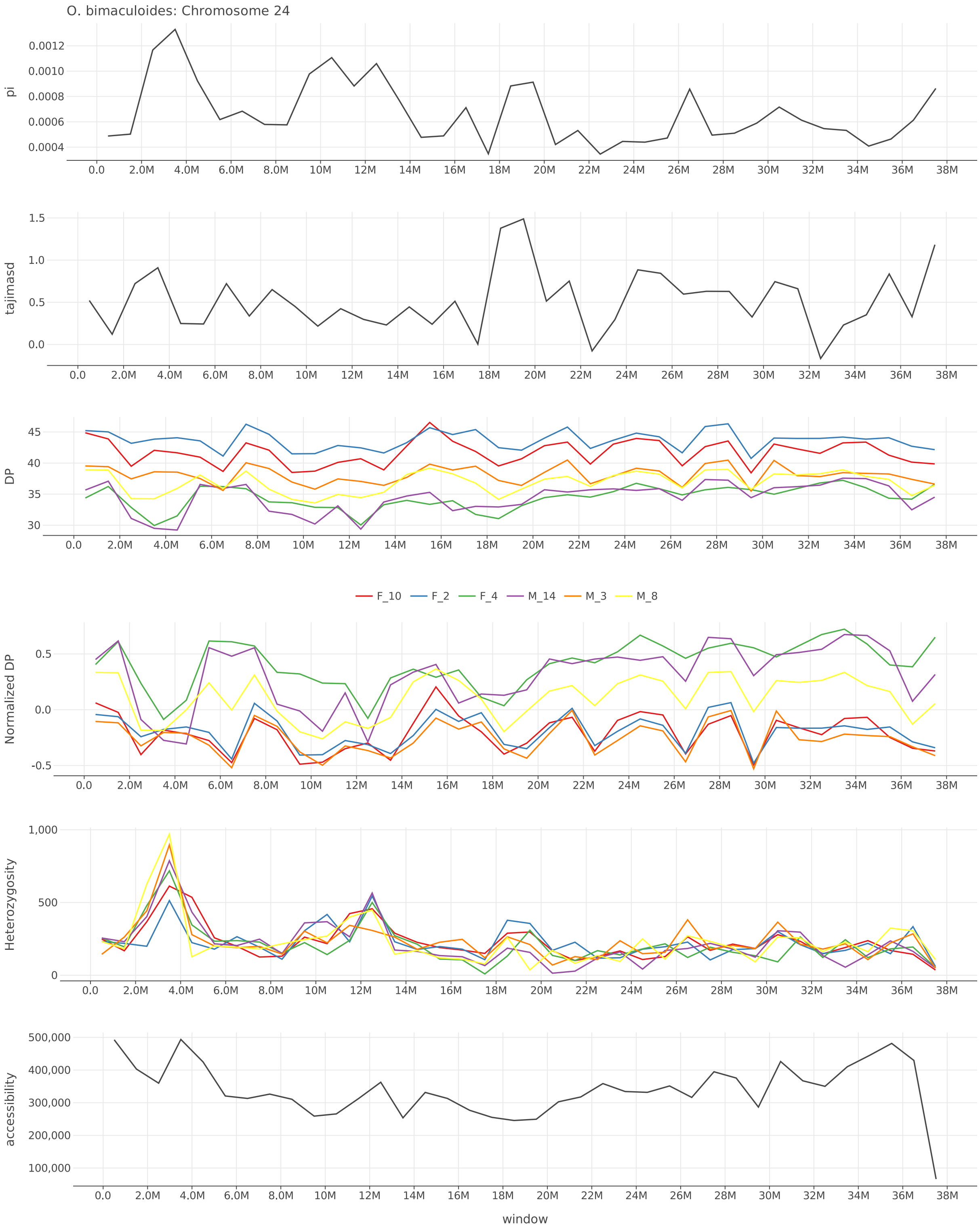


Figure S28: $\pi$, Tajima's $D$, depth, normalized depth, heterozygosity, and accessibility across chromosome 24 of *O. bimaculoides*.

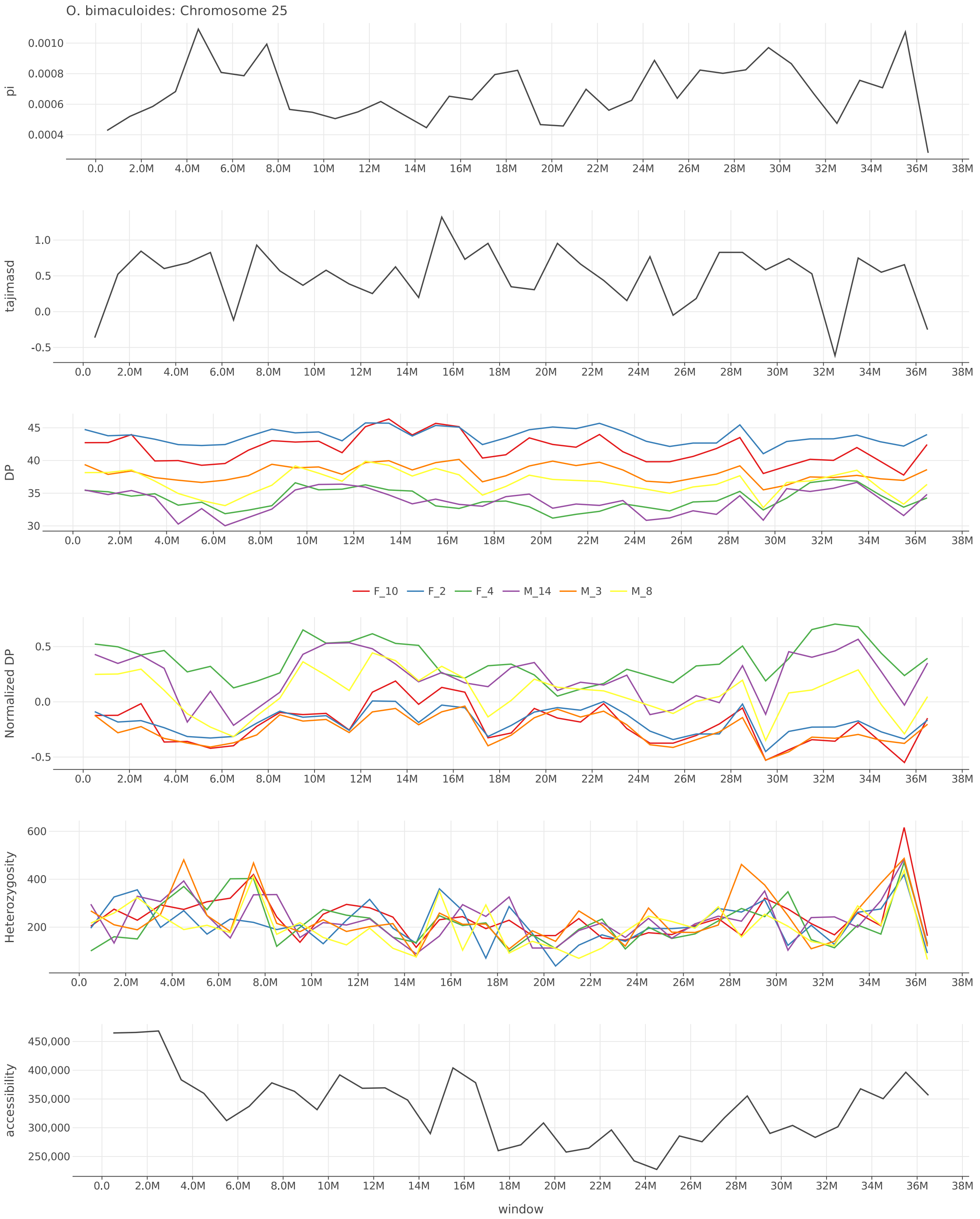


Figure S29: $\pi$, Tajima's $D$, depth, normalized depth, heterozygosity, and accessibility across chromosome 25 of *O. bimaculoides*.

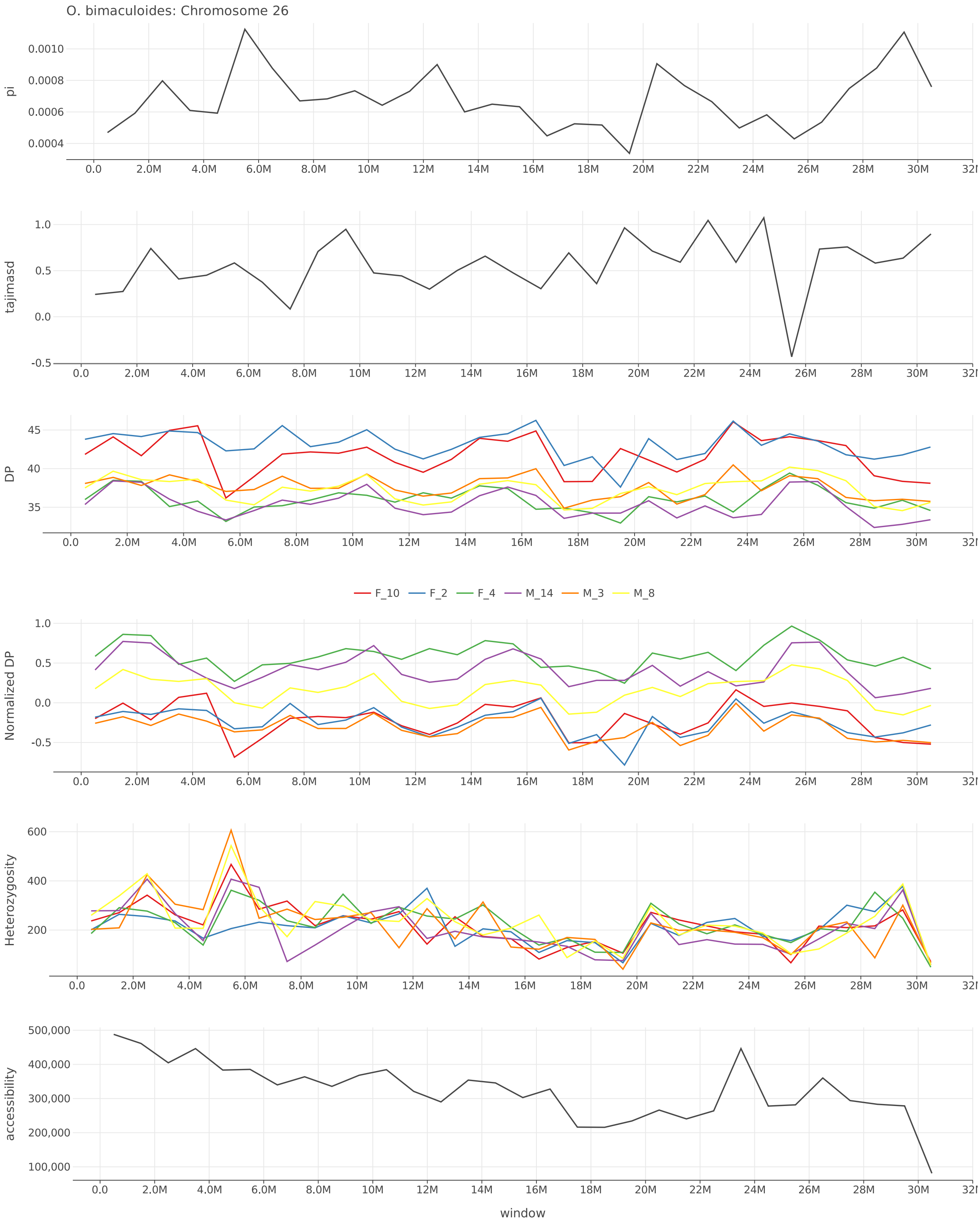


Figure S30: $\pi$, Tajima's $D$, depth, normalized depth, heterozygosity, and accessibility across chromosome 26 of *O. bimaculoides*.

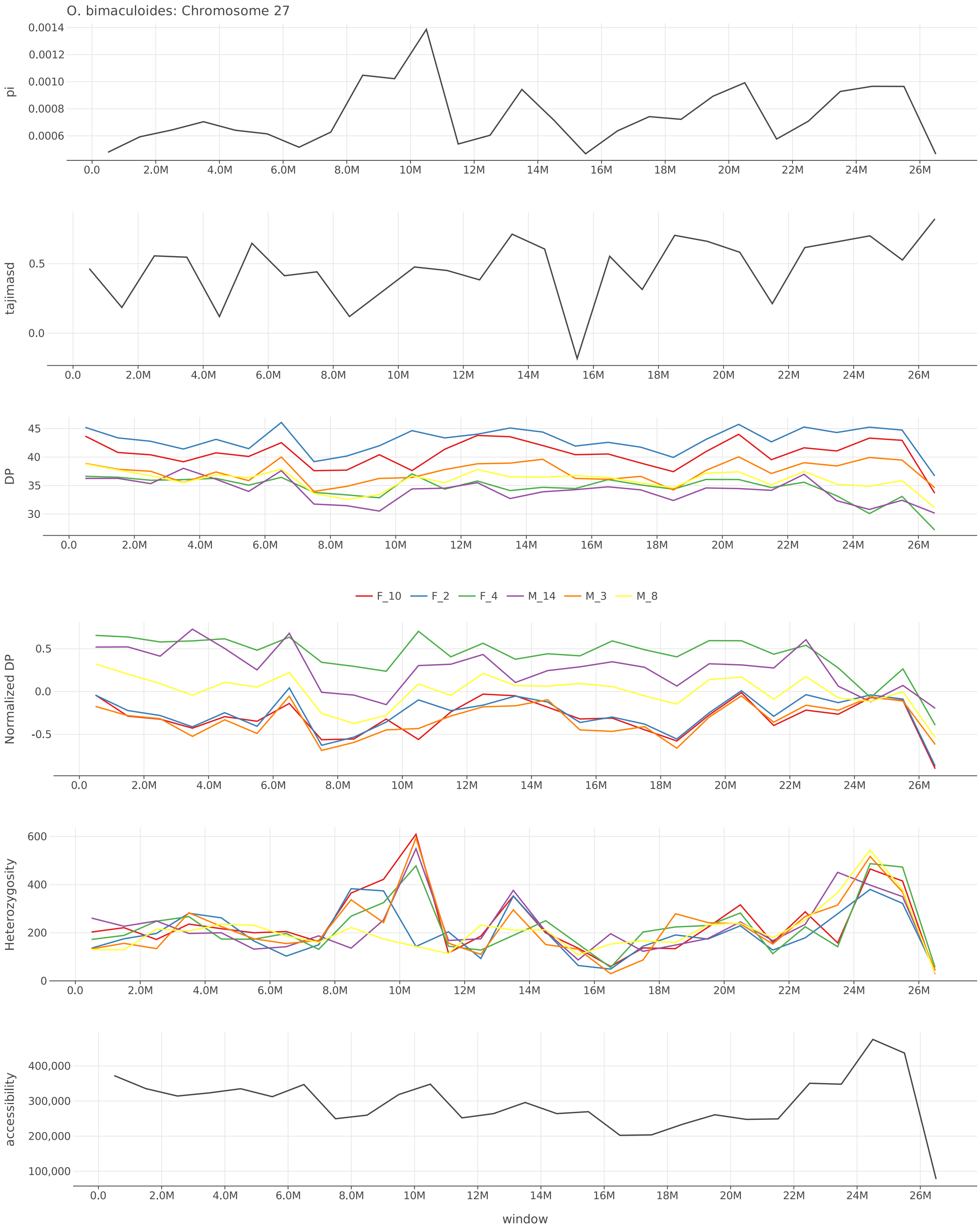


Figure S31: $\pi$, Tajima's $D$, depth, normalized depth, heterozygosity, and accessibility across chromosome 27 of *O. bimaculoides*.

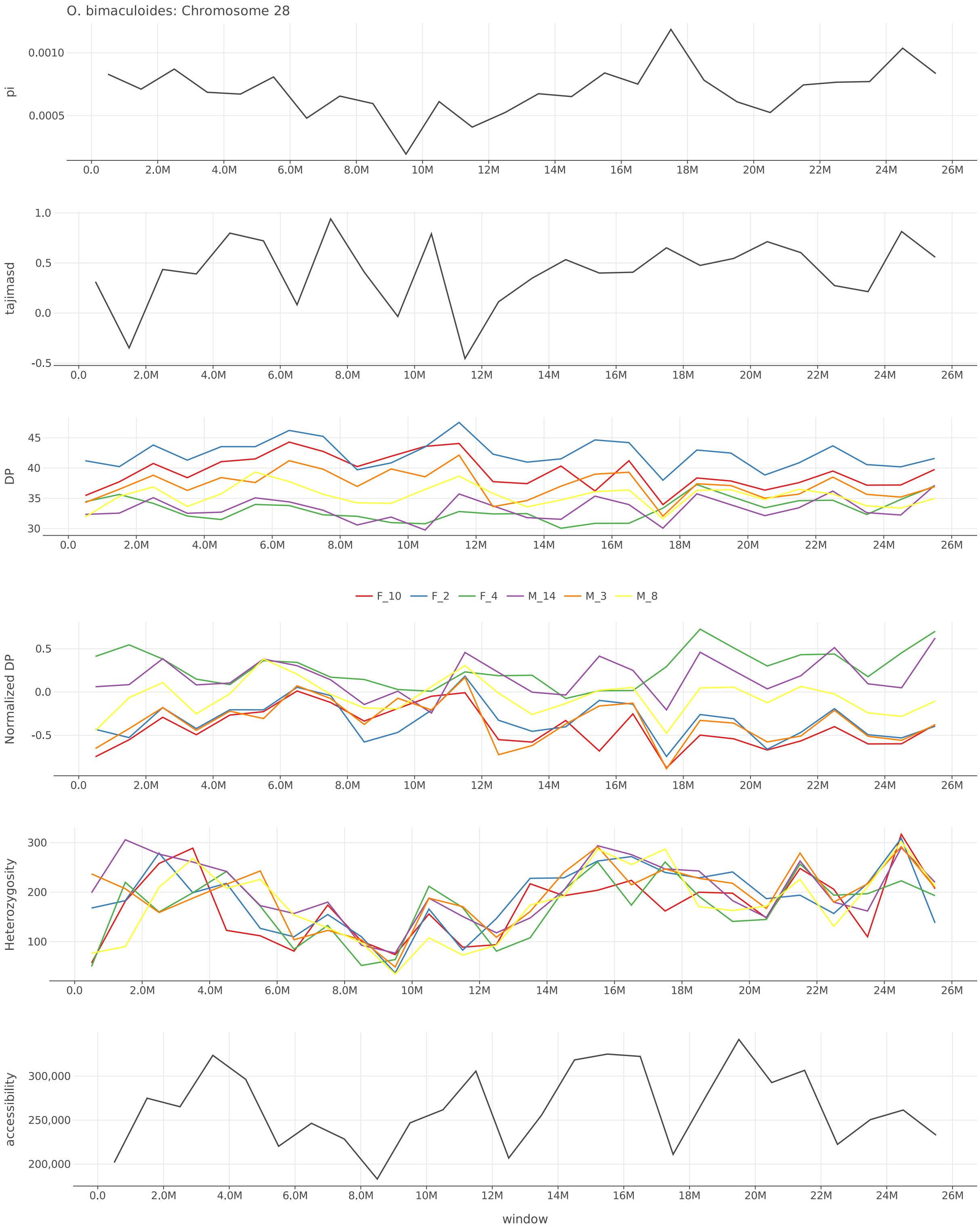


Figure S32: $\pi$, Tajima's $D$, depth, normalized depth, heterozygosity, and accessibility across chromosome 28 of *O. bimaculoides*.

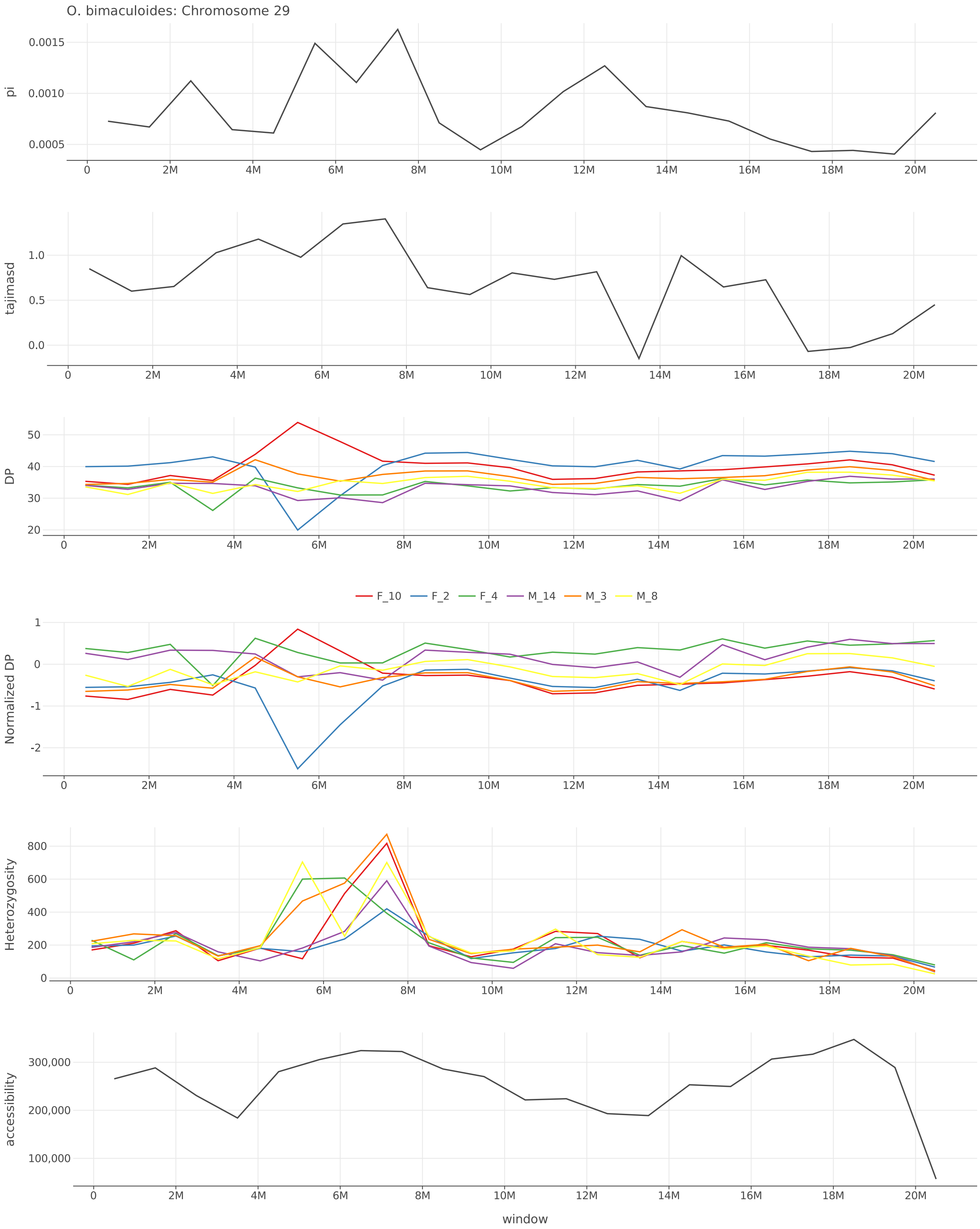


Figure S33: $\pi$, Tajima's $D$, depth, normalized depth, heterozygosity, and accessibility across chromosome 29 of *O. bimaculoides*.

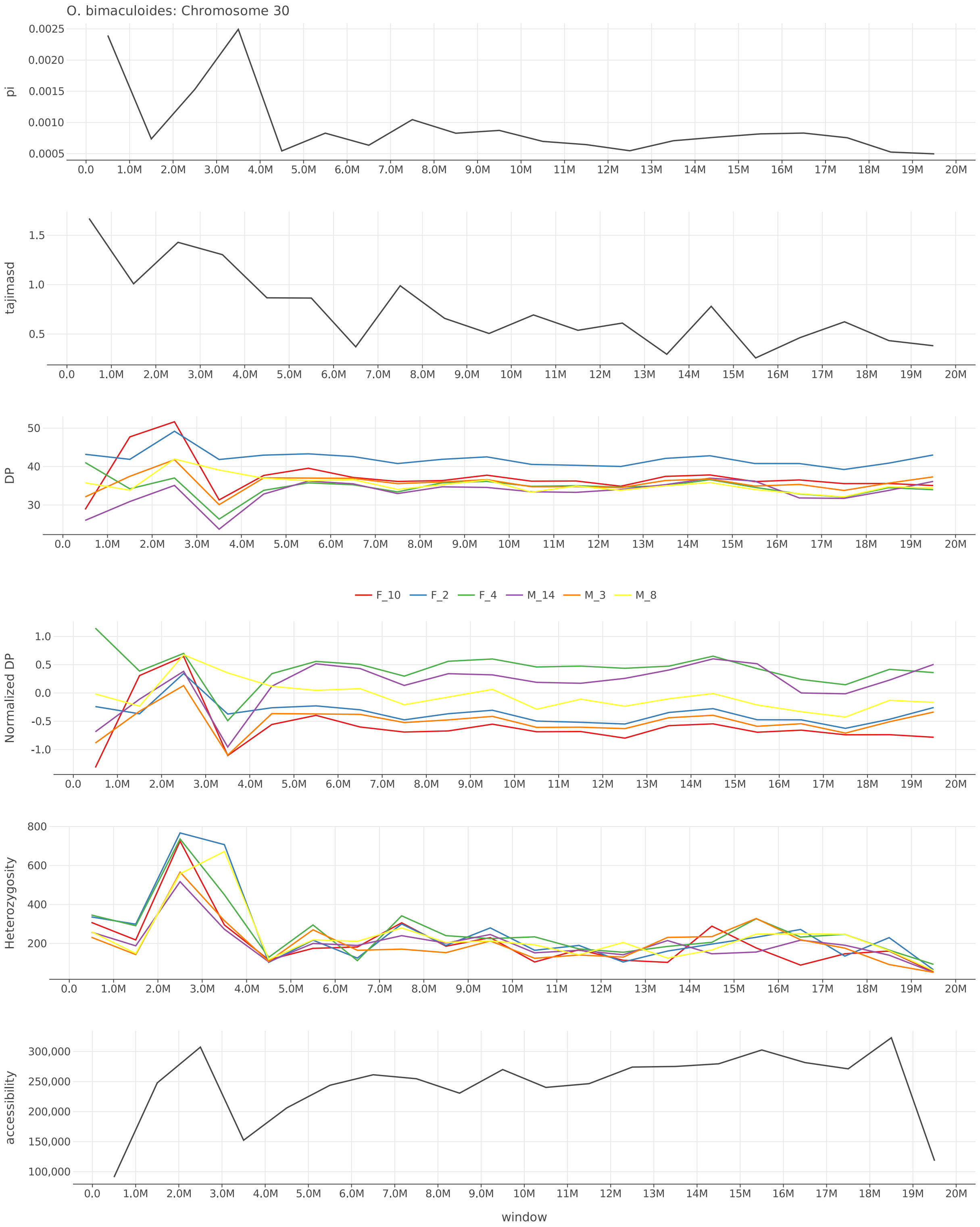


Figure S34: $\pi$, Tajima's $D$, depth, normalized depth, heterozygosity, and accessibility across chromosome 30 of *O. bimaculoides*.

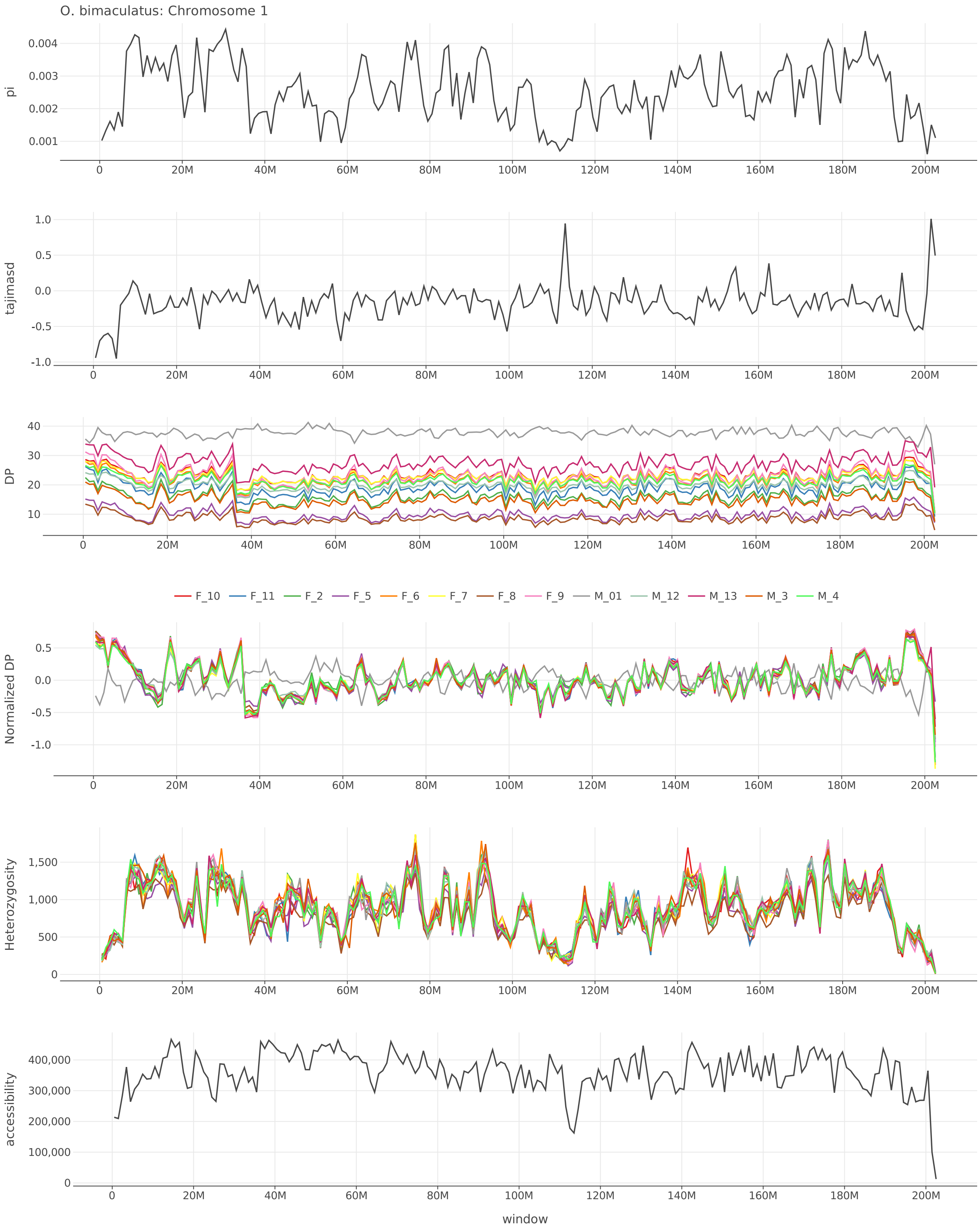


Figure S35: $\pi$, Tajima's $D$, depth, normalized depth, heterozygosity, and accessibility across chromosome 1 of *O. bimaculatus*.

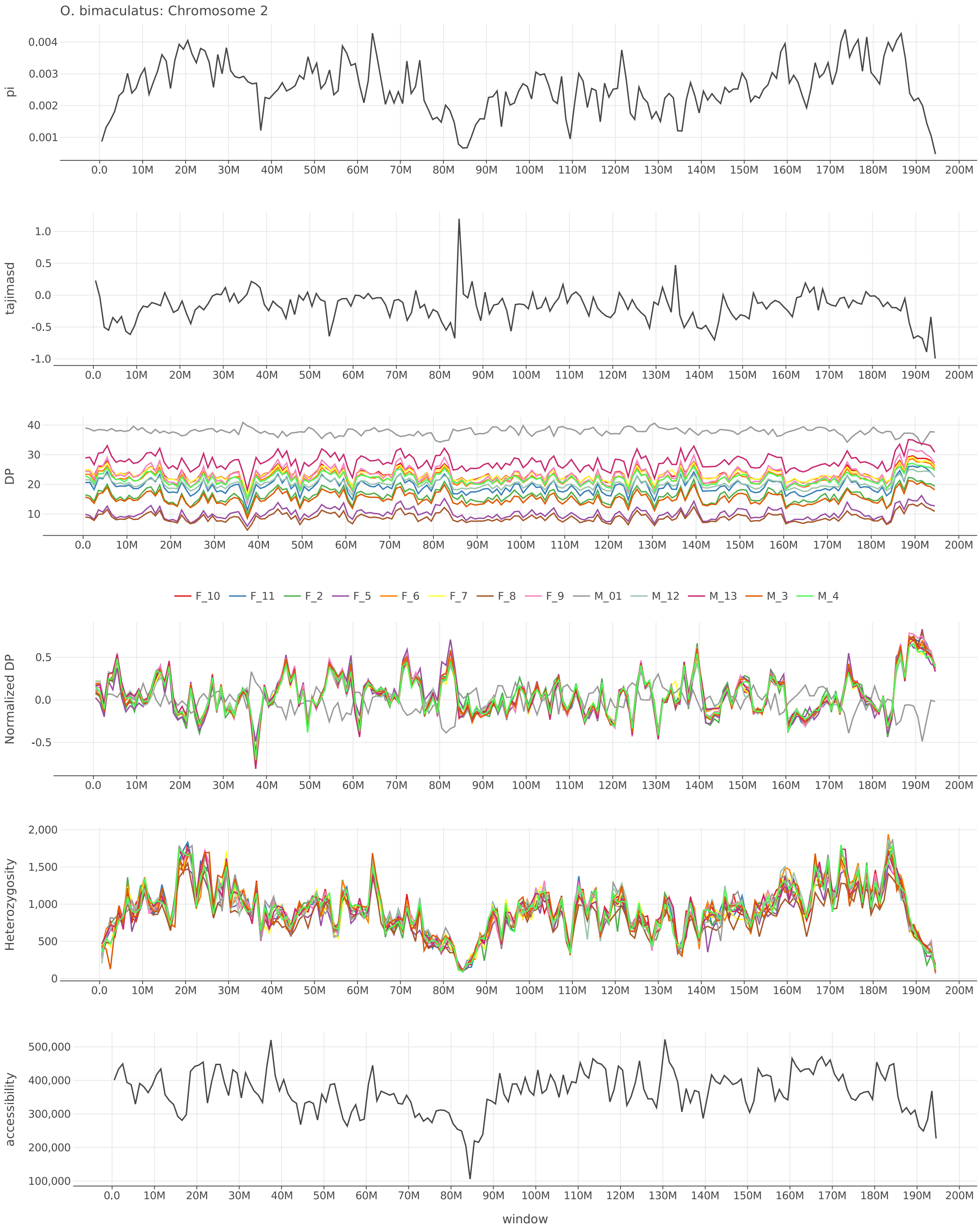


Figure S36: $\pi$, Tajima's $D$, depth, normalized depth, heterozygosity, and accessibility across chromosome 2 of *O. bimaculatus*.

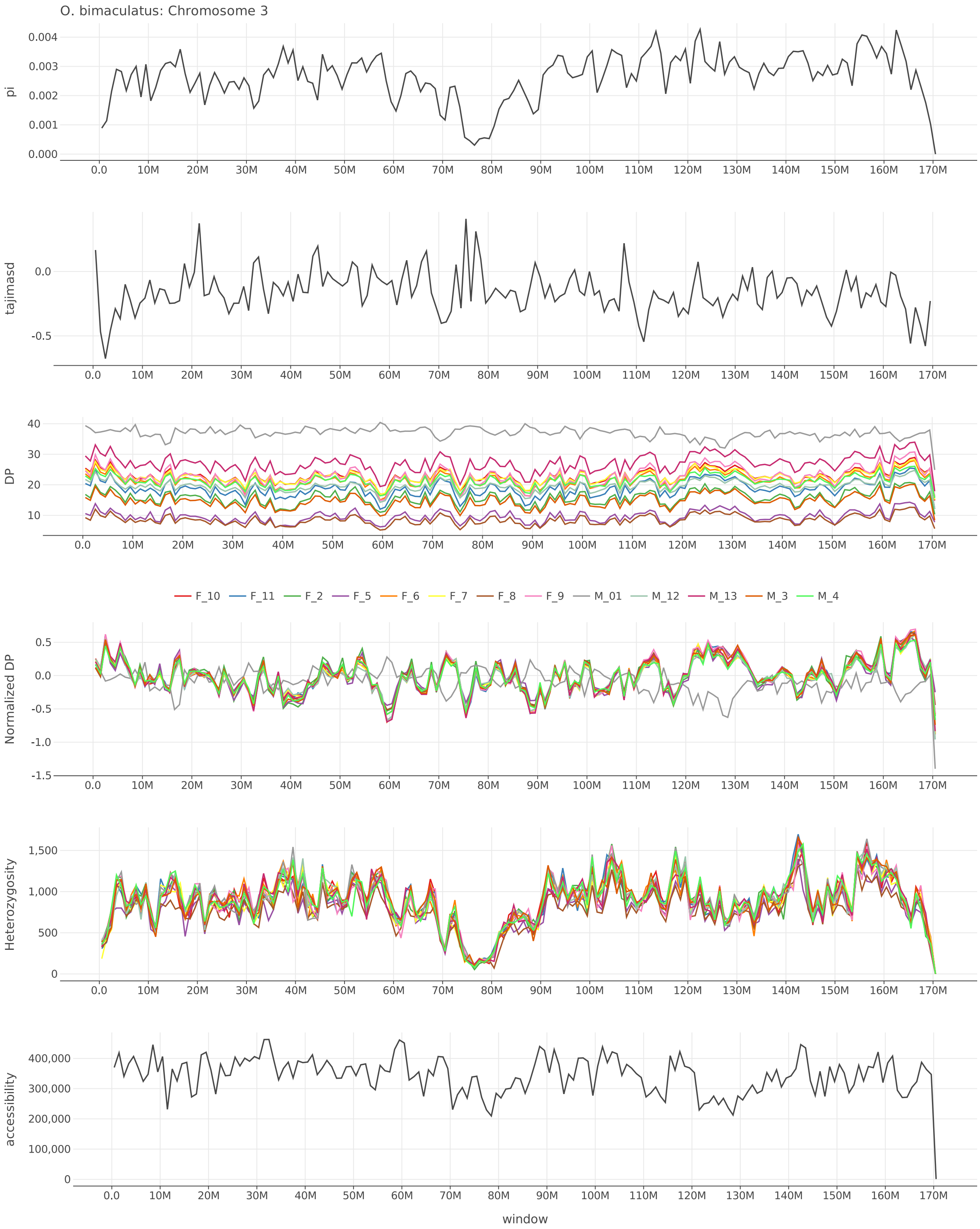


Figure S37: $\pi$, Tajima's $D$, depth, normalized depth, heterozygosity, and accessibility across chromosome 3 of *O. bimaculatus*.

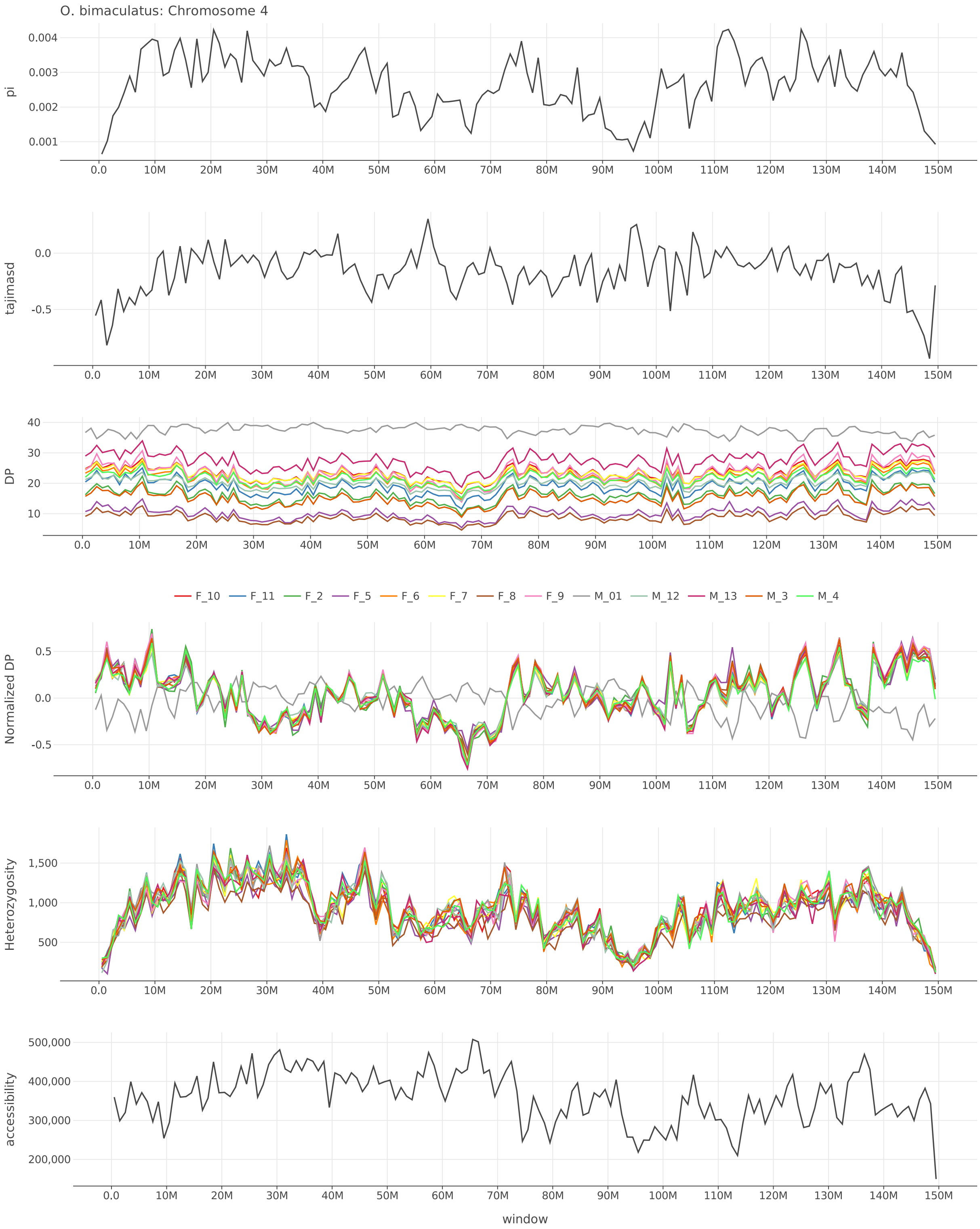


Figure S38: $\pi$, Tajima's $D$, depth, normalized depth, heterozygosity, and accessibility across chromosome 4 of *O. bimaculatus*.

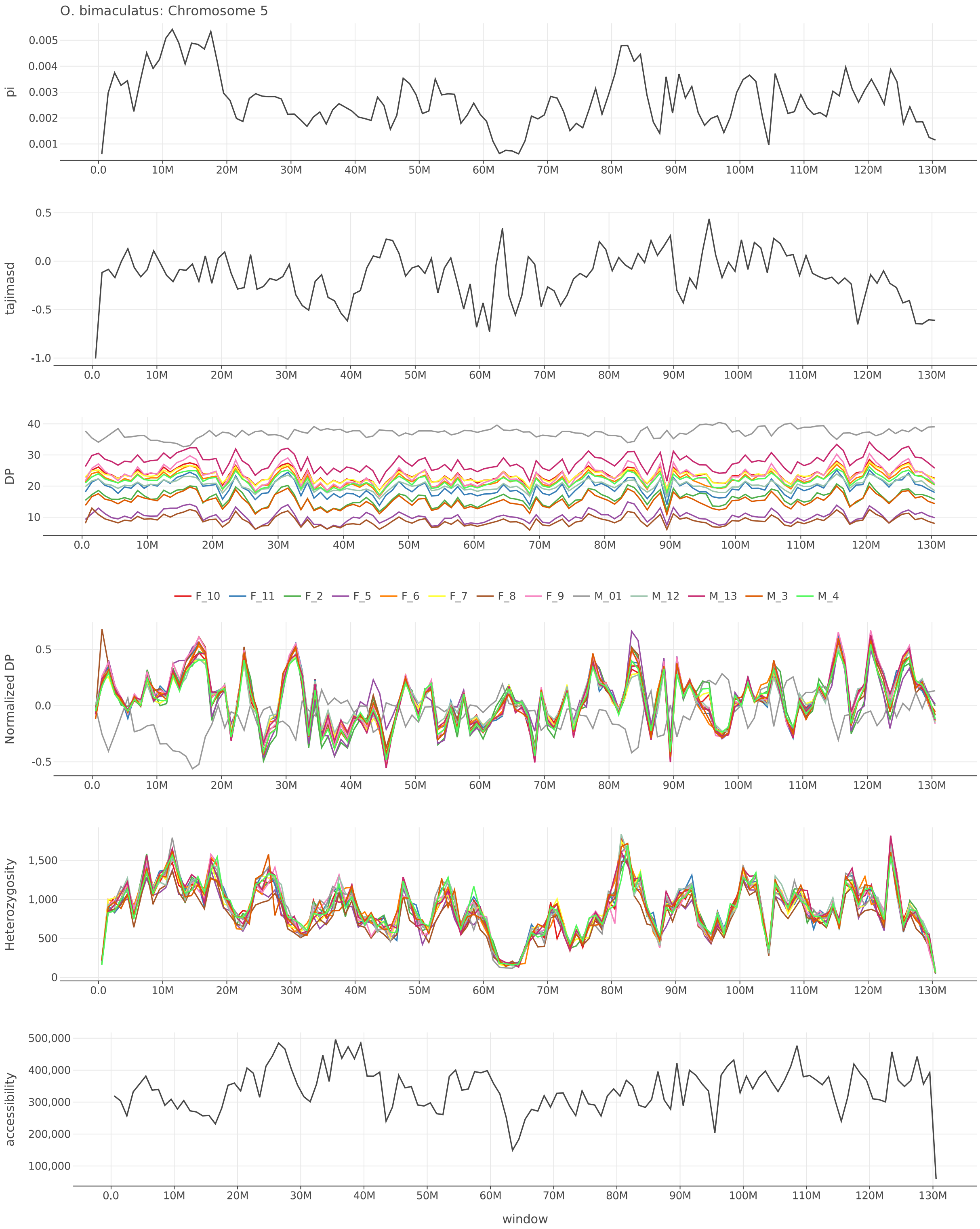


Figure S39: $\pi$, Tajima's $D$, depth, normalized depth, heterozygosity, and accessibility across chromosome 5 of *O. bimaculatus*.

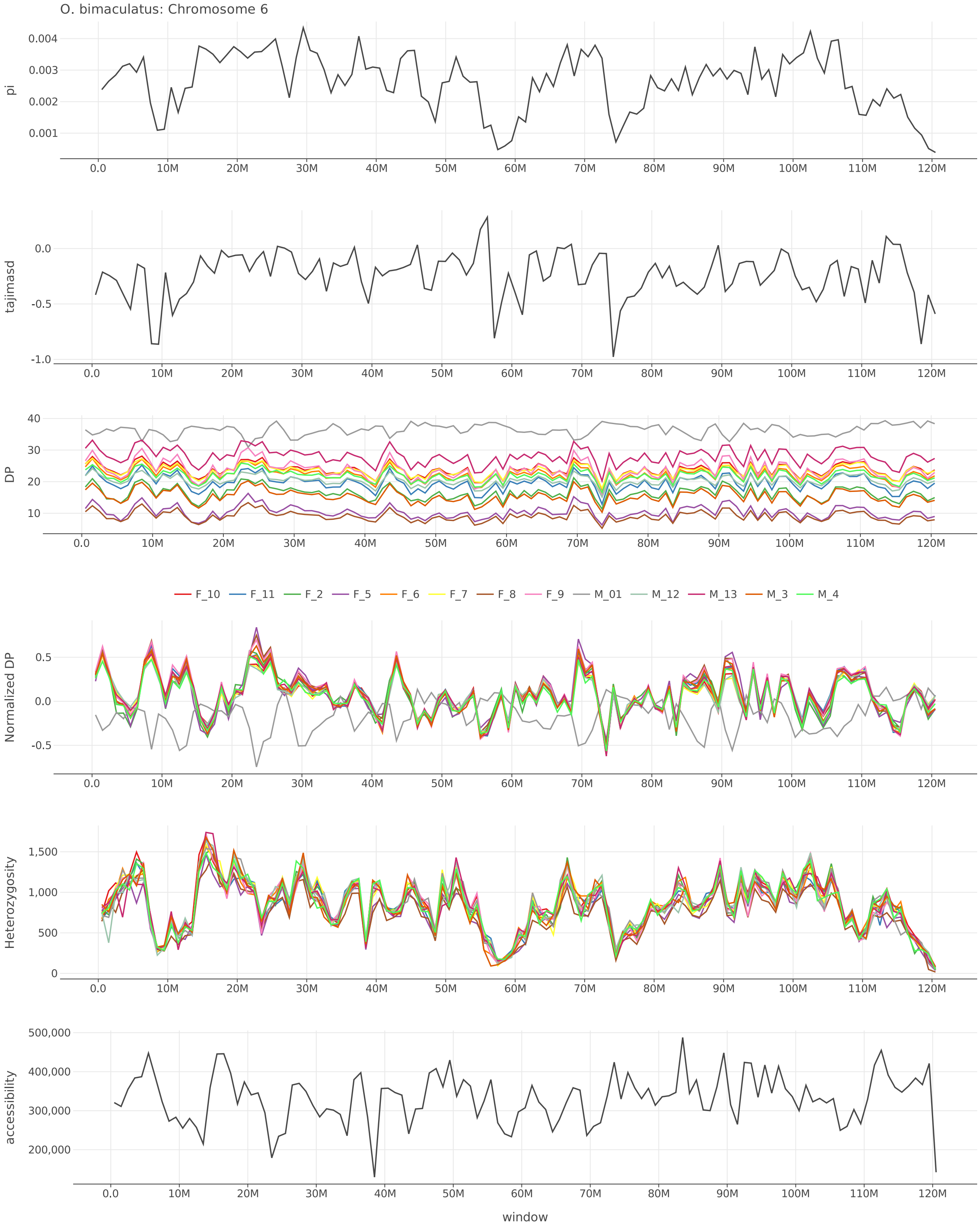


Figure S40: $\pi$, Tajima's $D$, depth, normalized depth, heterozygosity, and accessibility across chromosome 6 of *O. bimaculatus*.

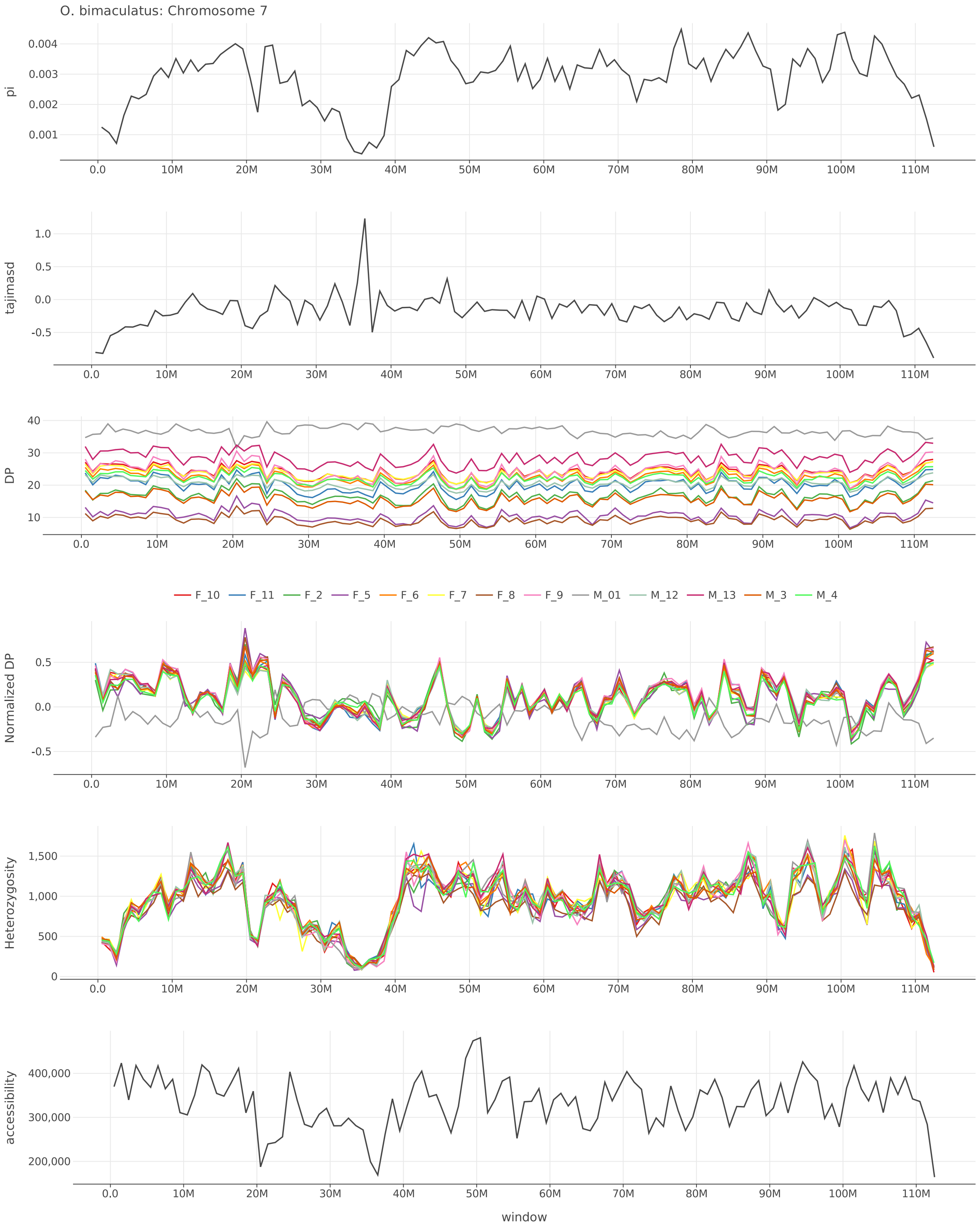


Figure S41: $\pi$, Tajima's $D$, depth, normalized depth, heterozygosity, and accessibility across chromosome 7 of *O. bimaculatus*.

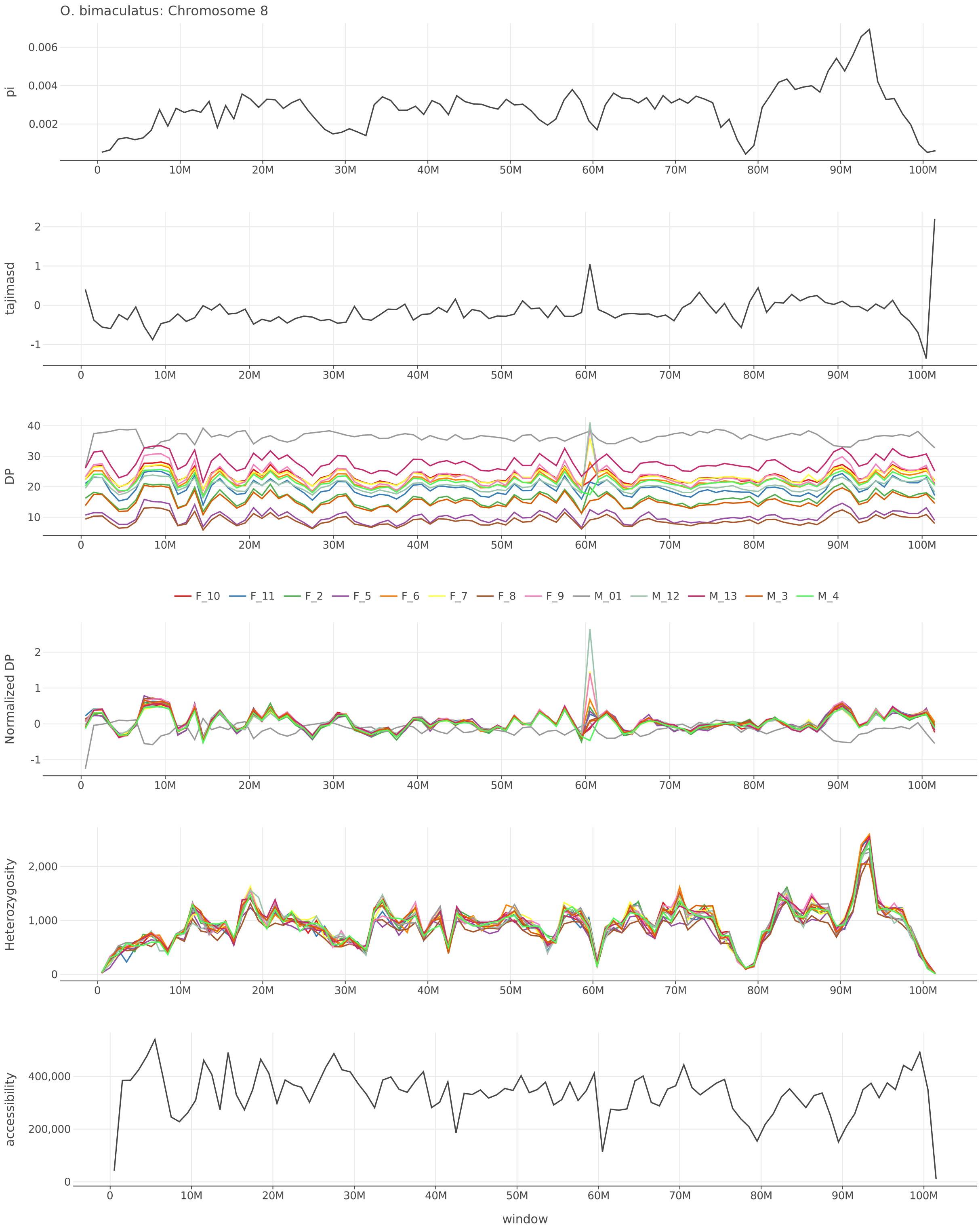


Figure S42: $\pi$, Tajima's $D$, depth, normalized depth, heterozygosity, and accessibility across chromosome 8 of *O. bimaculatus*.

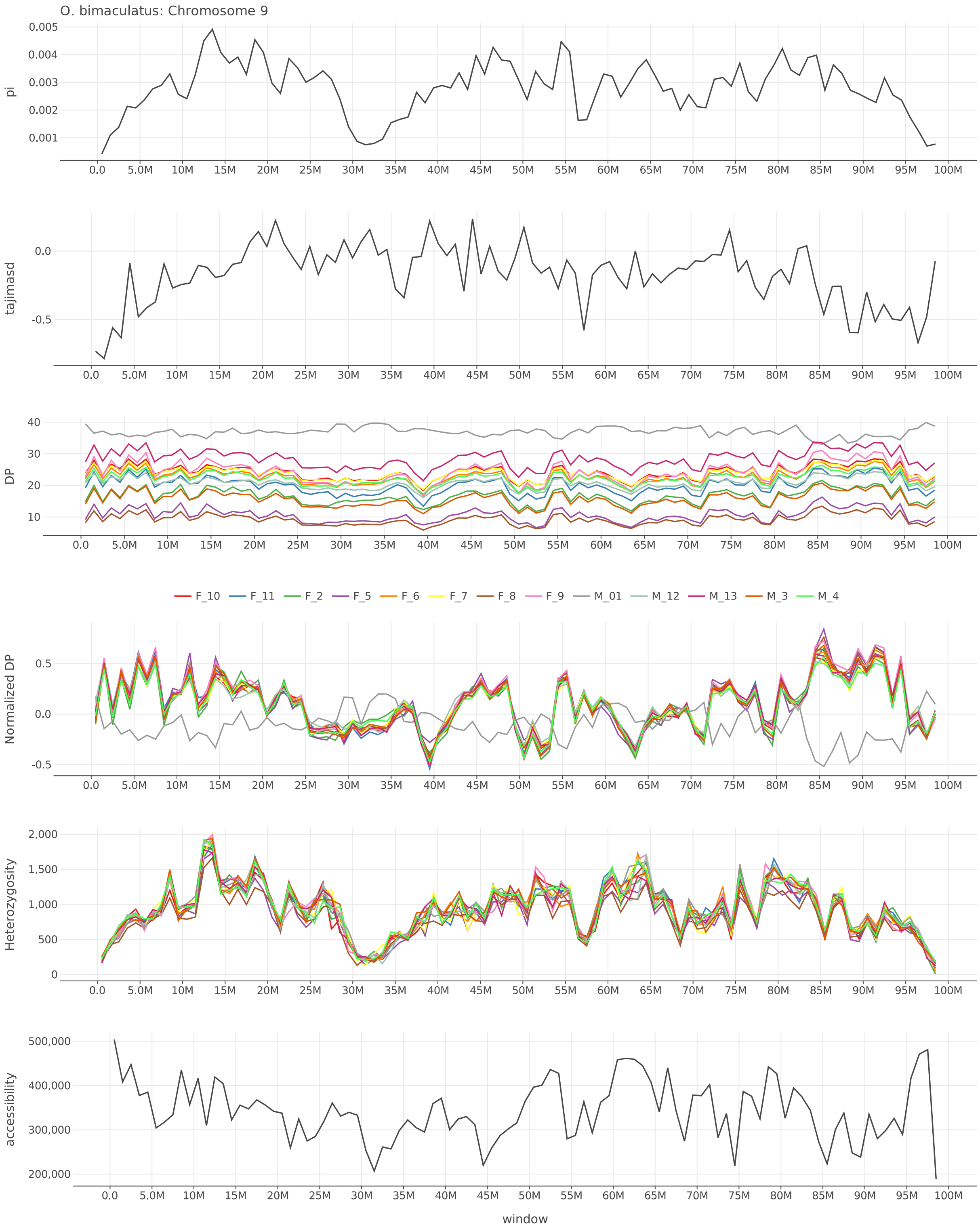


Figure S43: $\pi$, Tajima's $D$, depth, normalized depth, heterozygosity, and accessibility across chromosome 9 of *O. bimaculatus*.

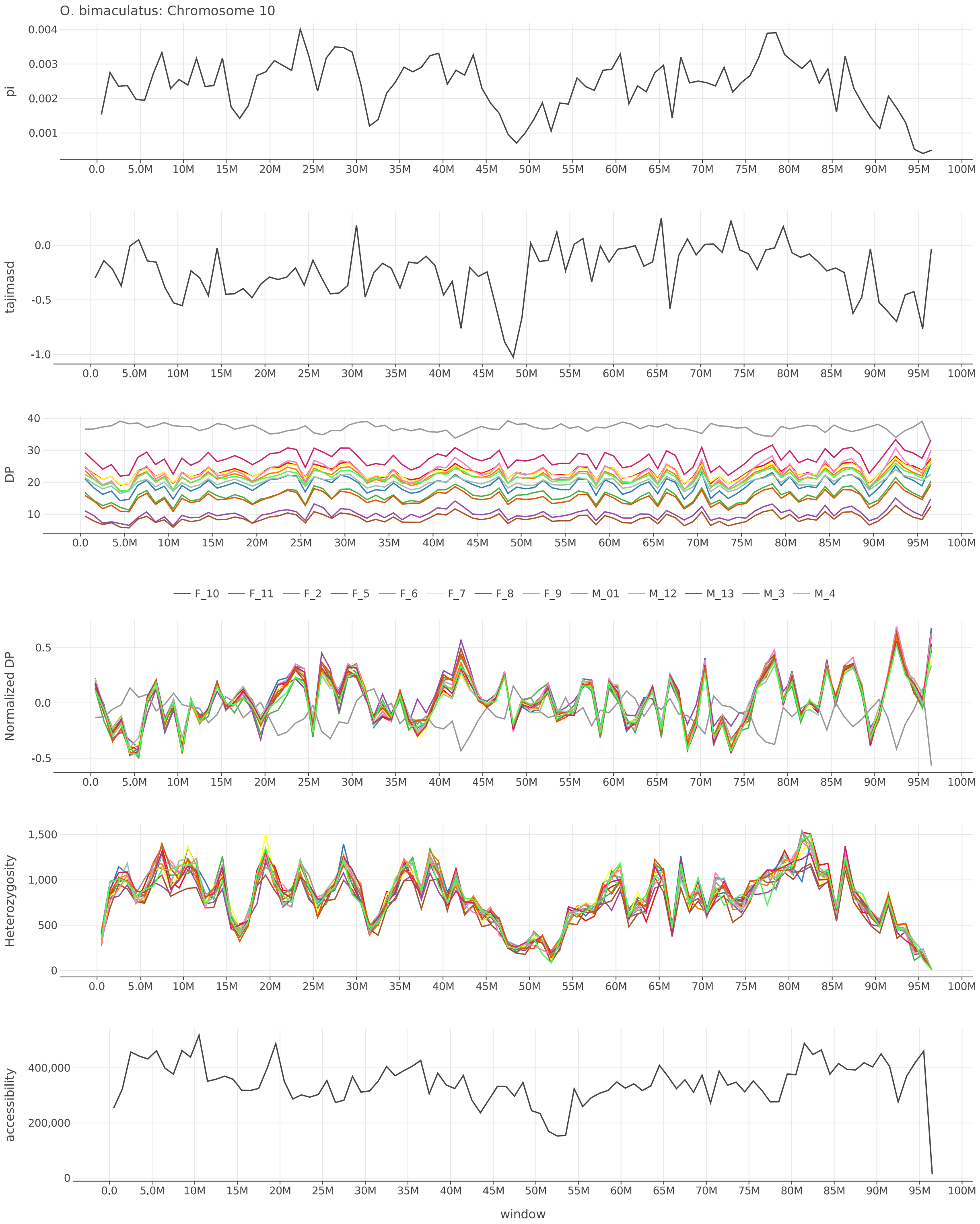


Figure S44: $\pi$, Tajima's $D$, depth, normalized depth, heterozygosity, and accessibility across chromosome 10 of *O. bimaculatus*.

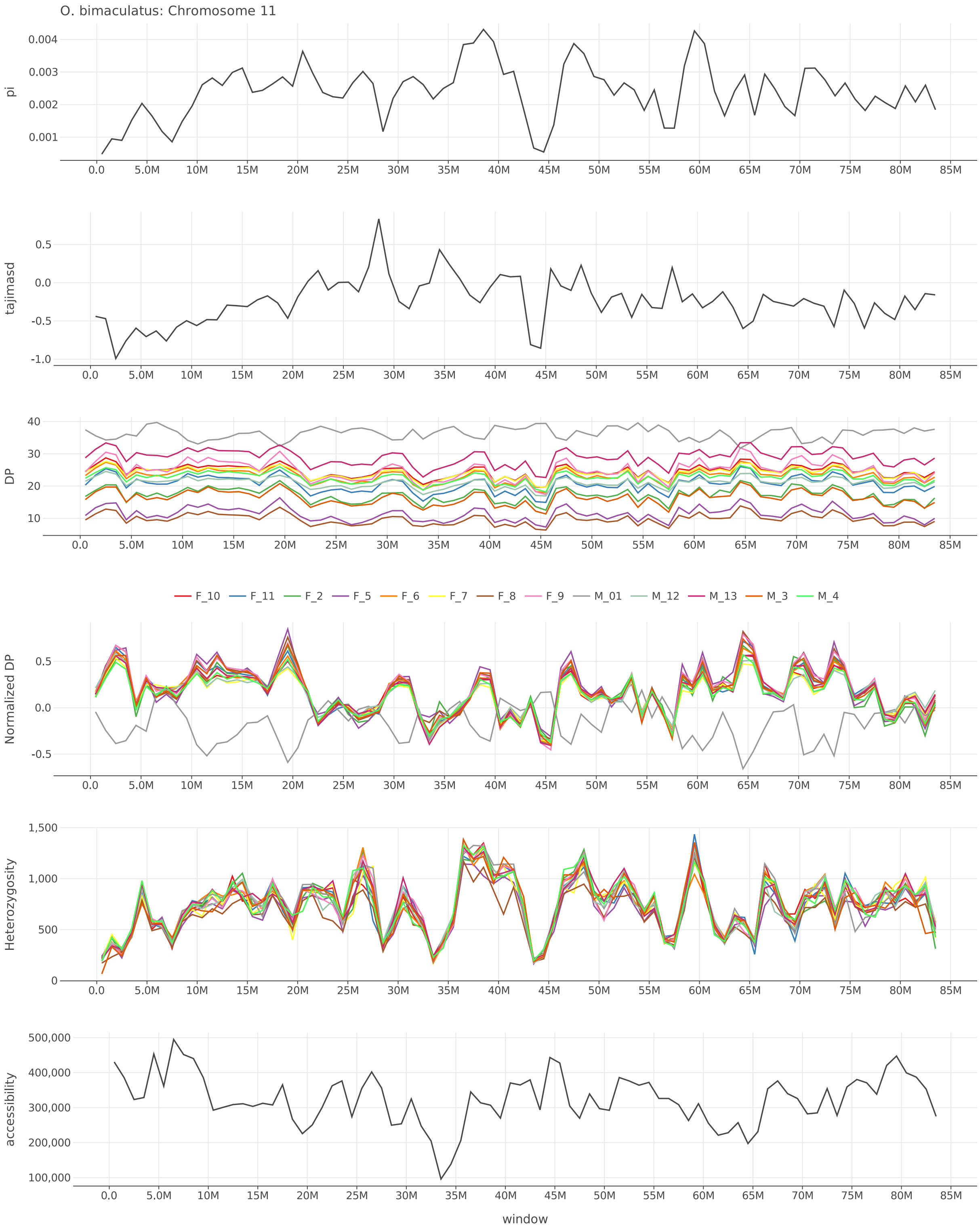


Figure S45: $\pi$, Tajima's $D$, depth, normalized depth, heterozygosity, and accessibility across chromosome 11 of *O. bimaculatus*.

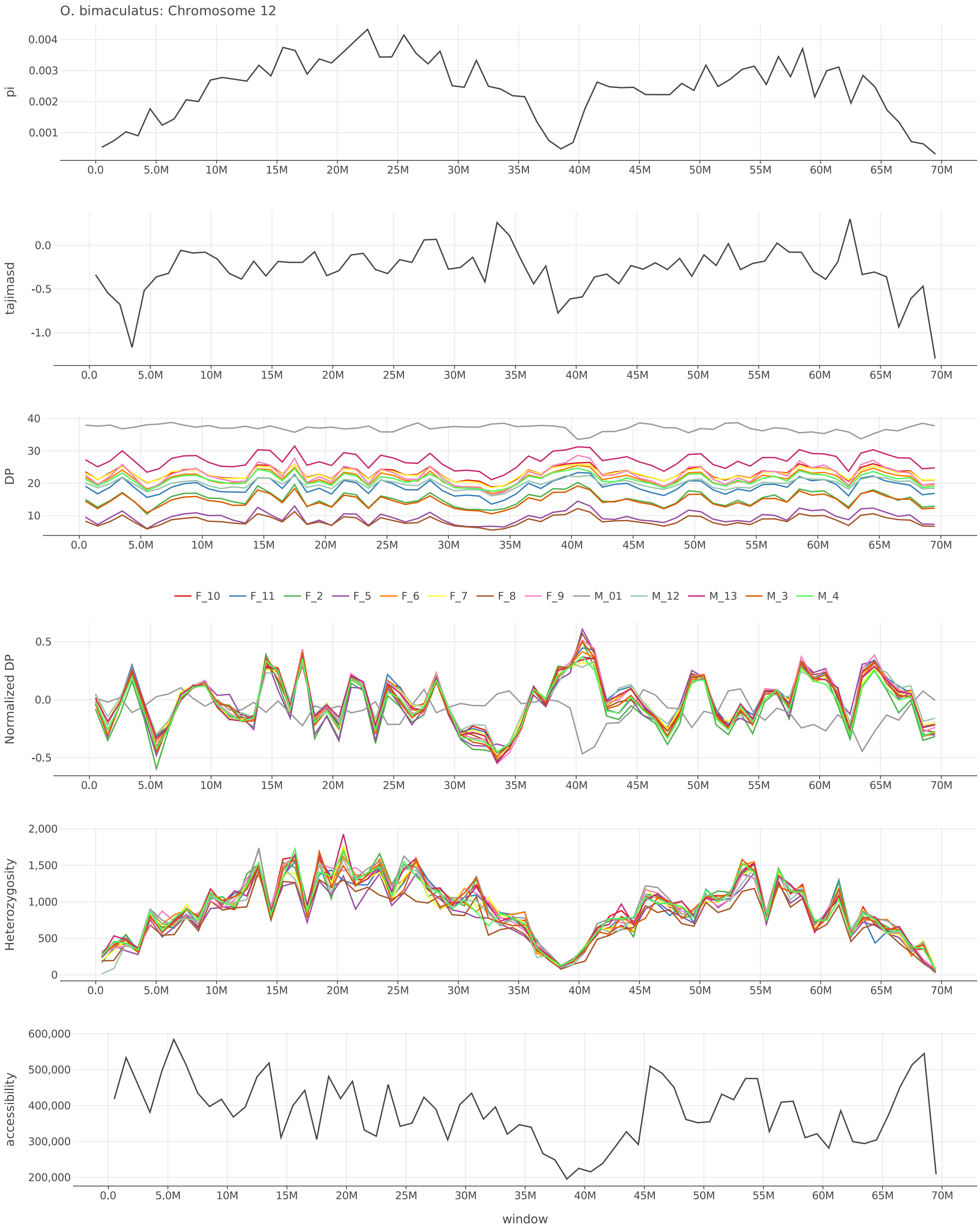


Figure S46: $\pi$, Tajima's $D$, depth, normalized depth, heterozygosity, and accessibility across chromosome 12 of *O. bimaculatus*.

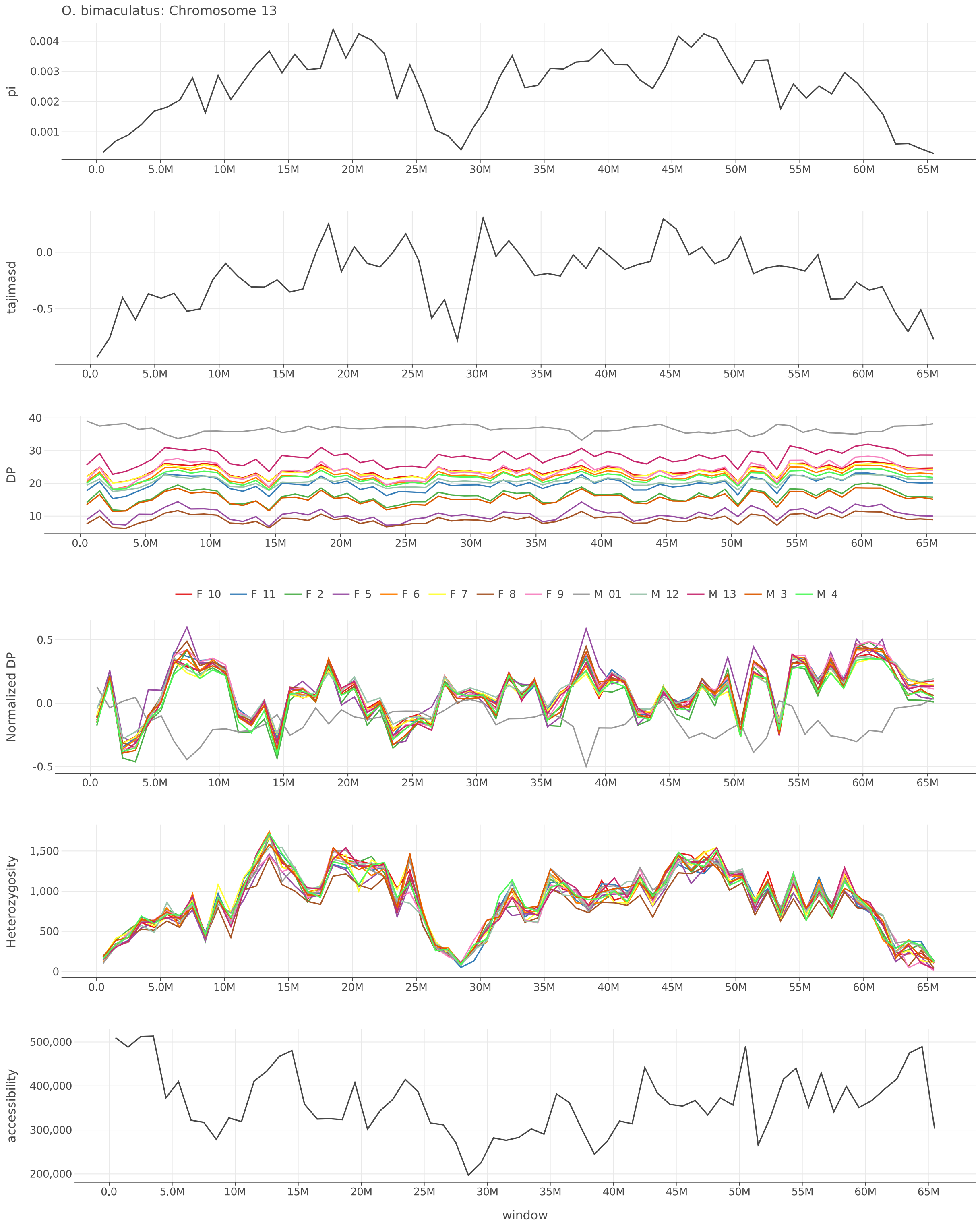


Figure S47: $\pi$, Tajima's $D$, depth, normalized depth, heterozygosity, and accessibility across chromosome 13 of *O. bimaculatus*.

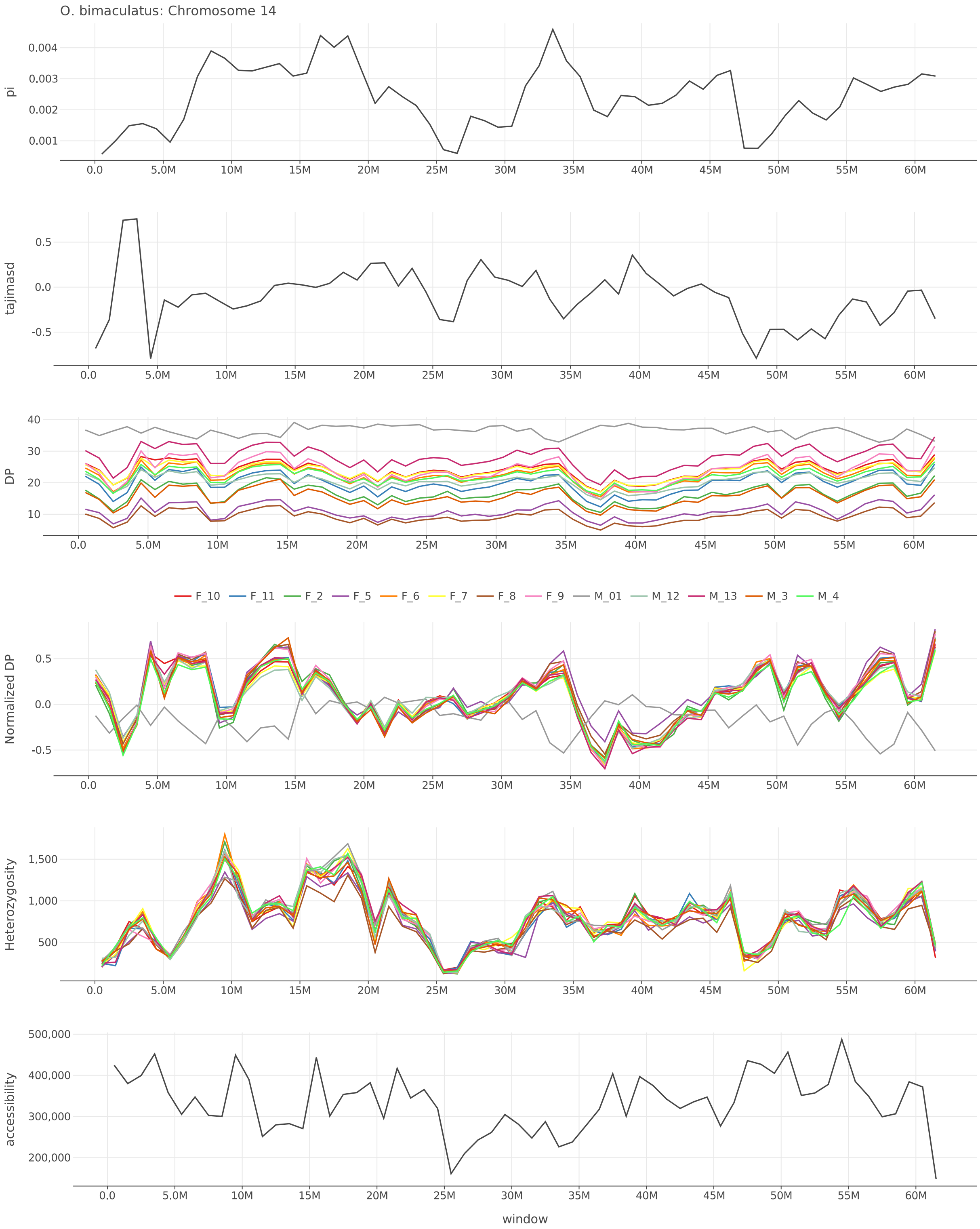


Figure S48: $\pi$, Tajima's $D$, depth, normalized depth, heterozygosity, and accessibility across chromosome 14 of *O. bimaculatus*.

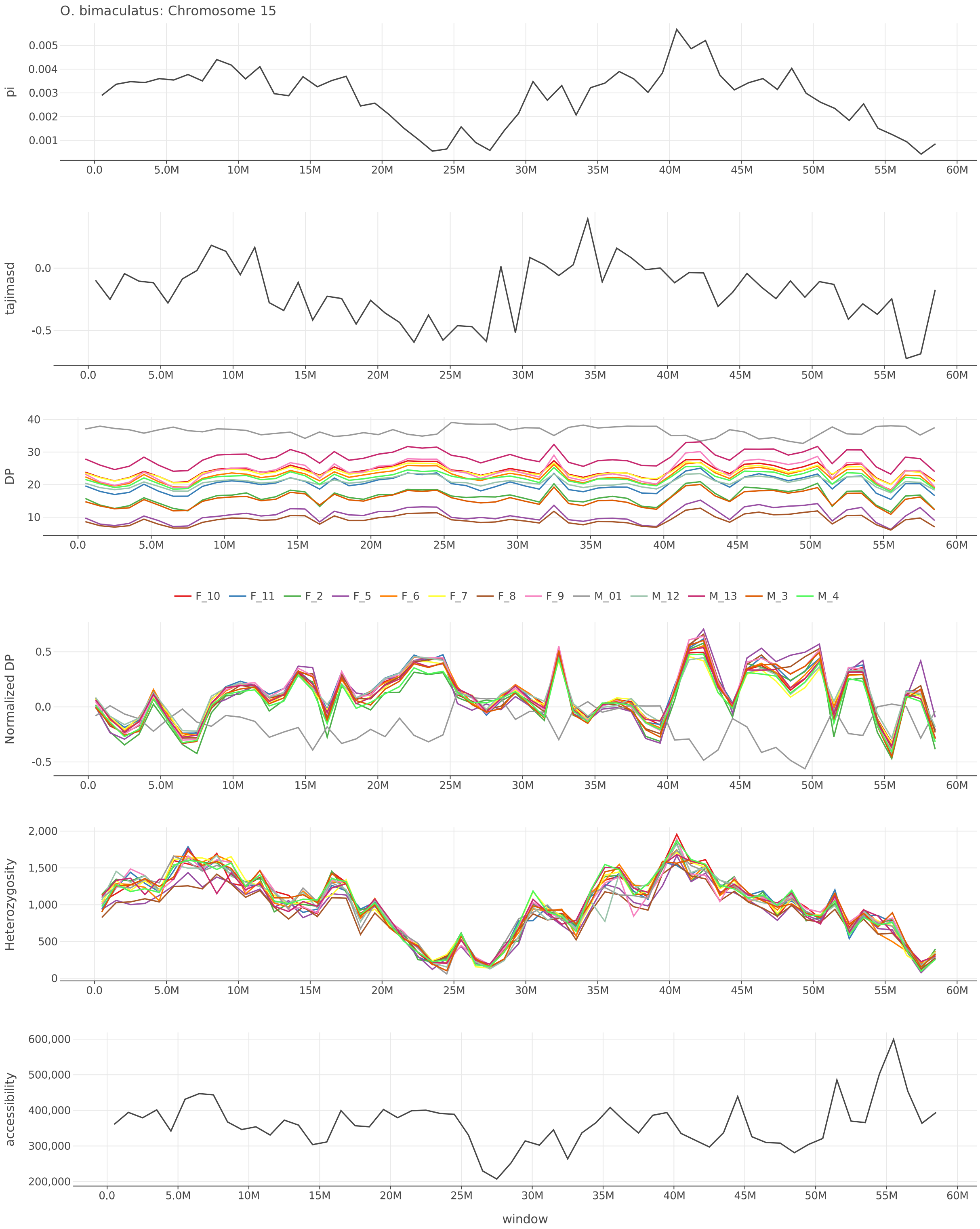


Figure S49: $\pi$, Tajima's $D$, depth, normalized depth, heterozygosity, and accessibility across chromosome 15 of *O. bimaculatus*.

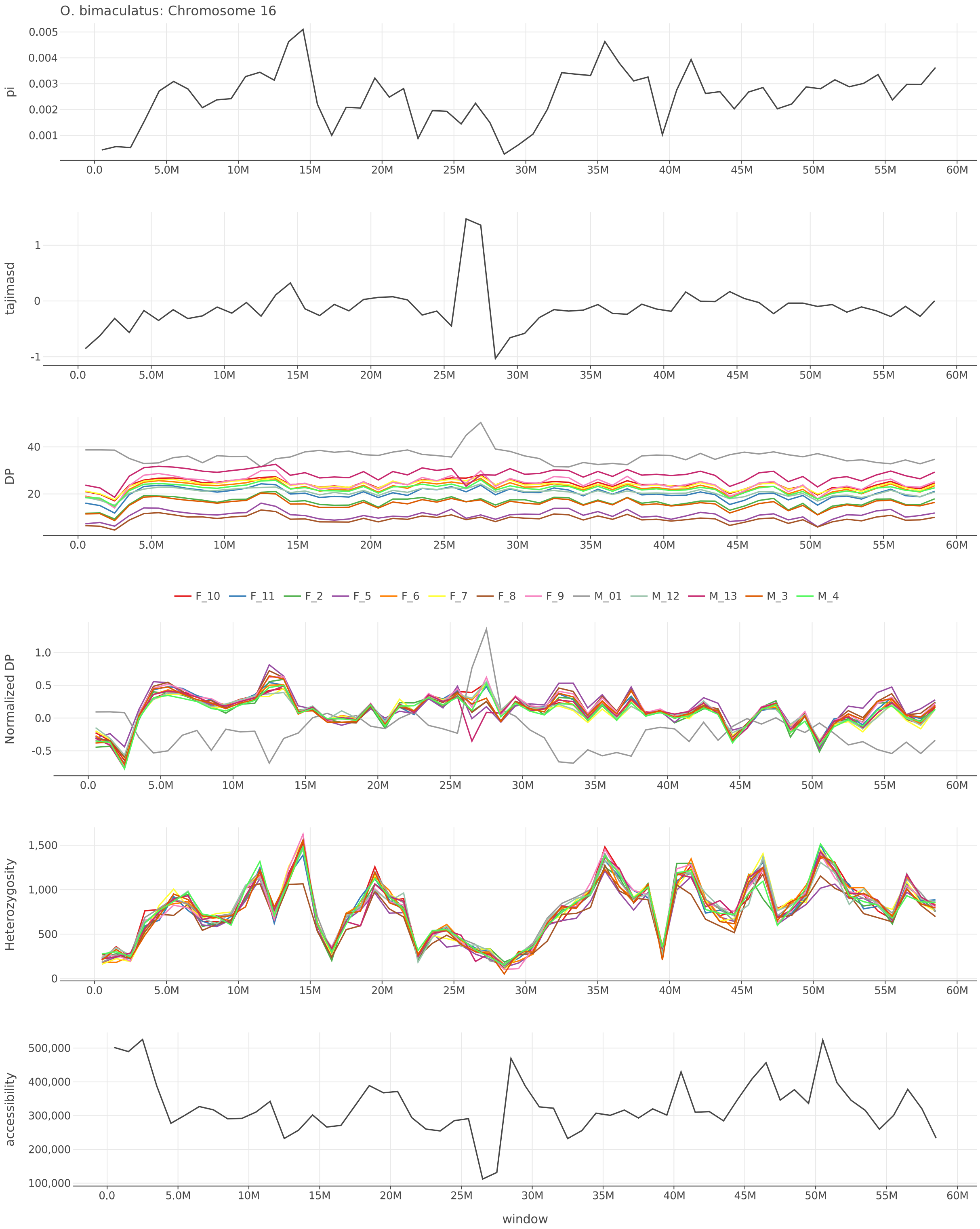


Figure S50: $\pi$, Tajima's $D$, depth, normalized depth, heterozygosity, and accessibility across chromosome 16 of *O. bimaculatus*.

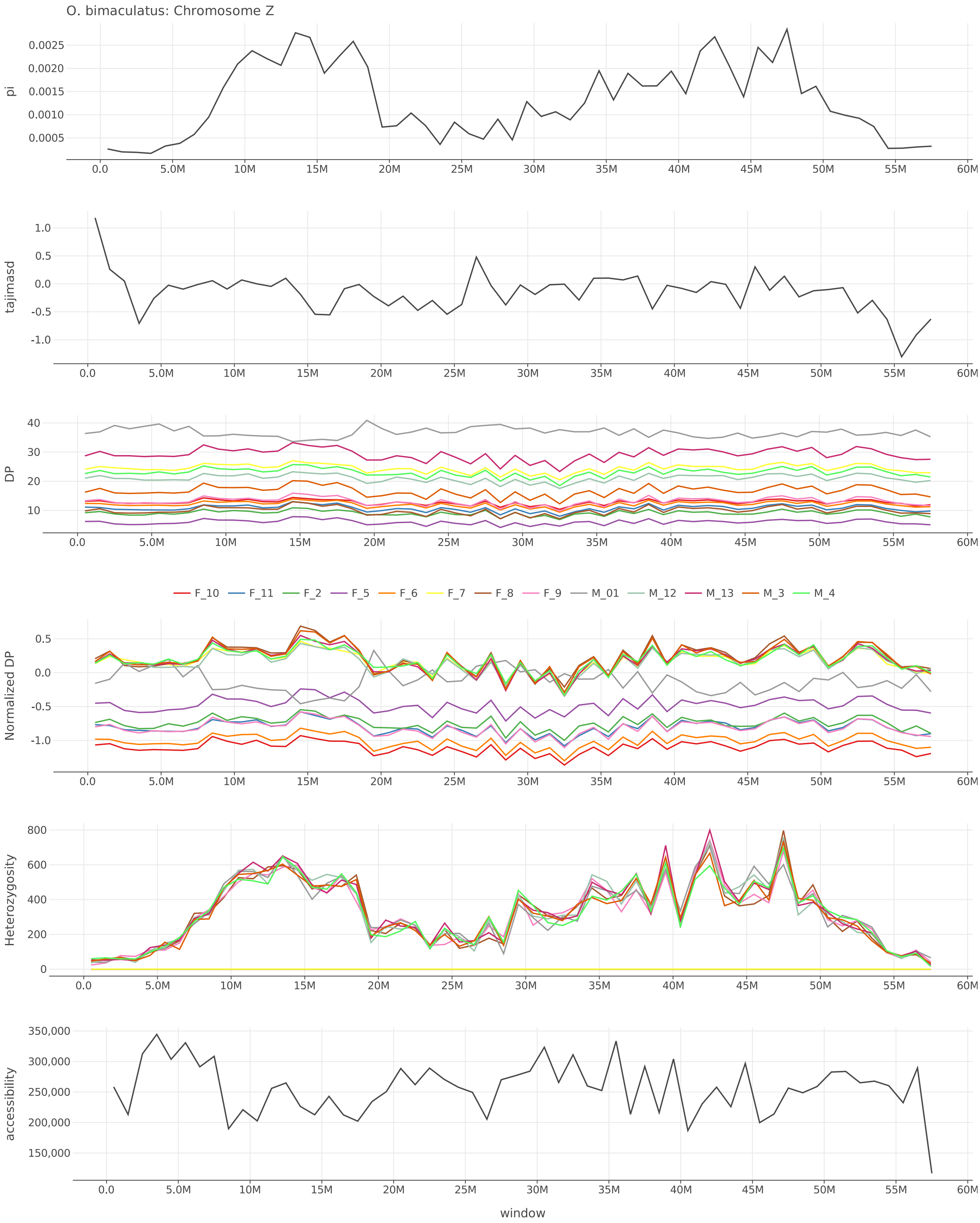


Figure S51: $\pi$, Tajima's $D$, depth, normalized depth, heterozygosity, and accessibility across chromosome Z of *O. bimaculatus*.

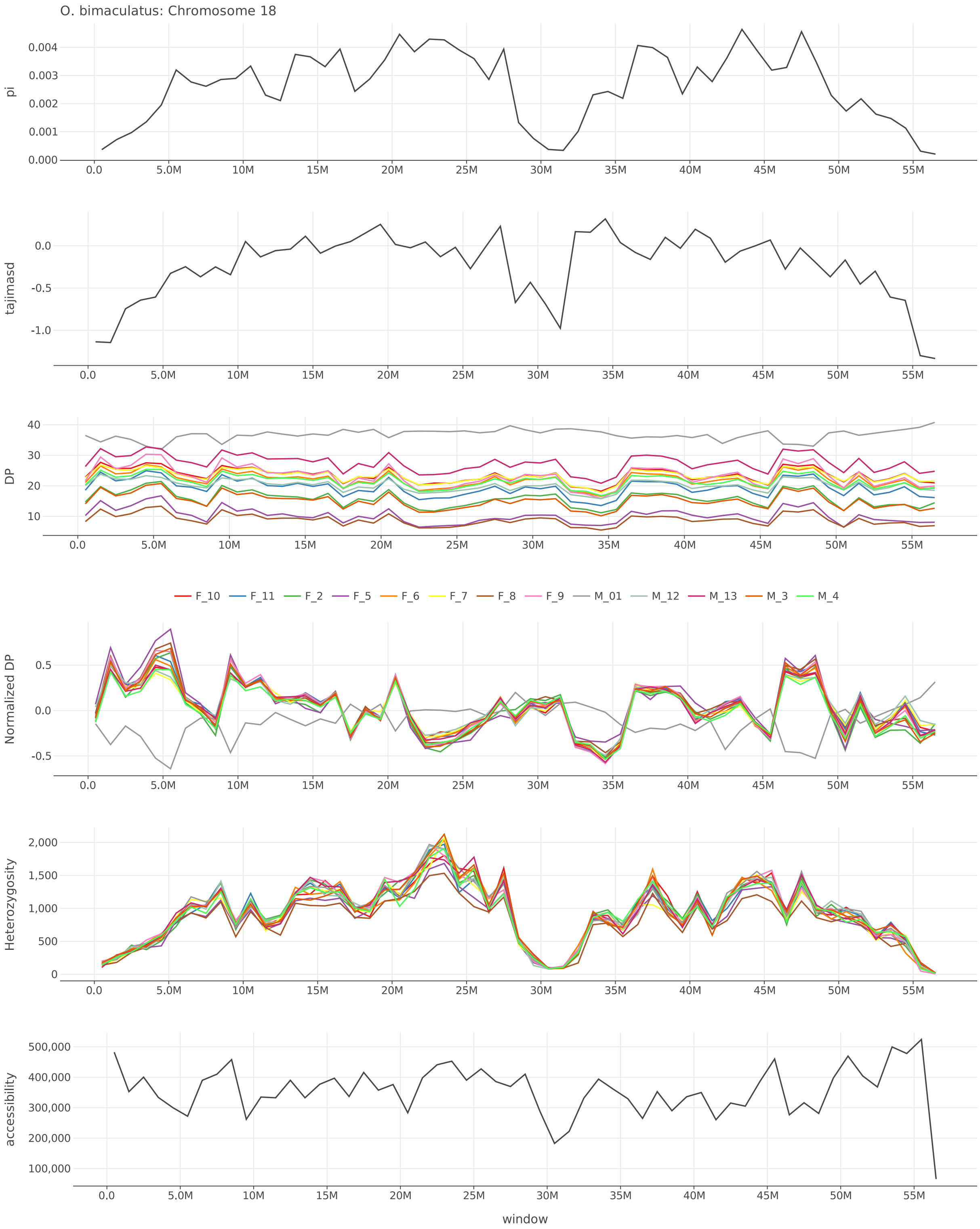


Figure S52: $\pi$, Tajima's $D$, depth, normalized depth, heterozygosity, and accessibility across chromosome 18 of *O. bimaculatus*.

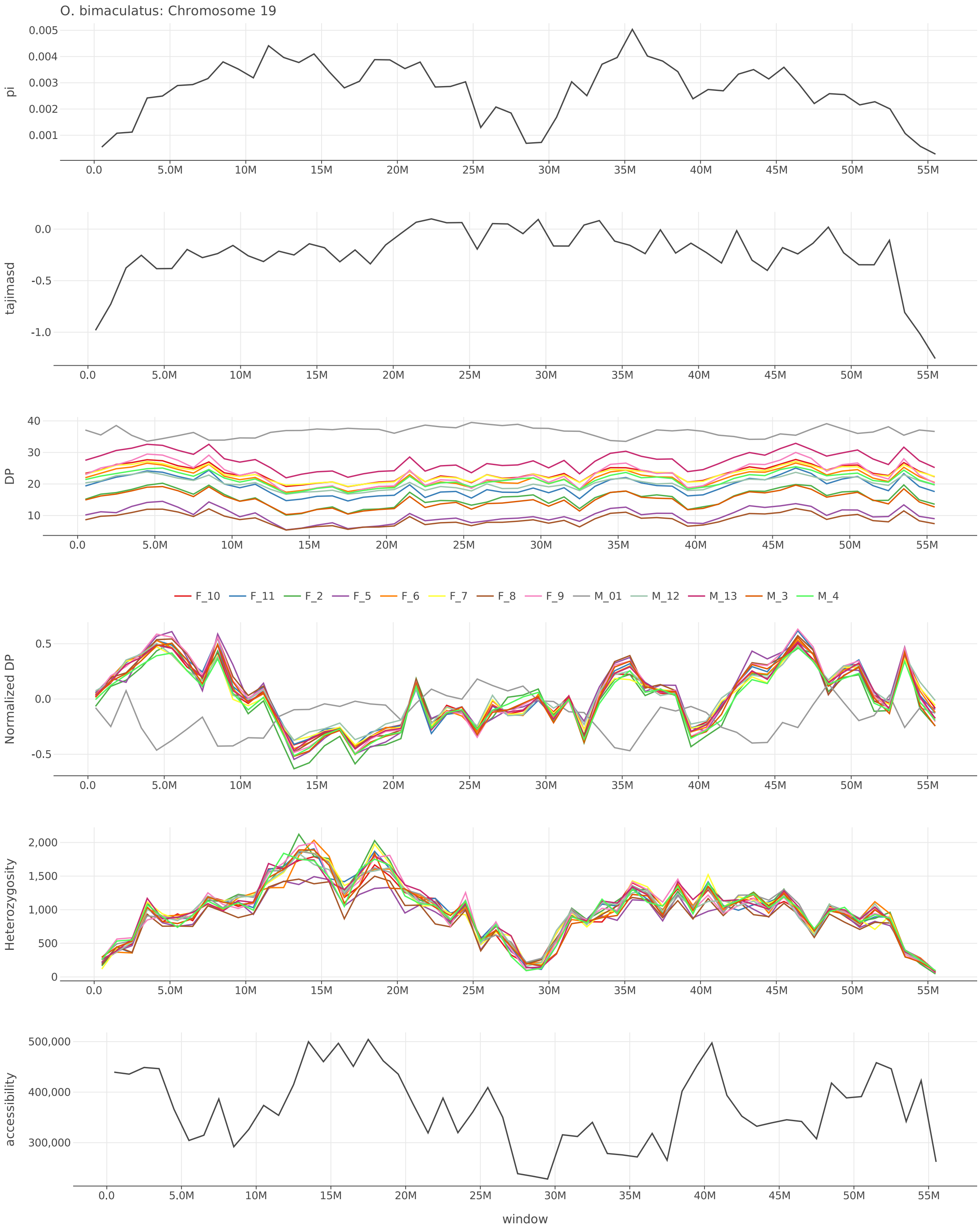


Figure S53: $\pi$, Tajima's $D$, depth, normalized depth, heterozygosity, and accessibility across chromosome 19 of *O. bimaculatus*.

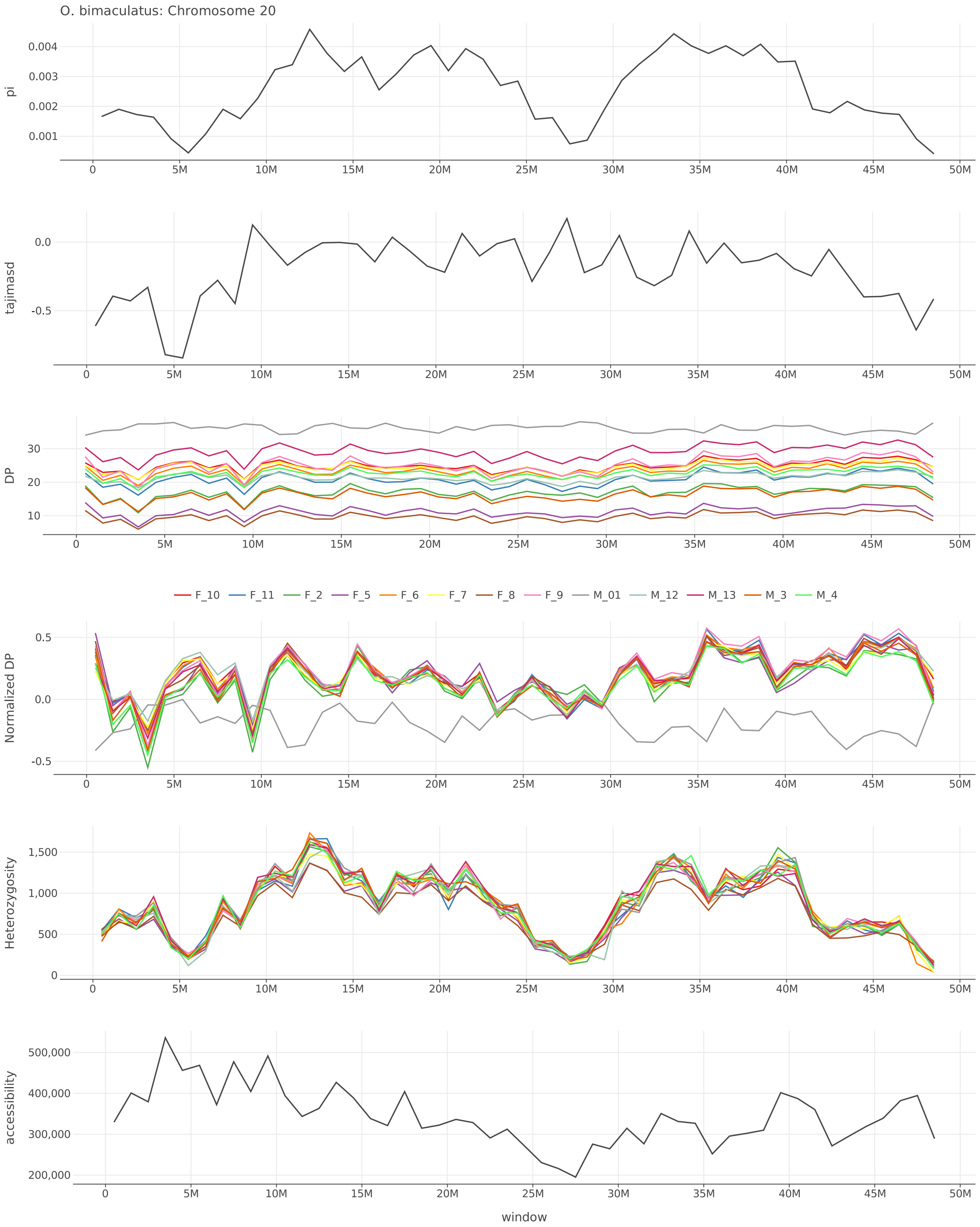


Figure S54: $\pi$, Tajima's $D$, depth, normalized depth, heterozygosity, and accessibility across chromosome 20 of *O. bimaculatus*.

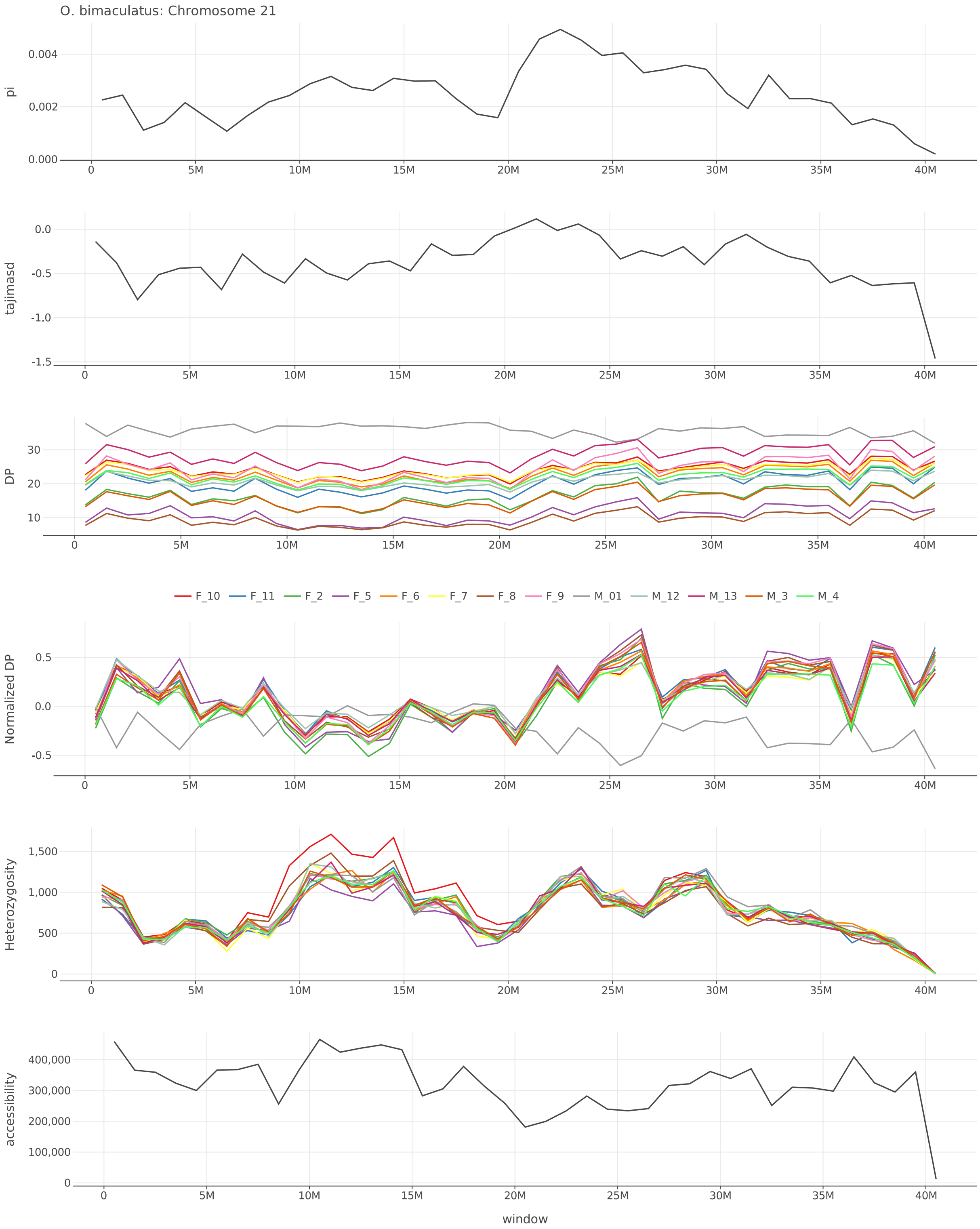


Figure S55: $\pi$, Tajima's $D$, depth, normalized depth, heterozygosity, and accessibility across chromosome 21 of *O. bimaculatus*.

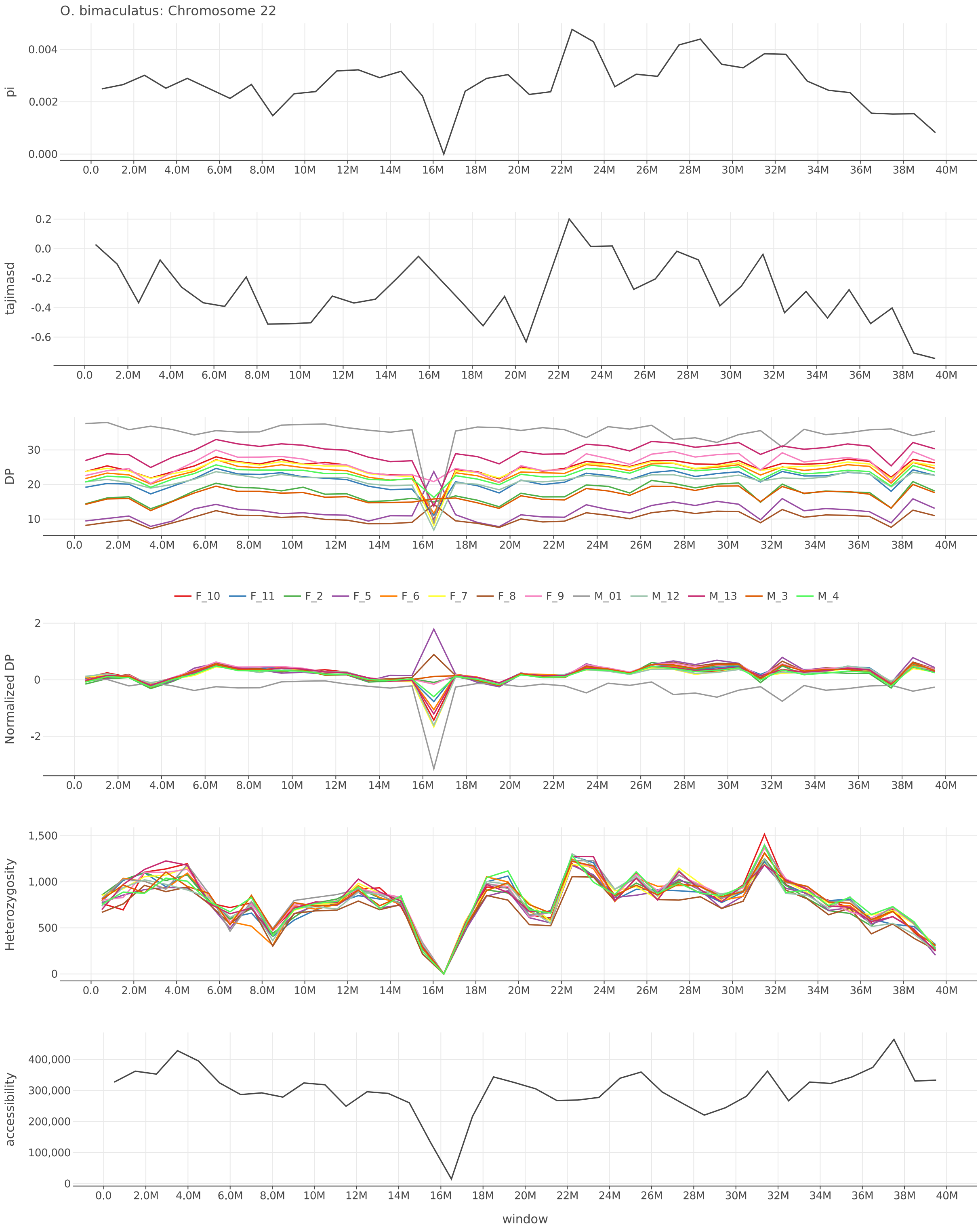


Figure S56: $\pi$, Tajima's $D$, depth, normalized depth, heterozygosity, and accessibility across chromosome 22 of *O. bimaculatus*.

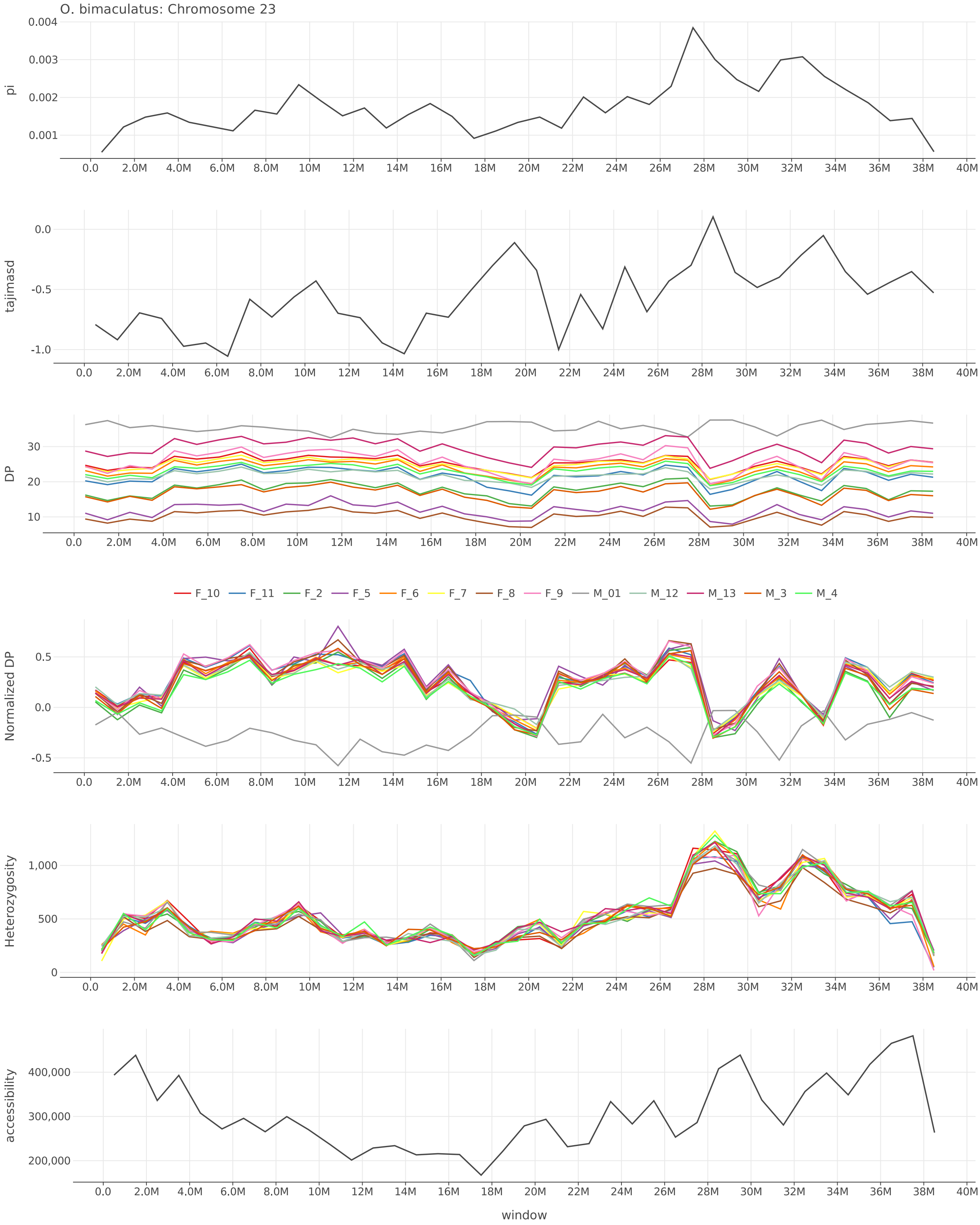


Figure S57: $\pi$, Tajima's $D$, depth, normalized depth, heterozygosity, and accessibility across chromosome 23 of *O. bimaculatus*.

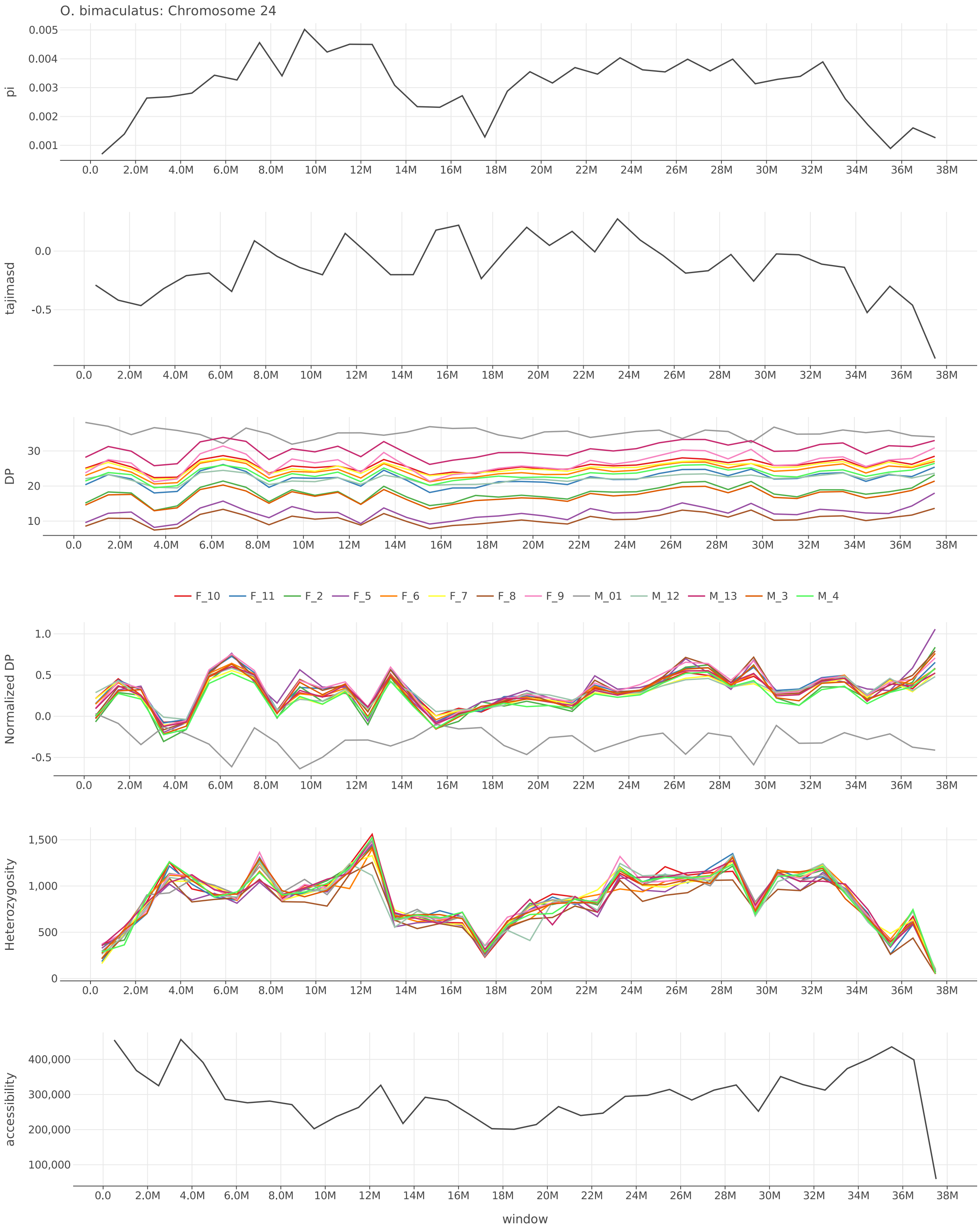


Figure S58: $\pi$, Tajima's $D$, depth, normalized depth, heterozygosity, and accessibility across chromosome 24 of *O. bimaculatus*.

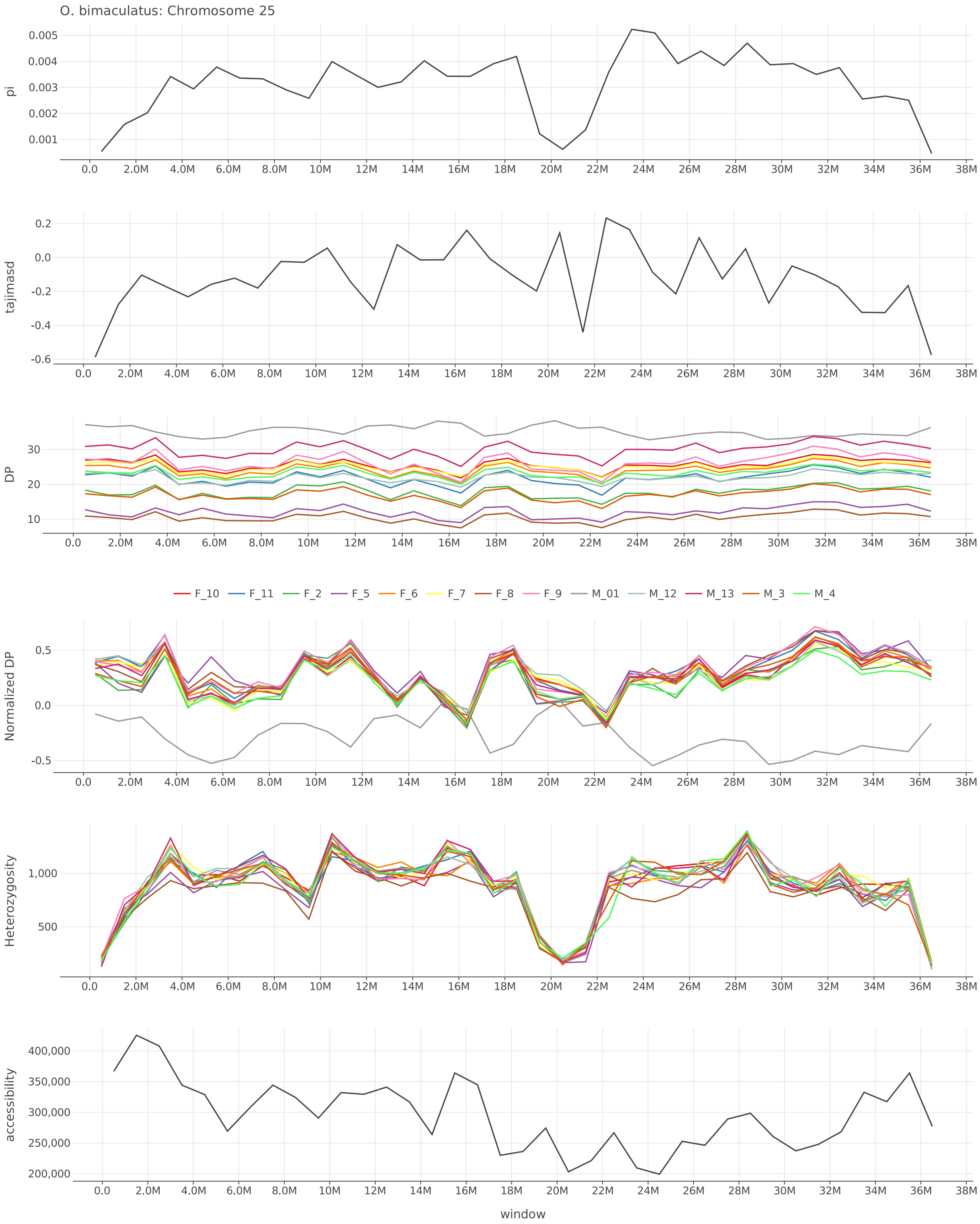


Figure S59: $\pi$, Tajima's $D$, depth, normalized depth, heterozygosity, and accessibility across chromosome 25 of *O. bimaculatus*.

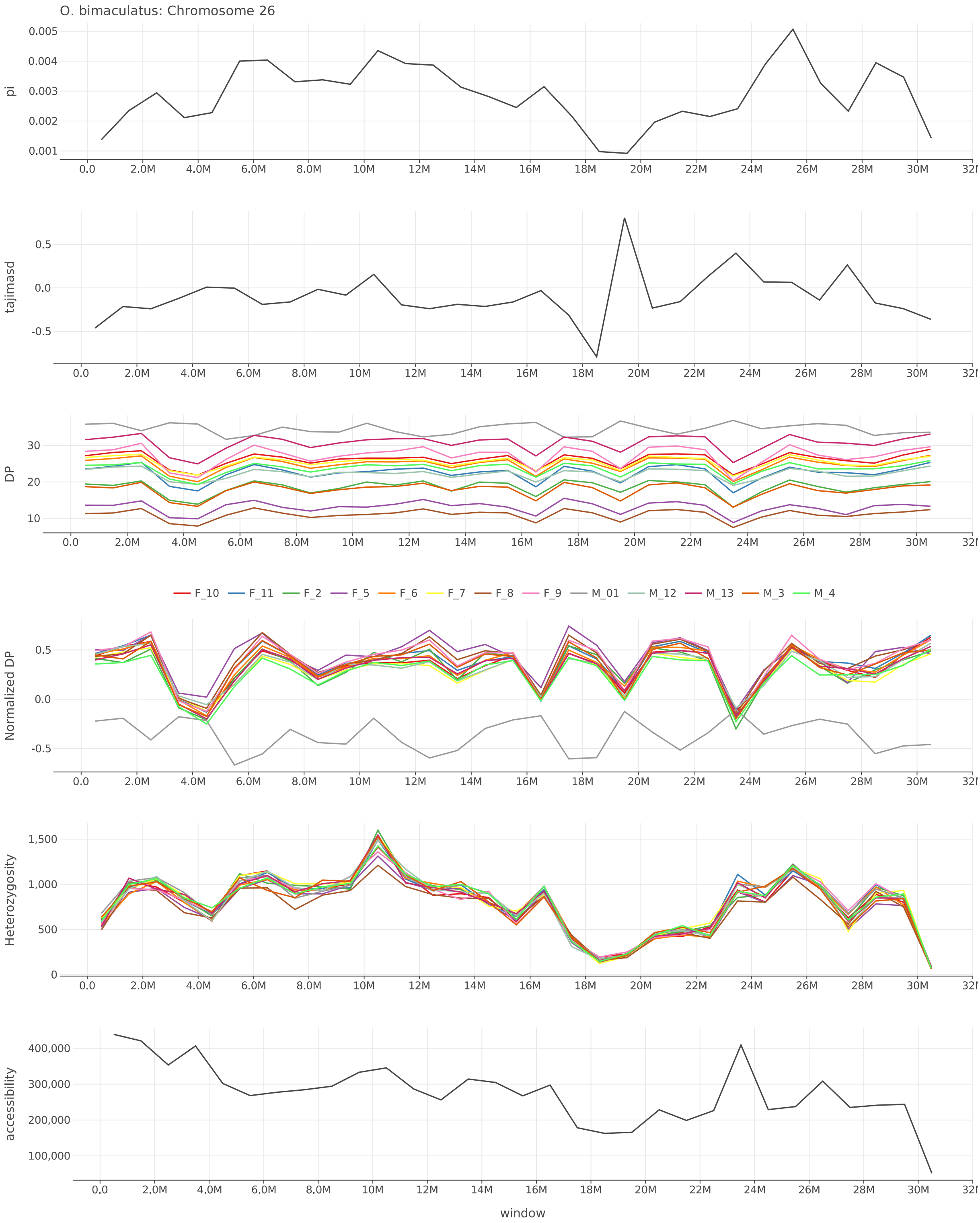


Figure S60: $\pi$, Tajima's $D$, depth, normalized depth, heterozygosity, and accessibility across chromosome 26 of *O. bimaculatus*.

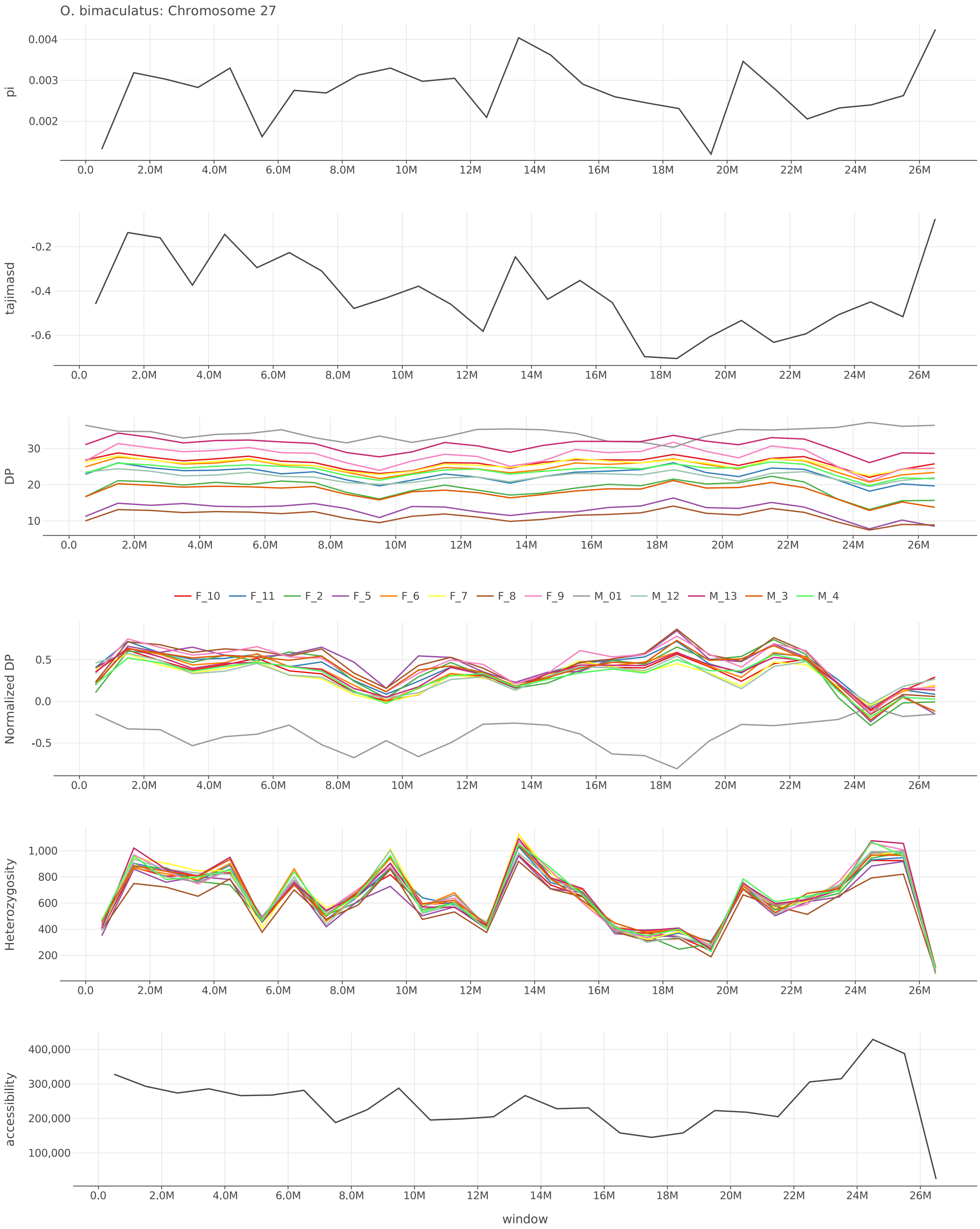


Figure S61: $\pi$, Tajima's $D$, depth, normalized depth, heterozygosity, and accessibility across chromosome 27 of *O. bimaculatus*.

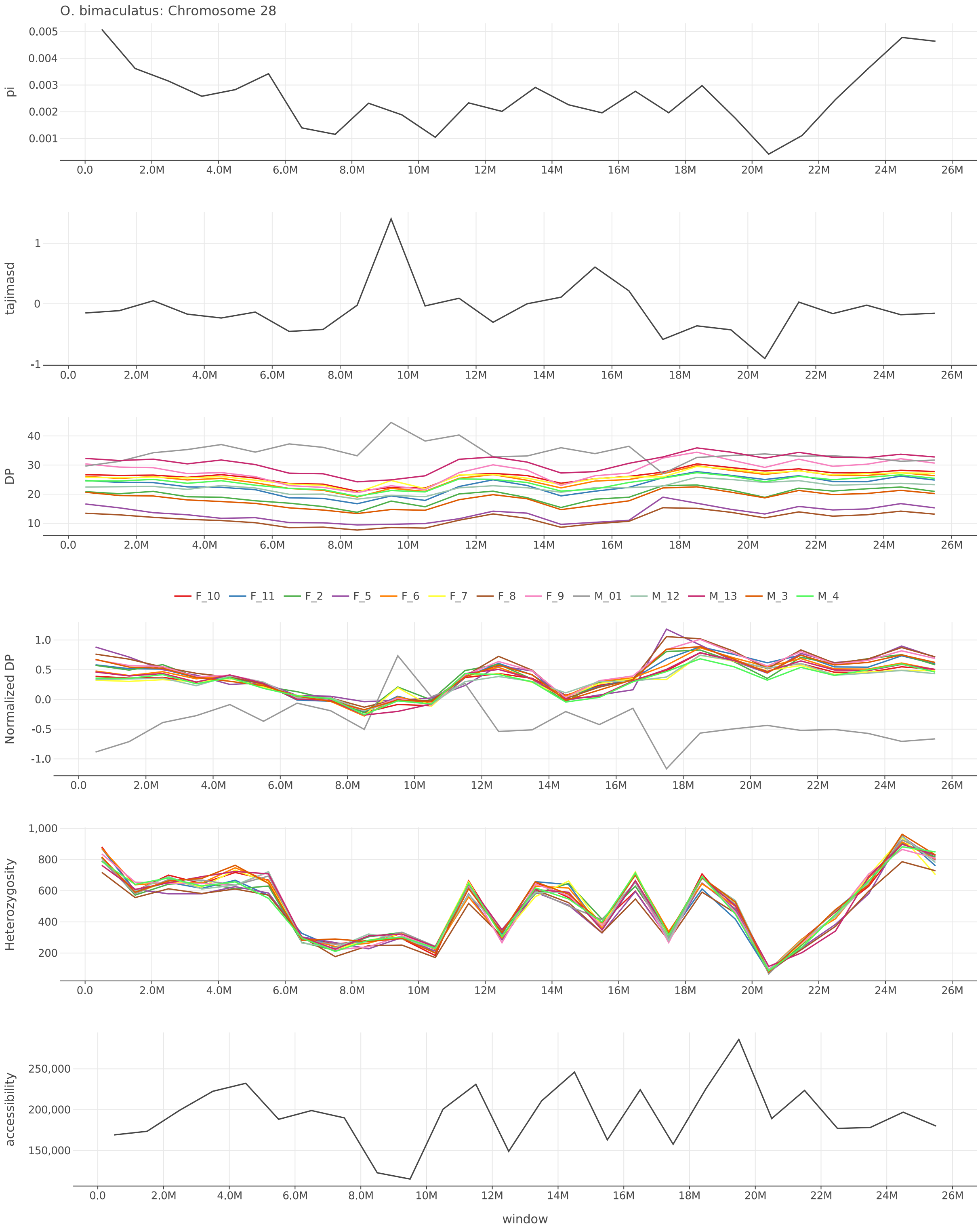


Figure S62: $\pi$, Tajima's $D$, depth, normalized depth, heterozygosity, and accessibility across chromosome 28 of *O. bimaculatus*.

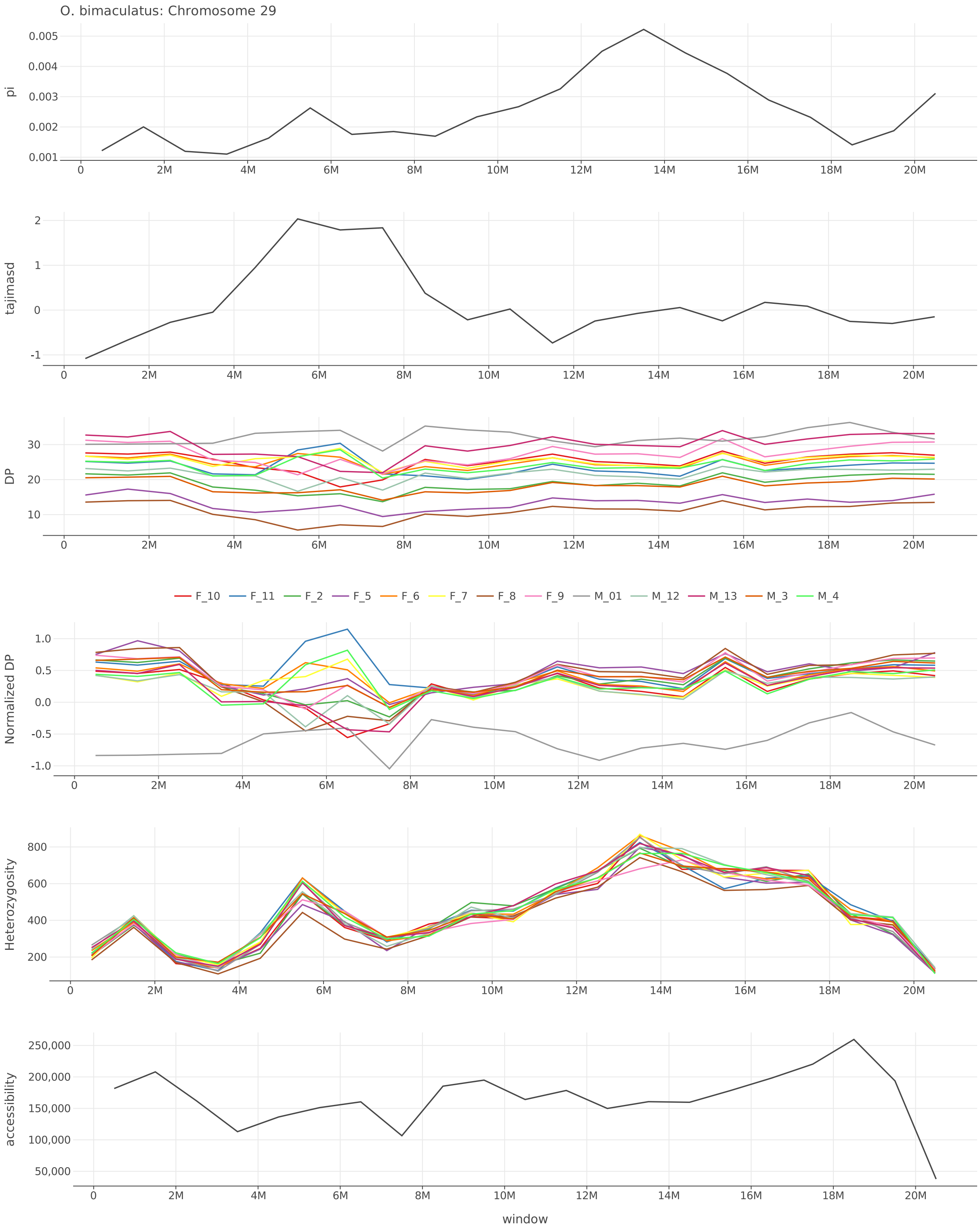


Figure S63: $\pi$, Tajima's $D$, depth, normalized depth, heterozygosity, and accessibility across chromosome 29 of *O. bimaculatus*.

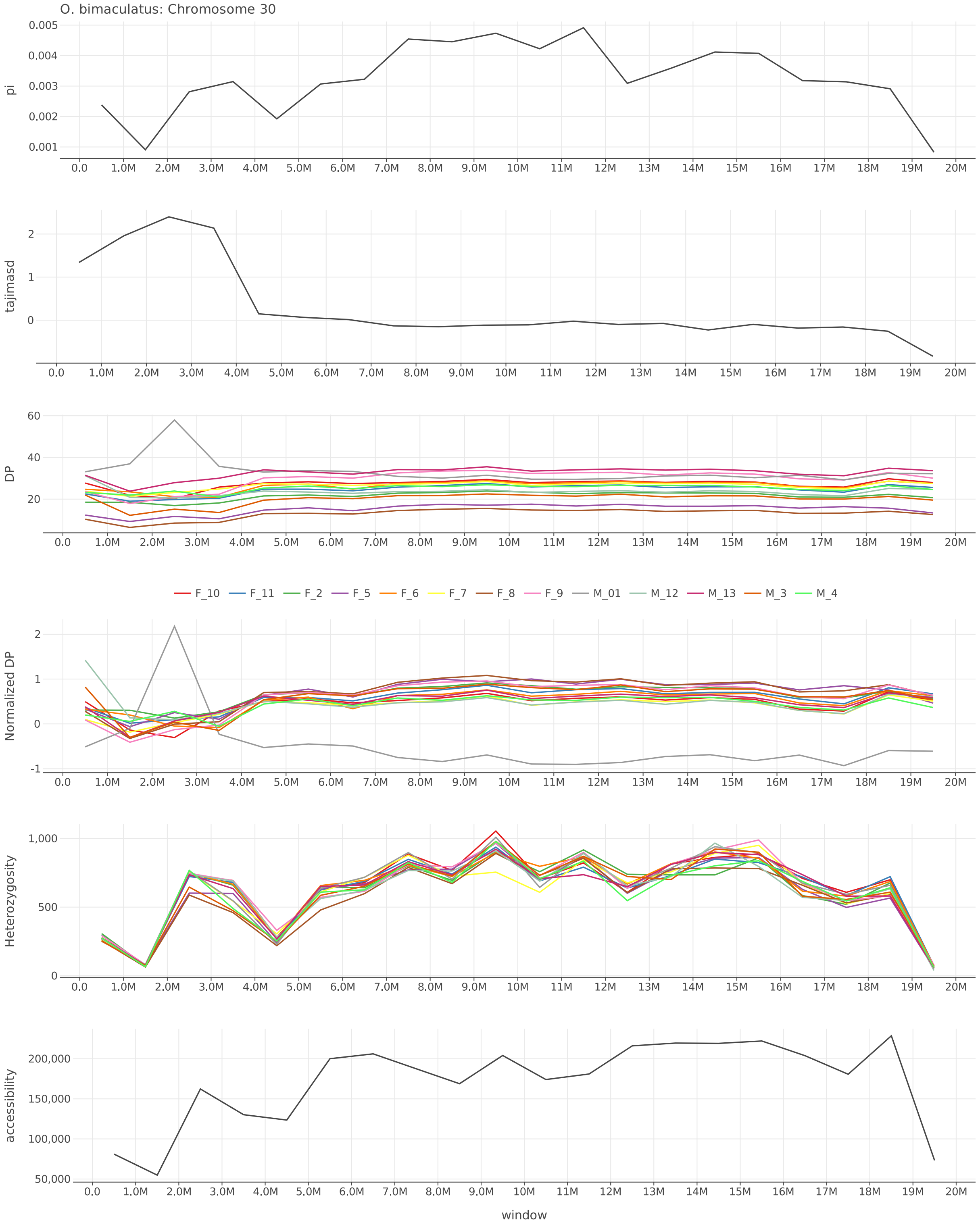


Figure S64: $\pi$, Tajima's $D$, depth, normalized depth, heterozygosity, and accessibility across chromosome 30 of *O. bimaculatus*.

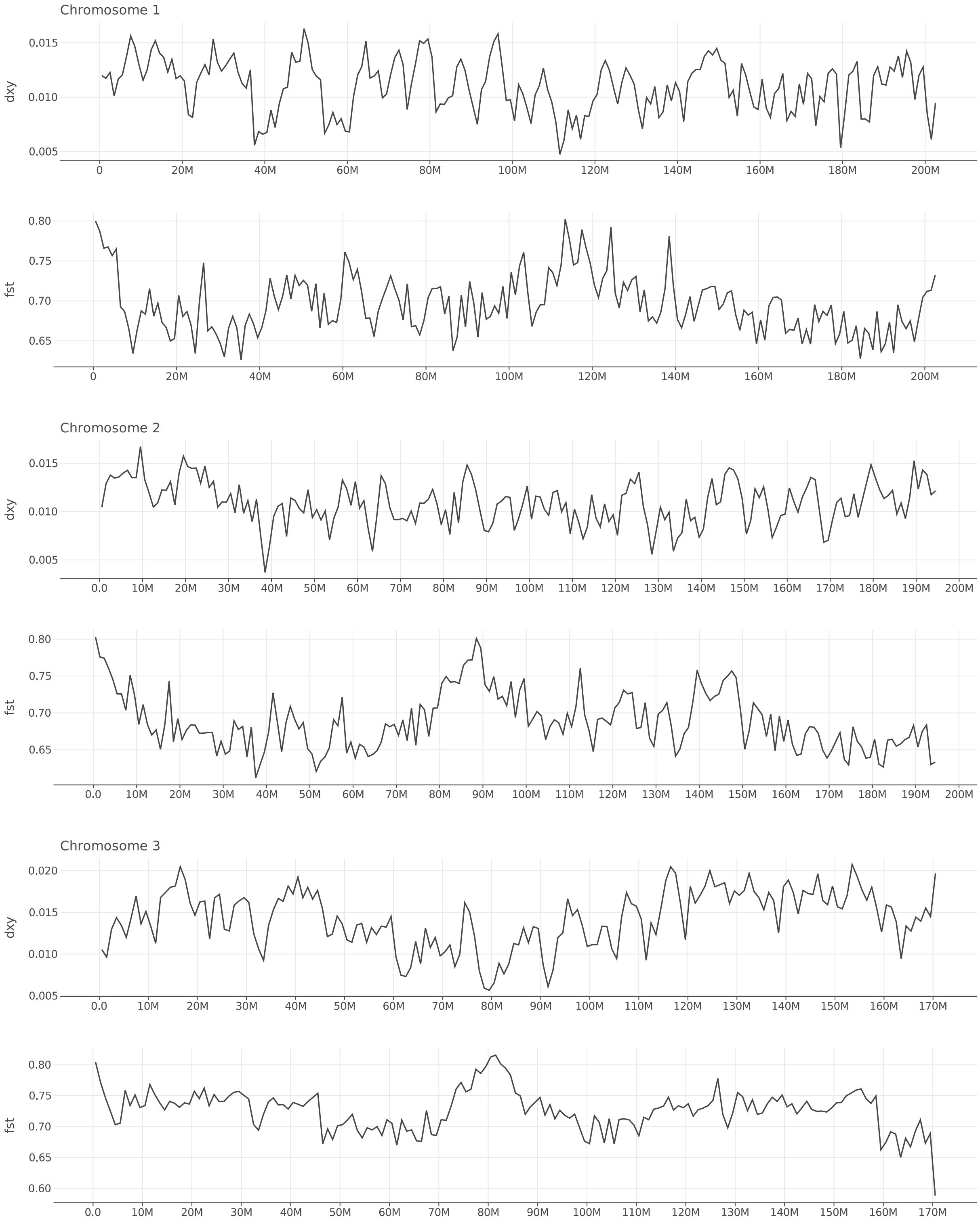


Figure S65: $d_{XY}$ and $F_{ST}$ between *O. bimaculoides* and *O. bimaculatus* along chromosomes 1–3.

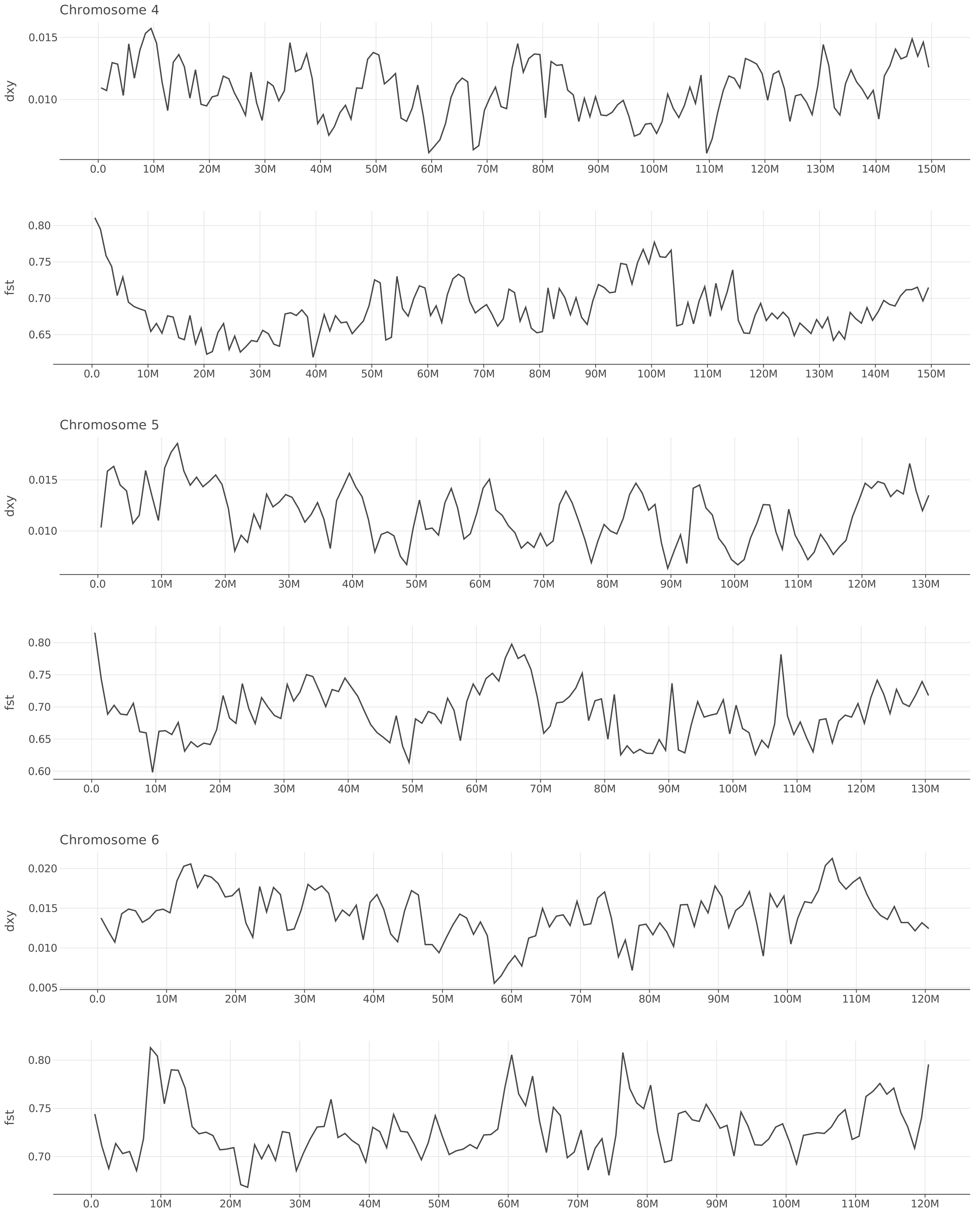


Figure S66: $d_{XY}$ and $F_{ST}$ between *O. bimaculoides* and *O. bimaculatus* along chromosomes 4–6.

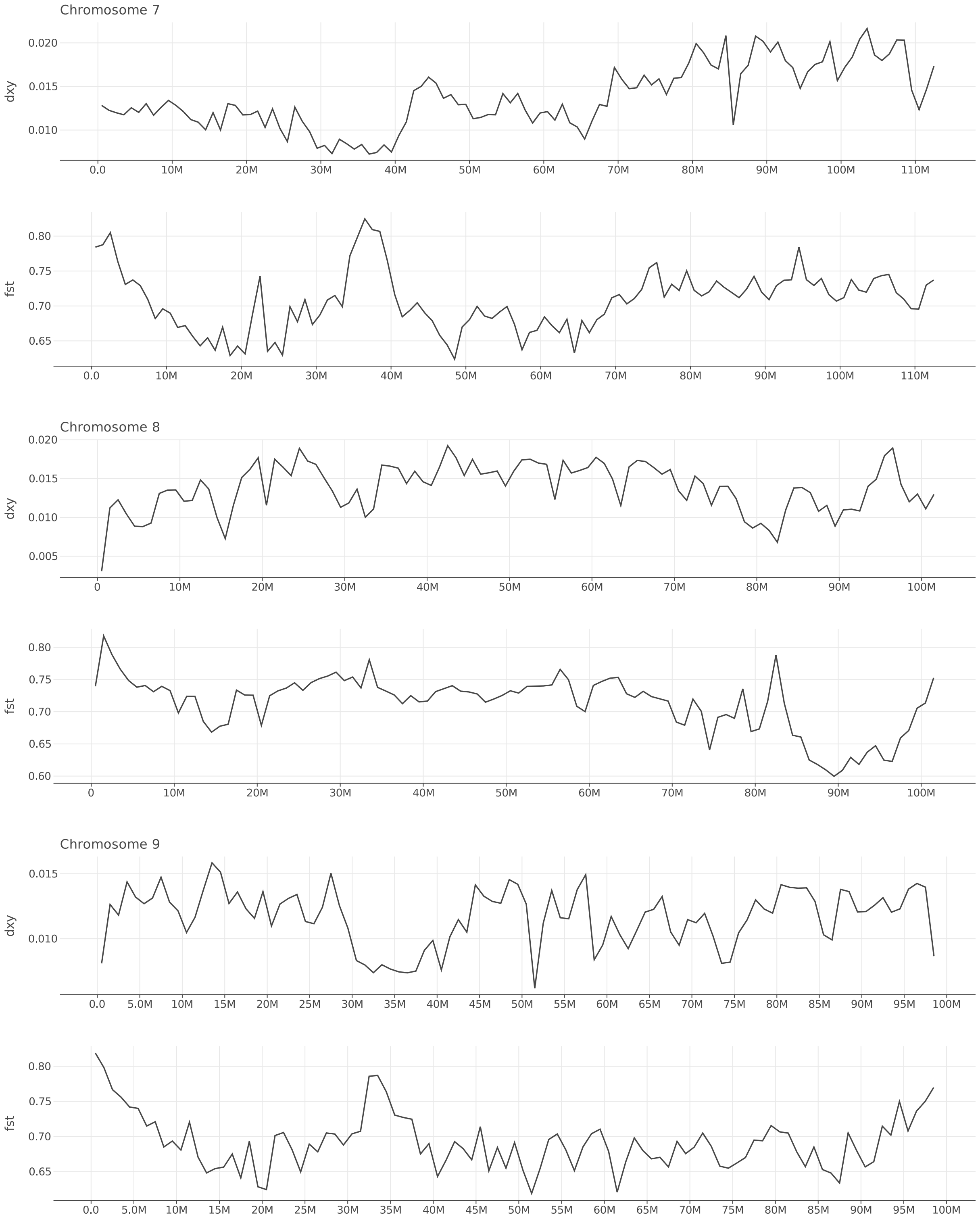


Figure S67: $d_{XY}$ and $F_{ST}$ between *O. bimaculoides* and *O. bimaculatus* along chromosomes 7–9.

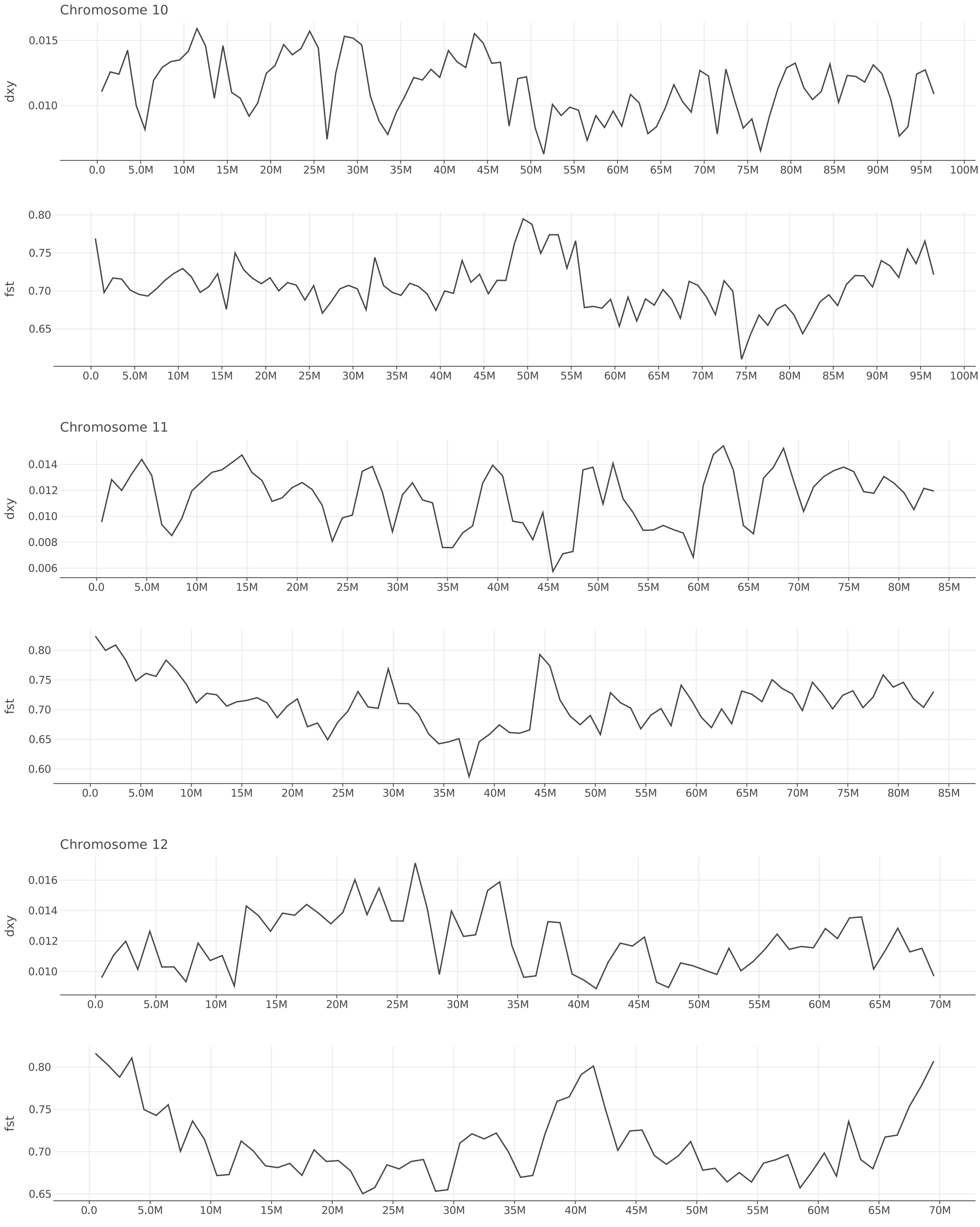


Figure S68: $d_{XY}$ and $F_{ST}$ between *O. bimaculoides* and *O. bimaculatus* along chromosomes 10–12.

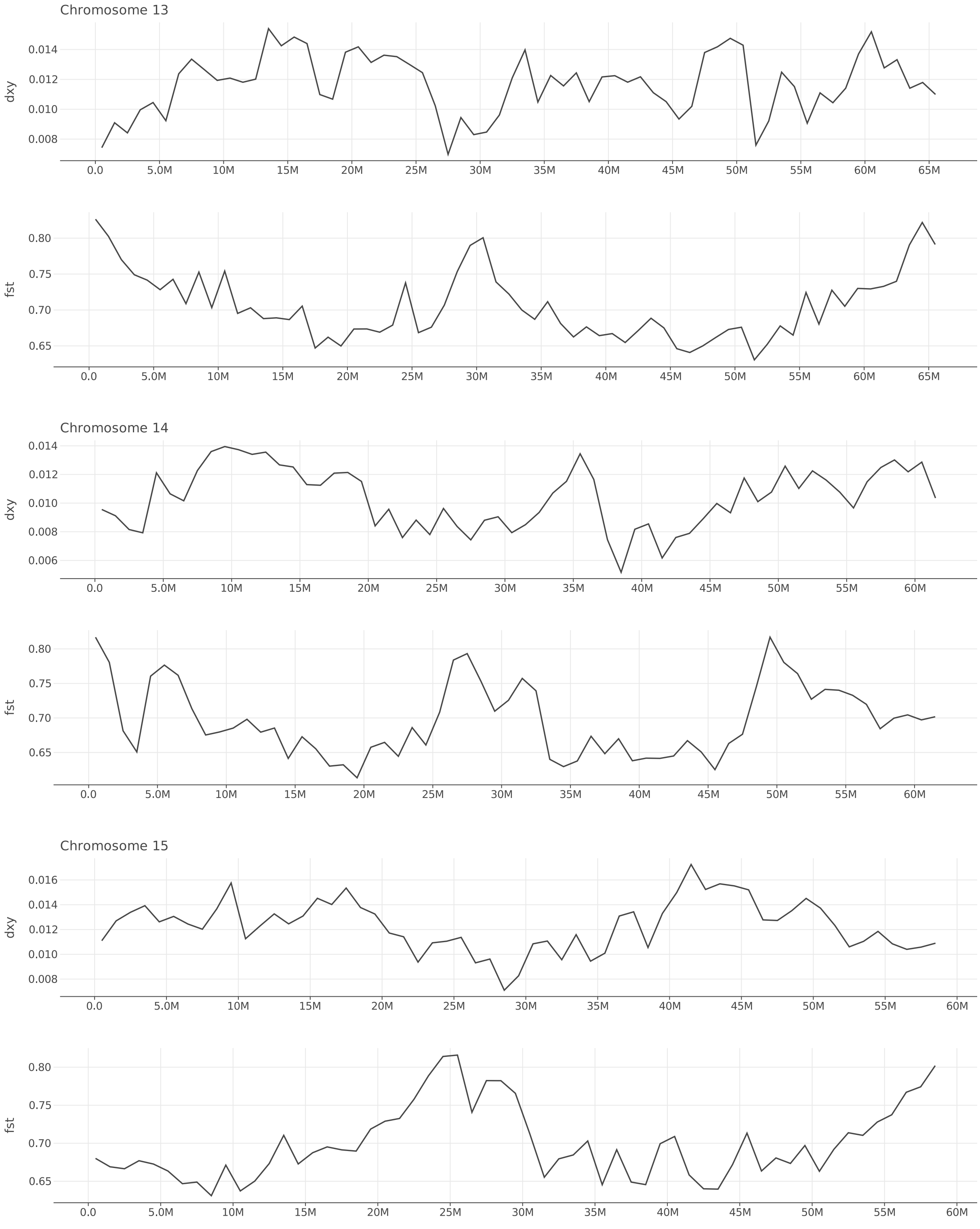


Figure S69: $d_{XY}$ and $F_{ST}$ between *O. bimaculoides* and *O. bimaculatus* along chromosomes 13–15.

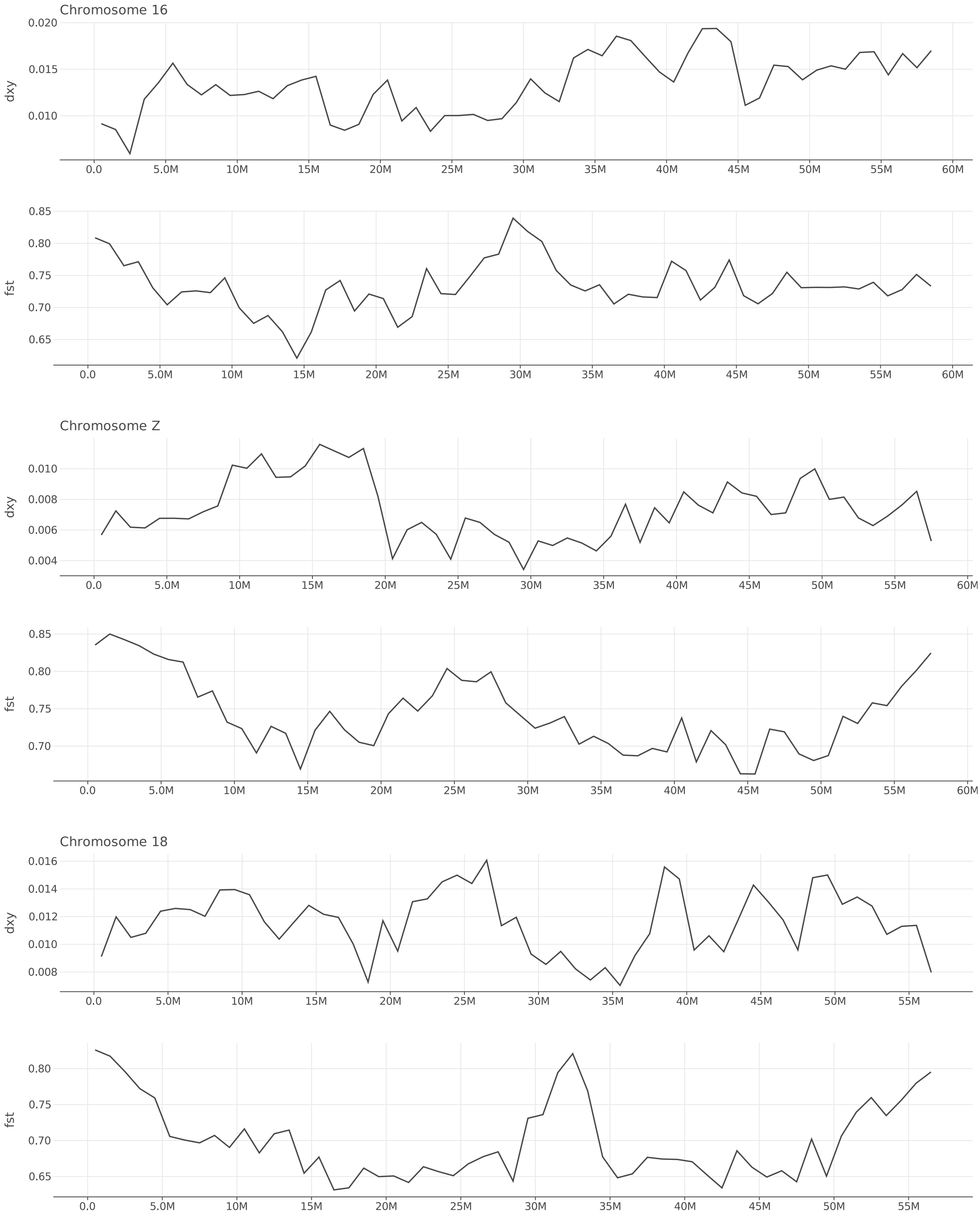


Figure S70: $d_{XY}$ and $F_{ST}$ between *O. bimaculoides* and *O. bimaculatus* along chromosomes 16–18.

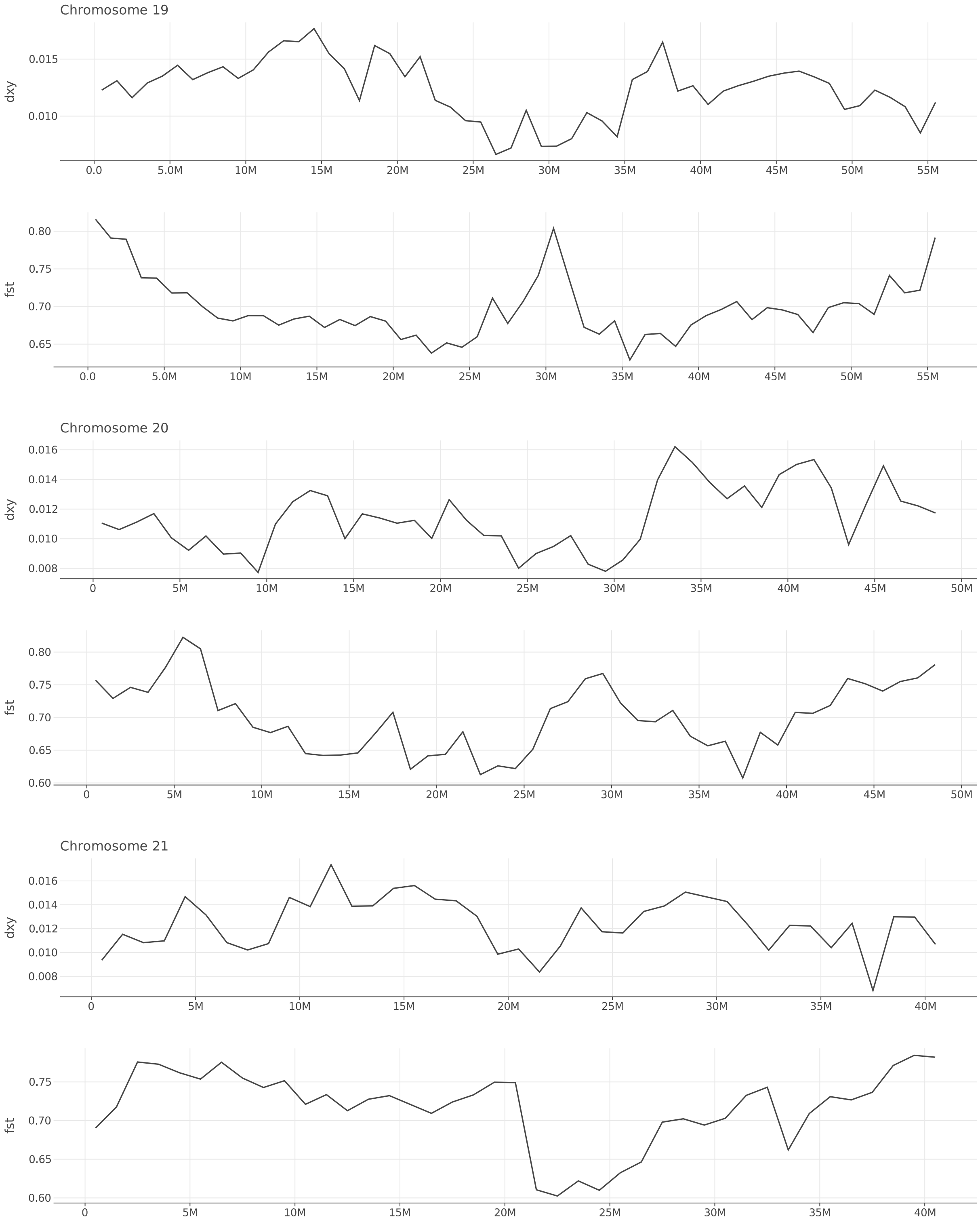


Figure S71: $d_{XY}$ and $F_{ST}$ between *O. bimaculoides* and *O. bimaculatus* along chromosomes 19–21.

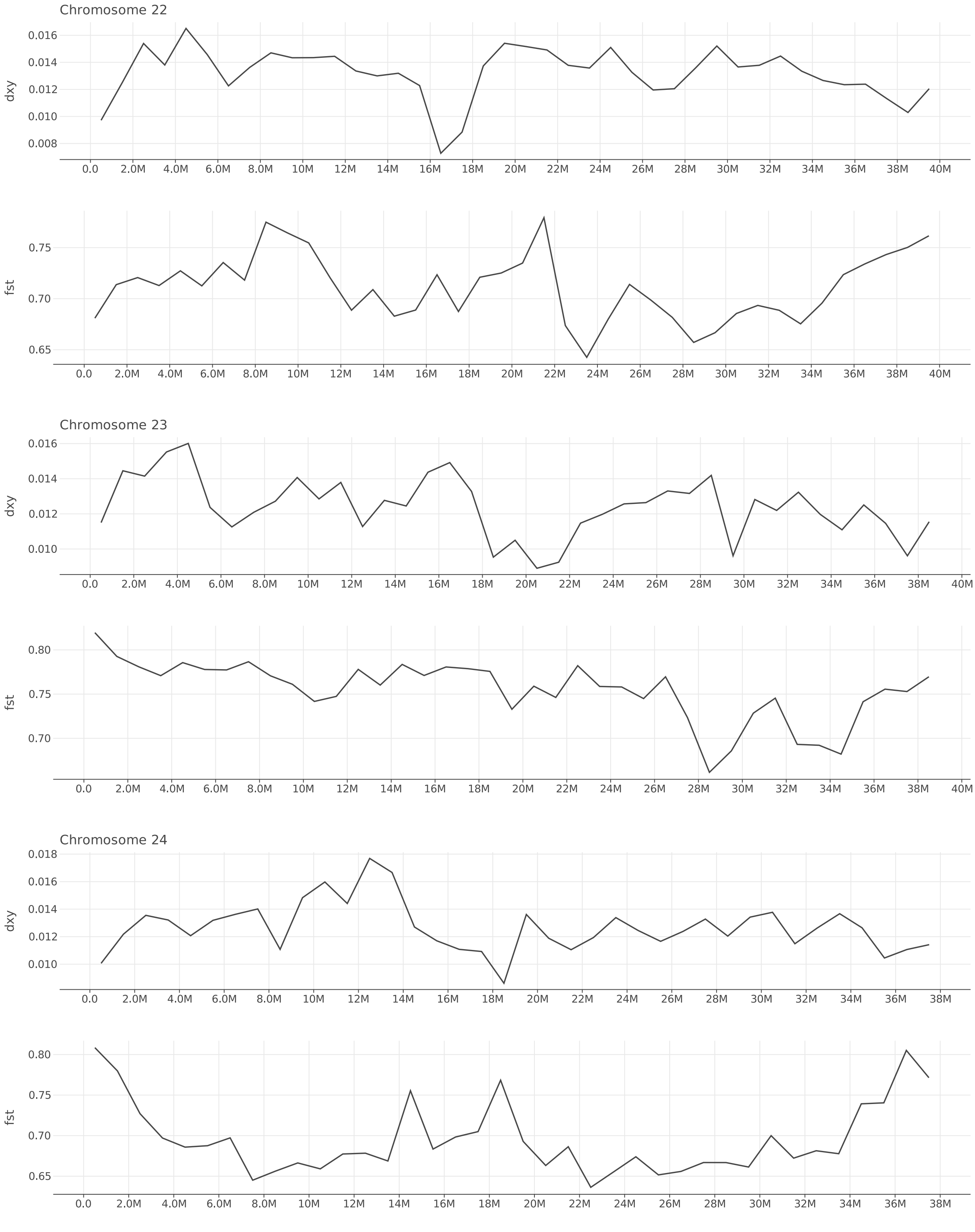


Figure S72: $d_{XY}$ and $F_{ST}$ between *O. bimaculoides* and *O. bimaculatus* along chromosomes 22–24.

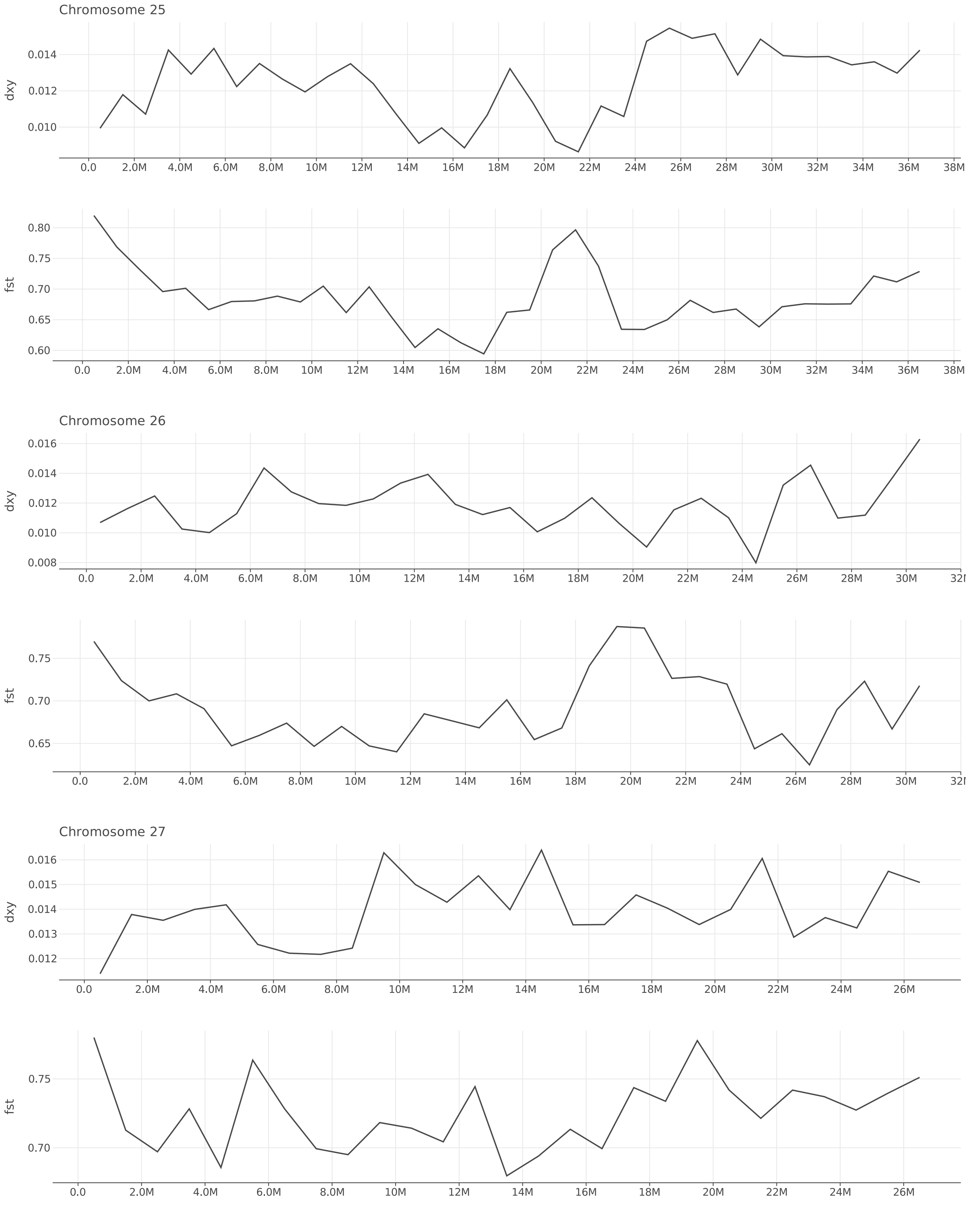


Figure S73: $d_{XY}$ and $F_{ST}$ between *O. bimaculoides* and *O. bimaculatus* along chromosomes 25–27.

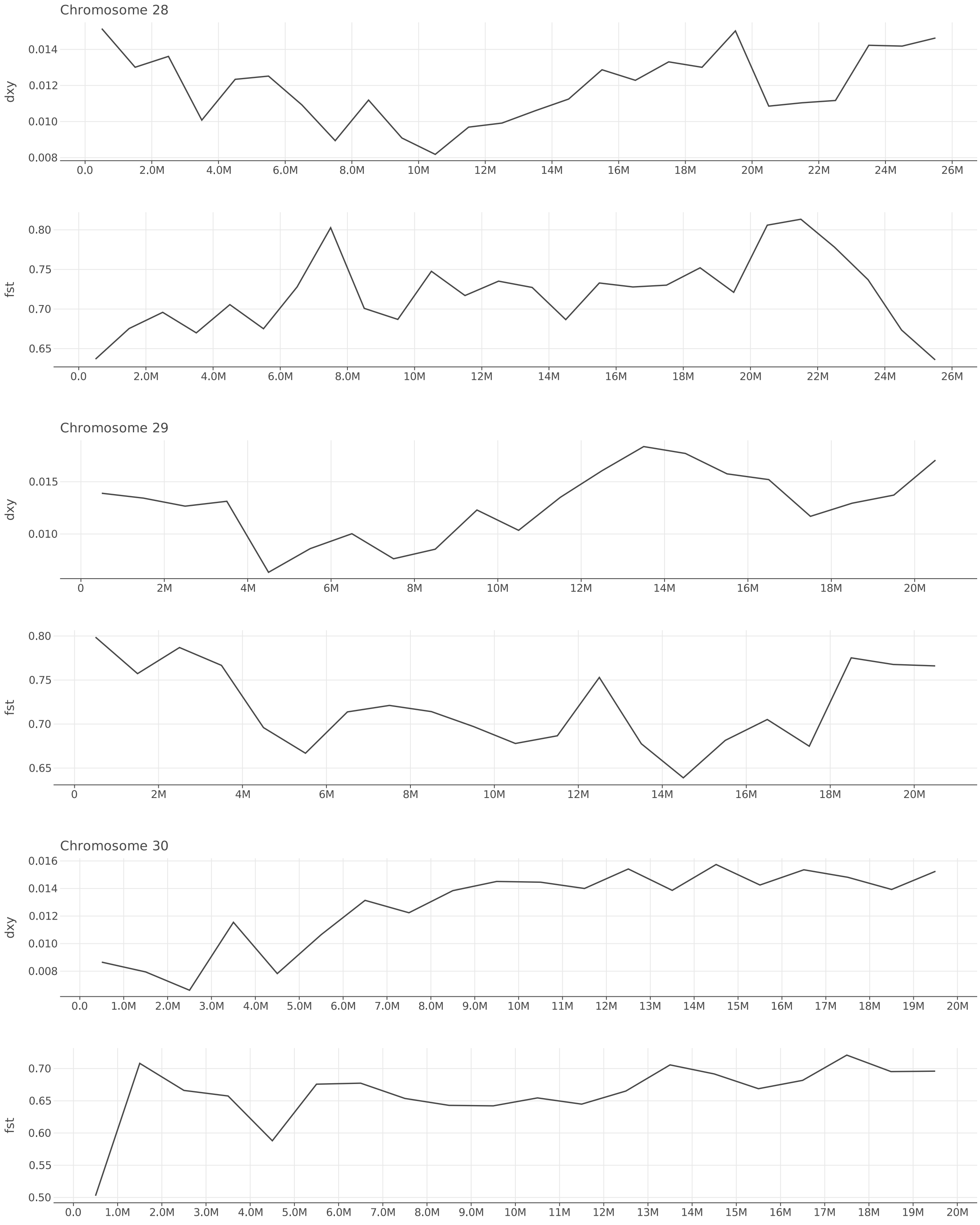


Figure S74: $d_{XY}$ and $F_{ST}$ between *O. bimaculoides* and *O. bimaculatus* along chromosomes 28–30.

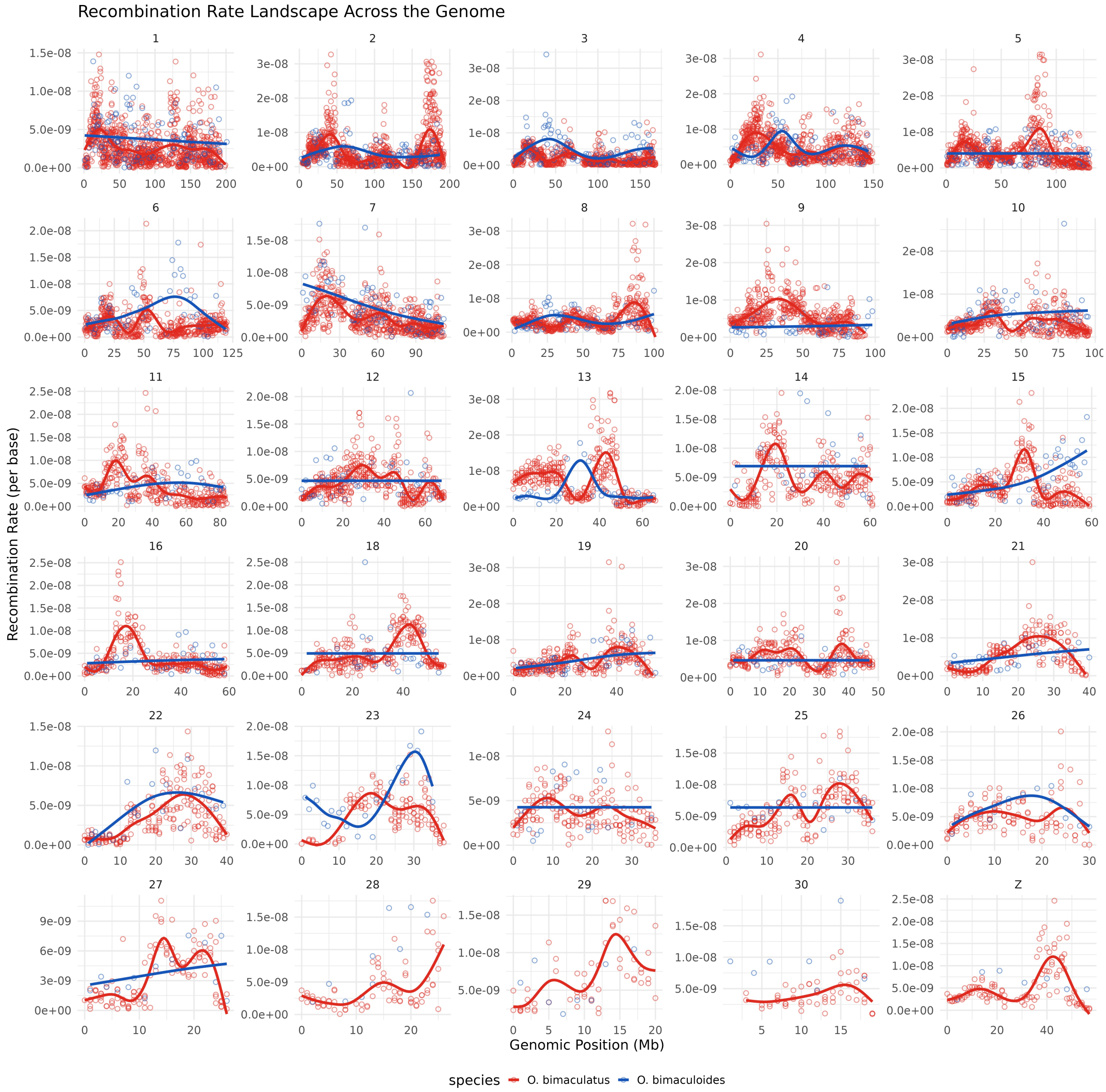


Figure S75: Variation in inferred recombination rate along each chromosome for *O. bimaculatus* (red) and *O. bimaculoides* (blue). Trend lines are best fitting cubic splines inferred using the mgcv R package.

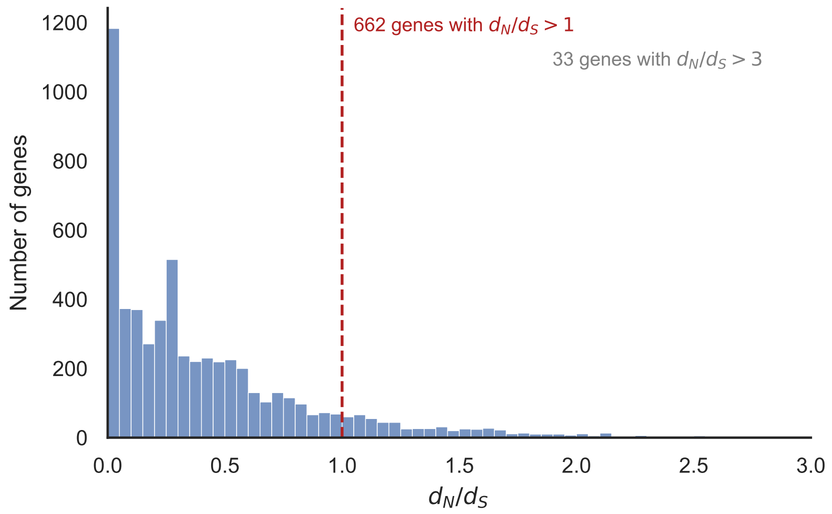


Figure S76: Distribution of per-gene $d_N/d_S$ between *O. bimaculoides* and *O. bimaculatus*, restricted to single-copy orthologs with $d_S > 0$ ($n = 5{,}825$). The dashed line marks $d_N/d_S = 1$; 662 genes (11.4%) fall above this threshold. The distribution is strongly skewed toward low values, consistent with pervasive purifying selection on coding sequence.

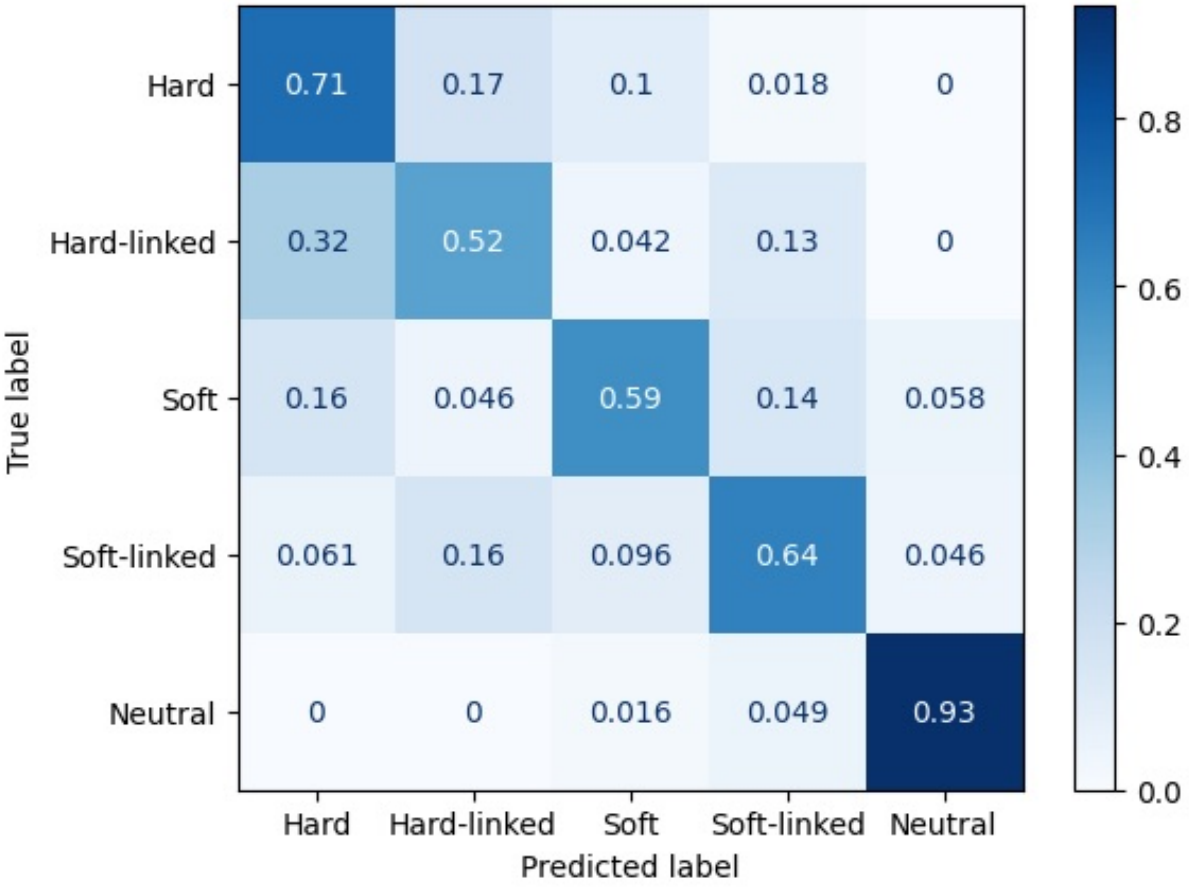


Figure S77: Confusion matrix showing diploS/HIC classification performance simulated under *O. bimaculatus* demographic history and sample size.

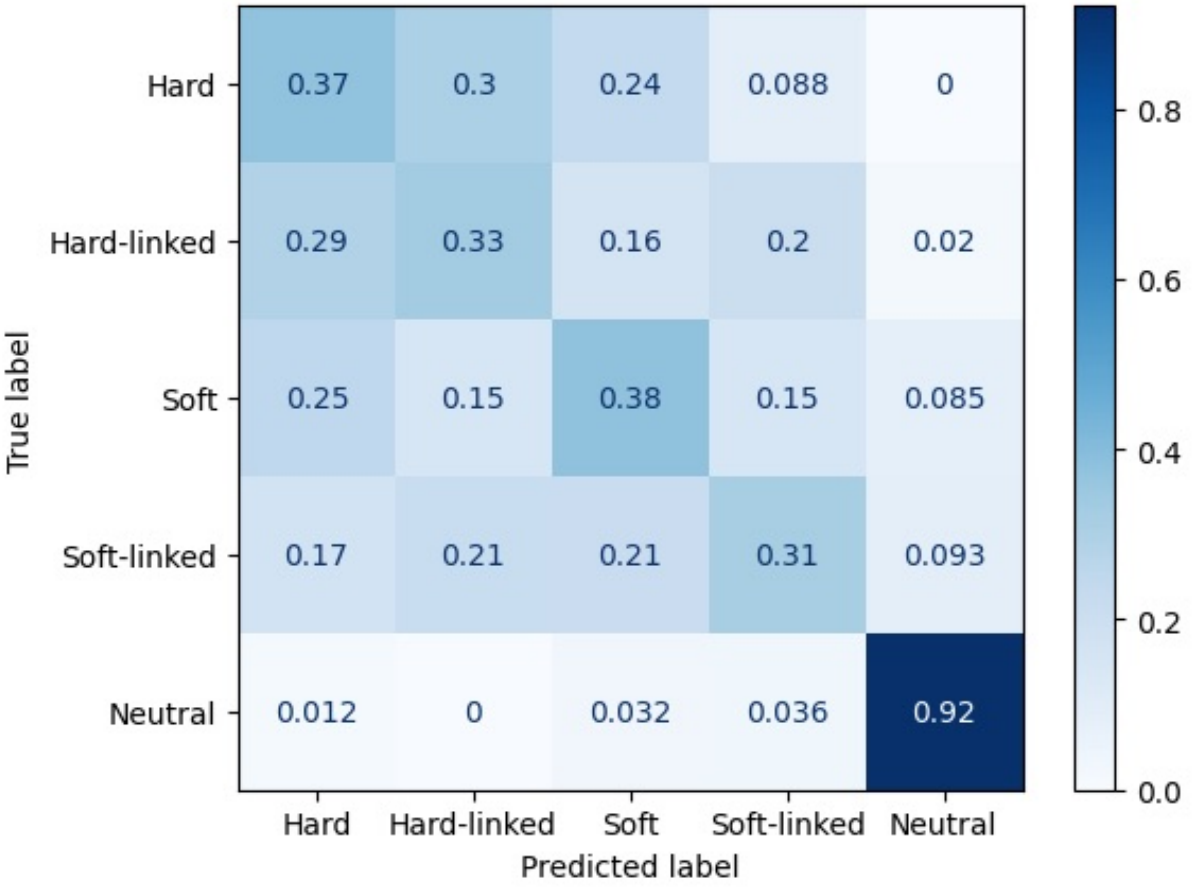


Figure S78: Confusion matrix showing diploS/HIC classification performance simulated under *O. bimaculoides* demographic history and sample size.

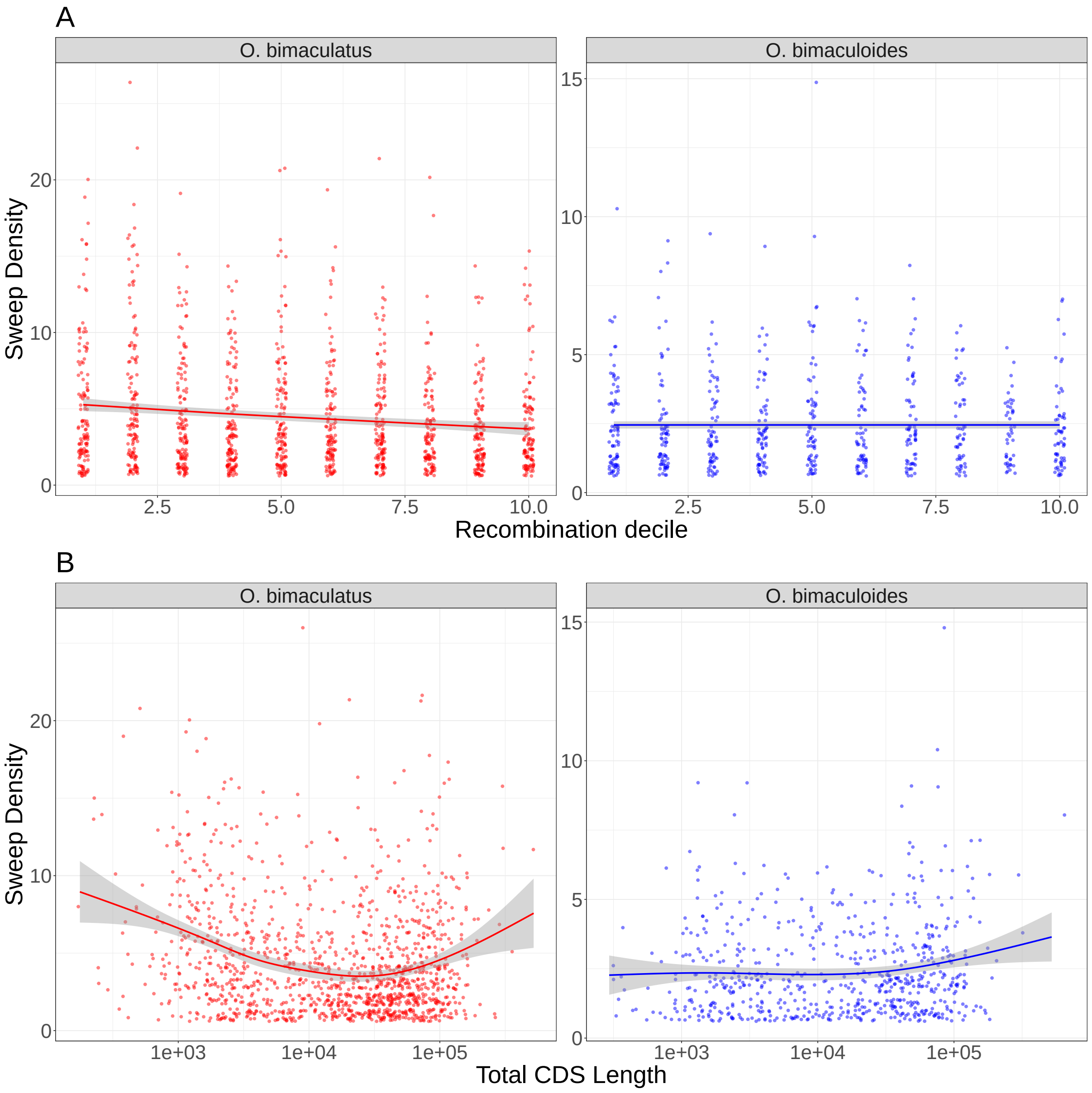


Figure S79: Density of inferred selective sweeps compared to deciles of recombination rate for *O. bimaculatus* (red) and *O. bimaculoides* (blue).

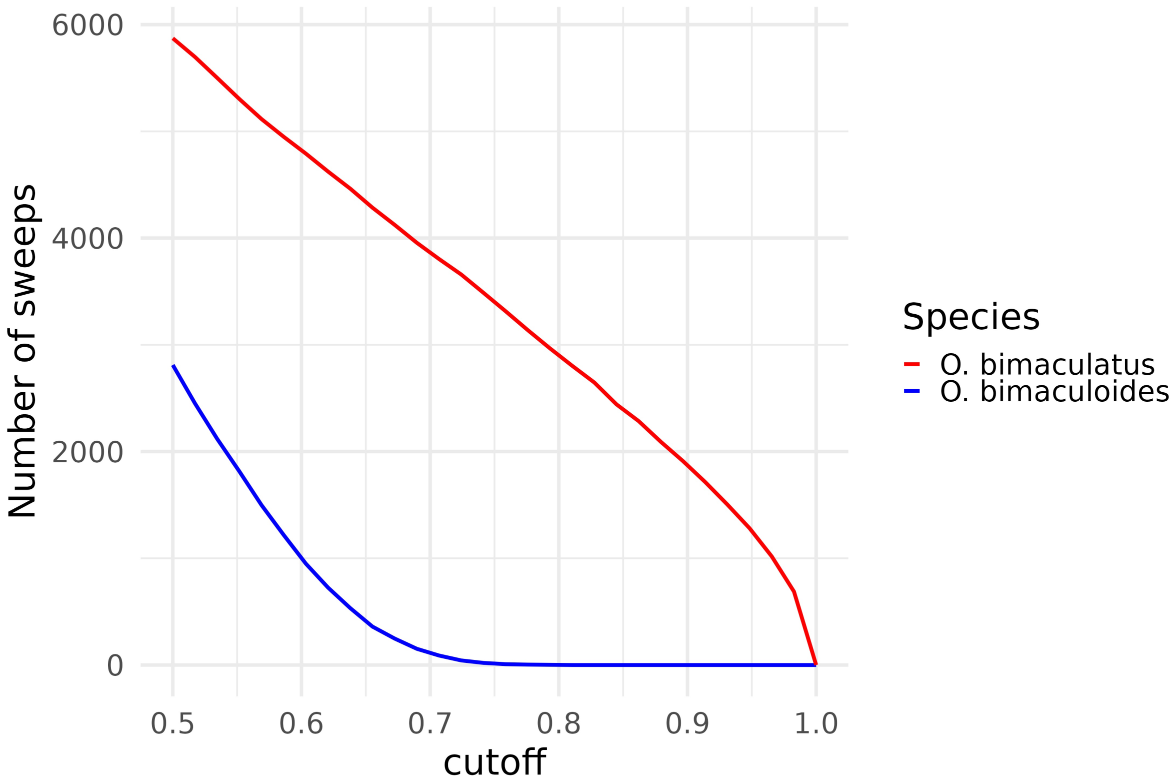


Figure S80: Total number of selective sweeps for *O. bimaculatus* (red) and *O. bimaculoides* (blue) over a range of sweep classification probability cutoffs.

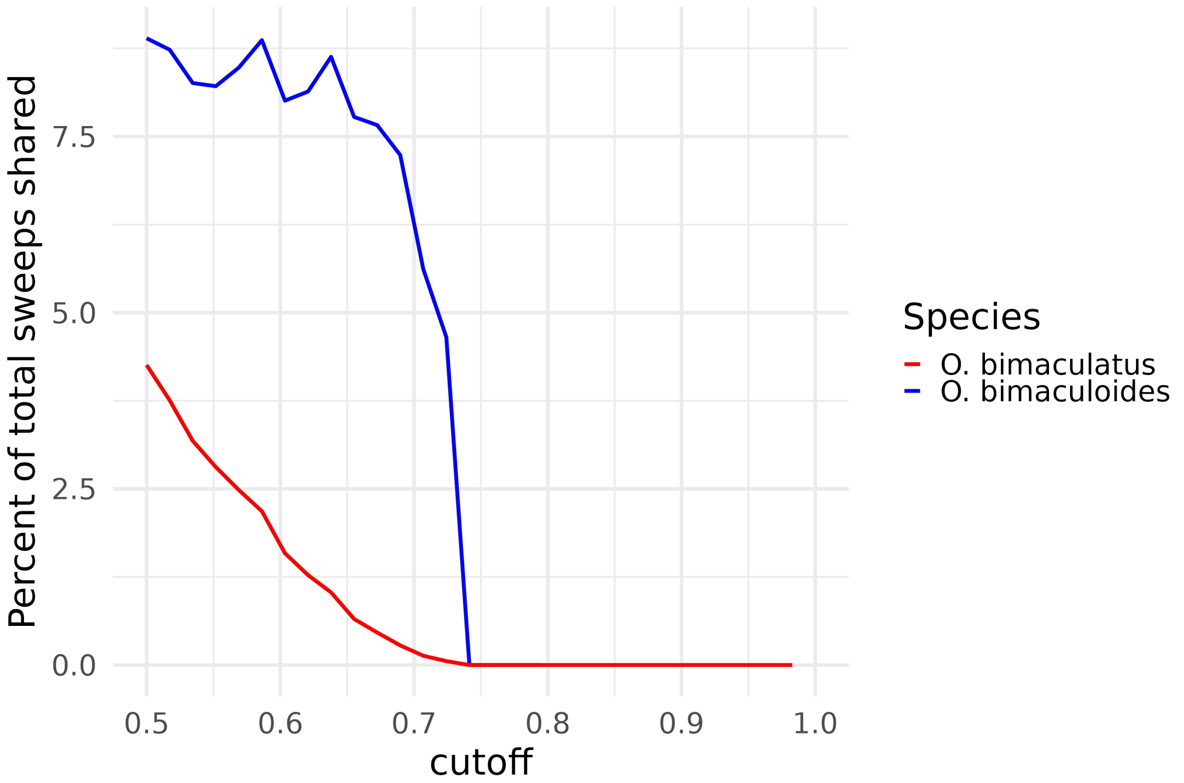


Figure S81: Percentage of selective sweeps shared between *O. bimaculatus* (red) and *O. bimaculoides* (blue) over a range of sweep classification probability cutoffs.

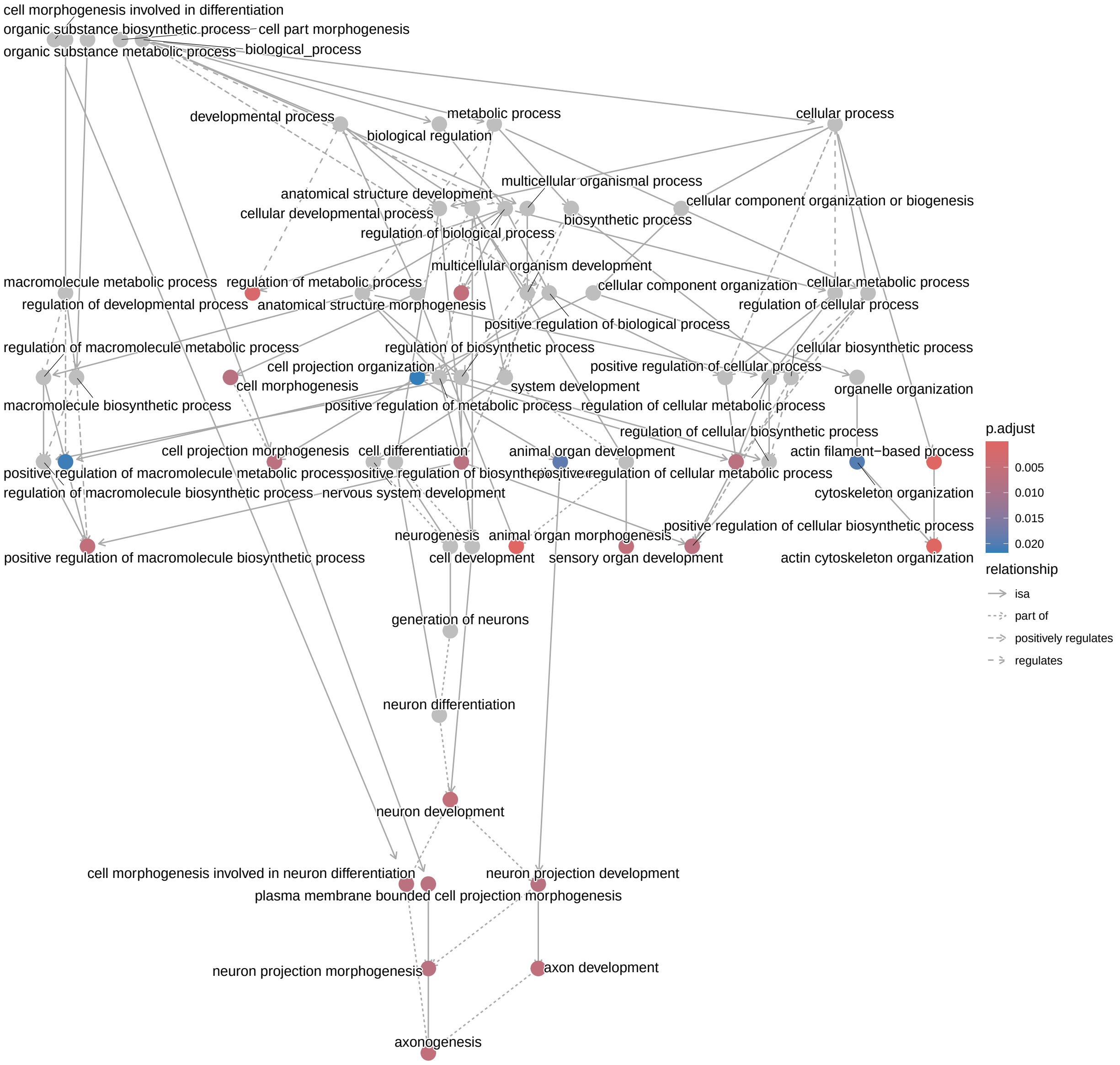


Figure S82: Significantly enriched gene ontology categories for genes overlapping with highly confident selective sweeps found in *O. bimaculatus* at a classification threshold of 0.95.

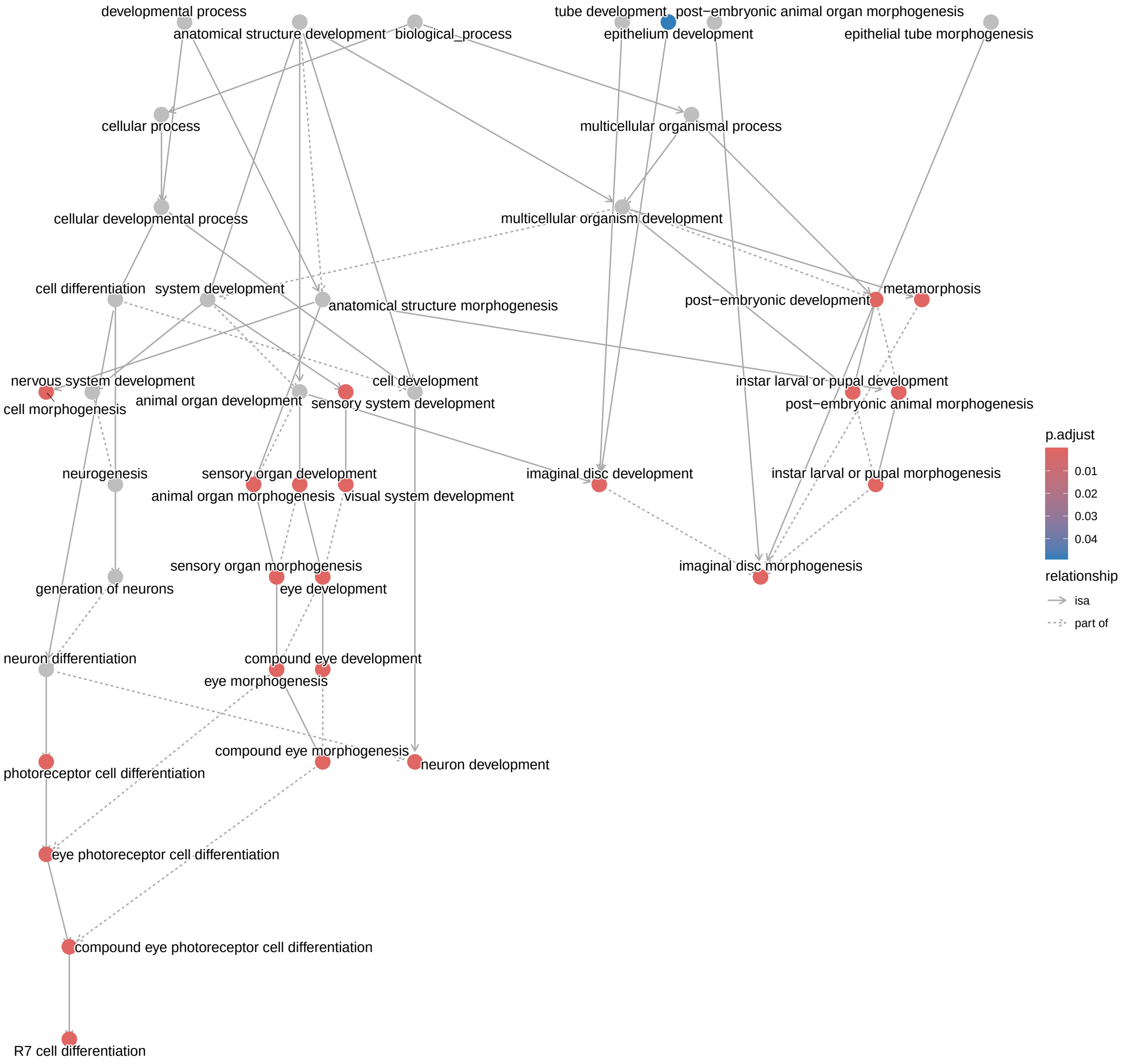


Figure S83: Significantly enriched gene ontology categories for genes overlapping with all selective sweeps shared between *O. bimaculatus* and *O. bimaculoides* at a classification threshold of 0.5.
